# A fast non-parametric test of association for multiple traits

**DOI:** 10.1101/2022.06.06.493041

**Authors:** Diego Garrido-Martín, Miquel Calvo, Ferran Reverter, Roderic Guigó

## Abstract

The increasing availability of multidimensional phenotypic data in large cohorts of genotyped individuals requires efficient methods to identify genetic effects on multiple traits. Permutational multivariate analysis of variance (PERMANOVA) offers a powerful non-parametric approach. However, it relies on permutations to assess significance, which hinders the analysis of large datasets. Here, we derive the limiting null distribution of the PERMANOVA test statistic, providing a framework for the fast computation of asymptotic *p* values. We show that the asymptotic test presents controlled type I error and high power, comparable to or higher than parametric approaches. We illustrate the applicability of our method in a number of use-cases. Using the GTEx cohort, we perform the first population-biased splicing QTL mapping study across multiple tissues. We identify thousands of genetic variants that affect alternative splicing differently depending on ethnicity, including potential disease markers. Using the UK Biobank cohort, we perform the largest GWAS to date of MRI-derived volumes of hippocampal subfields. Most of the identified loci have not been previously related to the hippocampus, but many are associated to cognition or brain disorders, thus contributing to understand the intermediate traits through which genetic variants impact complex organismal phenotypes.

## Introduction

In the past years, the availability of deep phenotype data in large cohorts of genotyped individuals has dramatically increased^1^. In addition, recent technological developments have enabled genome-wide profiling of a variety of molecular traits^2,3^. The vast majority of genome-wide association studies (GWAS) and molecular quantitative trait loci (QTL) mapping analyses test for association with genetic variants using a single trait at a time, even when multiple traits have been measured^2,4–7^. Multivariate methods, however, present several advantages over the standard univariate strategy. Many phenotypes share genetic and environmental influences, which may be reflected in their correlation structure^8^. Hence, multivariate analysis offers increased statistical power to detect genetic associations^9,10^. The multivariate setting is particularly suitable to investigate pleiotropy^11^, and can be advantageous even when only a small subset of the traits is affected by the genotype^12^. Additionally, it provides a unique framework to study the molecular mechanisms through which genetic variants act, allowing joint analyses across multiple phenotypic layers^13^. When the same trait is measured in different conditions (e.g. across tissues or environments) or over time, multivariate analyses can be used to characterize context-dependent genetic effects^14,15^. As multivariate analysis requires fewer individual tests, the multiple testing burden is also reduced.

Several methods have been proposed for multi-trait genetic association studies. MV-PLINK (canonical correlation analysis)^16^ and MultiPhen (ordinal regression)^17^ model the genotype as a dependent variable, and test for association with a weighted sum of phenotypes. However, this approach hinders the assessment of complex designs (e.g. interactions between the genotype and other covariates). In contrast, multivariate regression methods treat the phenotypes as dependent variables, offering more flexible modelling. This is the case of multivariate analysis of variance (MANOVA) or multivariate linear mixed models (LMMs)^18^. The latter have become very popular, as they can naturally account for genetic relatedness among individuals. However, fitting multivariate LMMs is computationally intensive and can be very slow in large datasets, despite continuous implementation enhancements^18,19^. A major drawback of most multivariate regression methods is the assumption that the model errors follow a multivariate normal distribution, which often does not hold (consider, for instance, multivariate proportions or count data). In practice, normalization of each individual trait, e.g. via rank-based inverse normal transformations, is commonly applied. This, however, does not guarantee multivariate normality^20^, and may reduce statistical power compared to modeling the untransformed traits^21^. Asymptotic multivariate normality is also required by approaches that leverage univariate summary statistics, such as MTAR^22^ or MOSTest^23^. In addition, estimating trait correlations from summary statistics is not straightforward, and can be affected by trait heritability or linkage disequilibrium patterns, among other factors, that need to be accounted for to avoid potential biases^22^. While several Bayesian approaches have been proposed for multi-trait association studies^12,24^, they often suffer from a large computational cost.

Altogether, this highlights the need of a fast non-parametric method suitable for multi-trait GWAS and QTL mapping. Anderson introduced a distance-based approach, known as permutational multivariate analysis of variance (PERMANOVA), that extends the univariate factorial linear model to multiple dimensions without requiring a known probability distribution of the dependent variables^25^. In PERMANOVA, the hypothesis of no-effects is tested by a permutation procedure based on a pseudo-F statistic, where the sums of squares in ANOVA are replaced by sums of inter-distances between observations. We previously employed PERMANOVA to study alternative splicing (AS) across different human populations, using the Hellinger distance between vectors of relative transcript abundances as dissimilarity metric^26^. However, while this approach remains conceptually appealing, the increased size and complexity of current datasets requires a precision for *p* value calculation that turns the permutational procedure unfeasible. Only for one-way fixed designs, Anderson and Robinson showed the asymptotic distribution of the numerator of the test statistic^27^. We used this result, implemented in sQTLseekeR, to identify genetic effects on alternative splicing across human populations^28^ and tissues^29^. Nevertheless, *p* value calculation in more complex designs still relied on permutations.

To overcome this limitation, here we obtain the asymptotic distribution of the PERMANOVA test statistic for complex designs in the Euclidean distance case. Our result also holds after any transformation of the data that preserves the independence of the observations. We develop a procedure to compute asymptotic *p* values, that we implement in the MANTA (Multivariate Asymptotic Non-parametric Test of Association) R package (available at https://github.com/dgarrimar/manta). We also provide a containerized Nextflow pipeline for multivariate GWAS using MANTA (available at https://github.com/dgarrimar/mvgwas-nf). In a typical GWAS setting (e.g. 5 traits measured in 10,000 individuals tested *vs* the genotype plus ten additional covariates), our asymptotic approach achieves over a 10^6^-fold reduction in the running time per test compared to computing 10^4^ permutations, while producing *p* values down to 10^−14^. Through a comprehensive set of simulations, we evaluate the type I error, statistical power and running time of the asymptotic test, in comparison with MANOVA and multivariate LMMs. Overall, our method presents controlled type I error and high power to detect genetic associations, comparable to the parametric approaches, and outperforming them in several settings (particularly with correlated traits when genetic effects and trait-to-trait correlations are concordant). It is also computationally more efficient than these methods, especially compared with LMMs.

We illustrate our approach in a number of real-case scenarios. First, we carry out the first population-biased splicing QTL (pb-sQTL) mapping analysis across multiple human tissues, using the Genotype-Tissue Expression (GTEx) project cohort, and identify genetic variants that affect alternative splicing differently in distinct ethnic groups. We show, in particular, the case of a pb-sQTL that is potentially a risk factor for kallikrein-5 (*KLK5*)-related diseases in European American, but not in African American individuals. Second, we perform a GWAS of the Magnetic Resonance Imaging (MRI)-derived volumes of hippocampal subfields in the UK Biobank cohort. This is the largest GWAS of hippocampal subfield volumes carried out so far. We identified 45 associated loci, dwarfing previous collections based on univariate approaches^4^. Most of these loci have not been previously related to the hippocampus. However, many of them have been associated with traits such as intellectual ability or neurodegenerative and neuropsychiatric disorders. This highlights the importance of analyzing MRI-derived endophenotypes to gain insight into the morphological alterations that mediate the impact of genetic variation in brain complex traits and diseases^30^.

## Results

### The asymptotic null distribution of the PERMANOVA test statistic

Consider a vector of *q* response variables and a vector of *p* predictor variables, both observed on *n* individuals. For example, response variables could be different brain measurements, and predictors may include the patient’s age, disease status and the genotype at a given SNP. We define **Y** as the *n* × *q* matrix of response variables, and **X** as the *n* × *p* matrix of predictors. The multivariate multiple linear regression (MMR) regresses **Y** on **X** as:

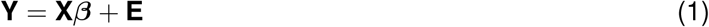

where ***β*** is a *p* × *q* matrix of parameters, and **E** a *n* × *q* matrix of random errors. Usually, model (1) includes several predictors of different types (e.g. main factors: genotype, interactions: genotype × disease status, continuous covariates: age). Anderson’s approach^25^, also known as PERMANOVA, can be used to assess the effect of these predictors by imposing conditions on a subset of parameters ***β***_0_ (i.e. ***β***_0_ = **0**). For instance, when assessing the effect of the genotype in our example above, ***β***_0_ would be the subset of parameters corresponding to the genotype term. From the general PERMANOVA pseudo-F statistic, the following expression can be derived (see Methods and Supplementary Note 1):

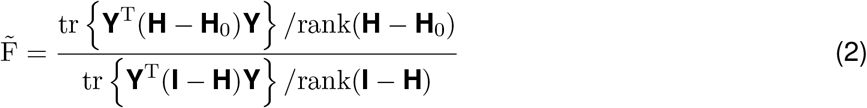

where tr denotes the trace, and **H** = **X**(**X**^T^**X**)^−1^**X**^T^ is the usual projection matrix (or *hat* matrix) in linear models. Analogously, **H**_0_ is the projection matrix corresponding to **X**_0_, which is **X** dropping the columns associated to the subset of parameters ***β***_0_ in the hypothesis.

We show that the numerator in (2) has the following limiting distribution (see Lemma 2 in Supplementary Note 1):

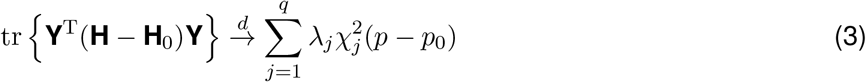

therefore the trace converges to a weighted sum of *q* independent chi-square variables with *p* − *p*_0_ = rank(**H**) − rank(**H**_0_) degrees of freedom. The coefficients *λ*_*j*_ are the eigenvalues of **Σ**, estimated in practice by the eigendecomposition of the sample covariance matrix of the residuals of the full model. The denominator in (2) converges in probability to the constant 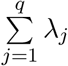 (see Supplementary Note 1).

In order to obtain the asymptotic distribution of (2), we apply some conditions to model (1). Mainly, the rows of **E** must be independent with identical *q* × *q* covariance matrix **Σ** (see Theorem 1 in Supplementary Note 1). Mutual independence of the rows of **E** is guaranteed if the observations are independently sampled. Hence, any transformation of **Y** that preserves the independence of the observations results in the same type of limiting distribution (see Supplementary Note 1). This includes, among others, the logarithm or the square root. In the specific case of proportion data, a square root transformation in (2) is equivalent to using the Hellinger distance between observations in Anderson’s pseudo-F general expression (see (9) in Supplementary Note 1). Notably, our approach does not make any assumption about the distribution model in **E**, and presents several practical advantages over standard PERMANOVA (see below).

The cumulative distribution function (CDF) of the asymptotic distribution in (3) can be used to compute *p* values for the predictors in custom MMR models. Although the CDF of a weighted sum of chi-square random variables does not have a closed form, it can be approximated with high accuracy, and several algorithms have been developed for this purpose^/^(Supplementary Figure S1). We have implemented this approach in the MANTA R package, available at https://github.com/dgarrimar/manta (see Methods).

### Comparison between asymptotic and permutational approaches

To study the properties of our asymptotic approach, we first considered a model with two categorical predictors (*A, B*) plus their interaction (*AB*). We simulated *n* = 300 observations of *q* = 3 multivariate normal response variables, with *B* under *H*_1_ (see Methods). We used our result in (3) to obtain the asymptotic null distribution of the test statistic in (2) for the interaction term (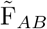), and compared it with the distribution of permuted 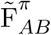 values (equivalently, we could have studied *A* instead of *AB*). Permutations were restricted to occur within the levels of factor *B* (see Methods). As shown in Figure 1a, the distribution that we derived matches exactly the one obtained by permutations, even in the upper tail region. We also provide empirical evidence that our theoretical result holds regardless of the distribution of **Y** (Supplementary Figure S2). This contrasts to the result obtained by McArtor et al.^34^, who suggested a different asymptotic distribution for the pseudo-F statistic. Their proposal departs substantially from the distribution obtained by permutations (Supplementary Figure S3), unless all the model parameters are assumed to be zero in the null hypothesis (i.e. the omnibus test).

**Figure 1.**
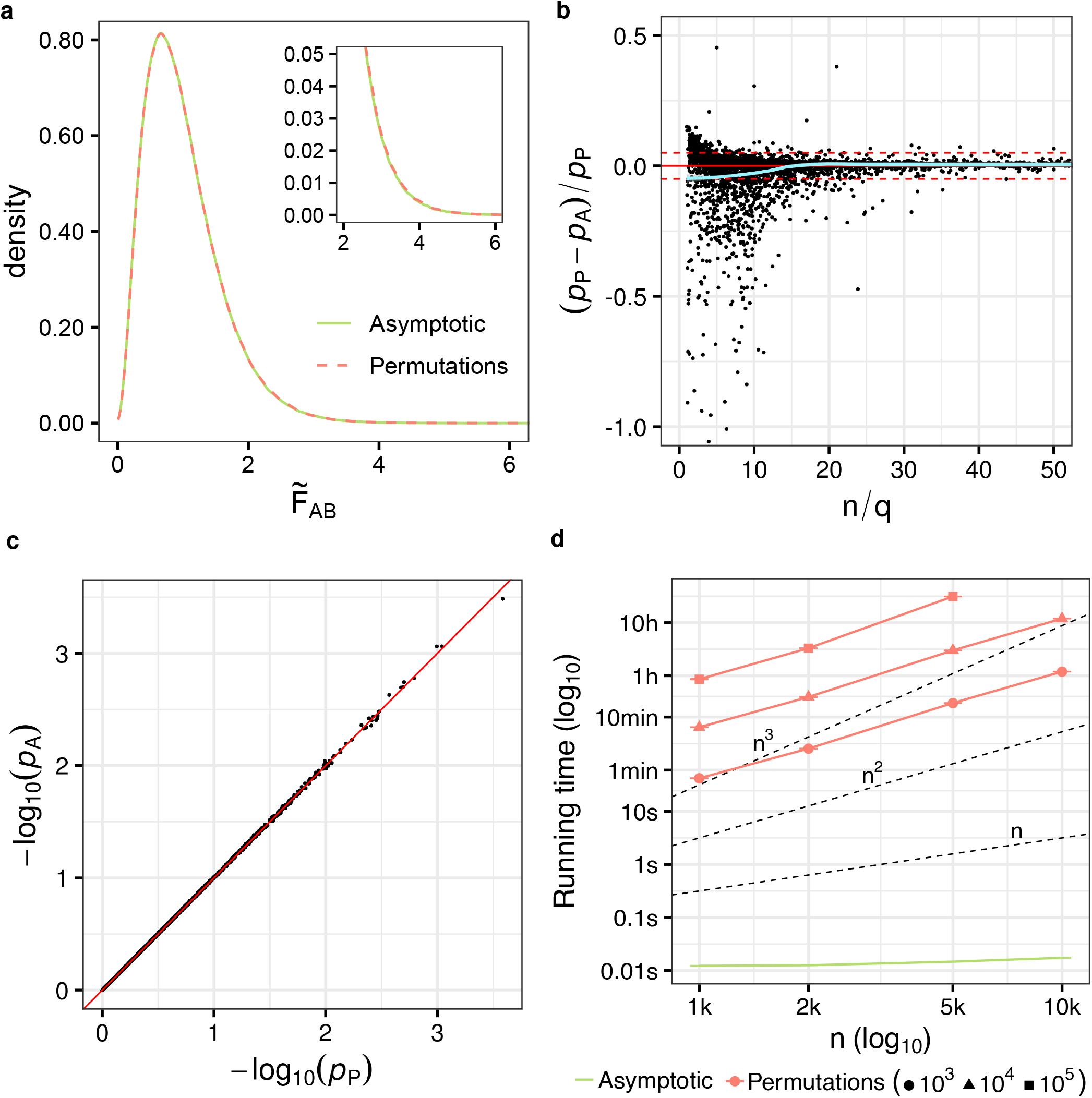
**a)** Null distribution of the PERMANOVA test statistic. Asymptotic distribution of 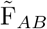 (green solid line) obtained by simulation as proposed in (3), scaled by 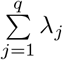, compared to the empirical distribution obtained using 10^6^ permutations (red dashed line). Simulation details: *n* = 300, *q* = 3, model (5), ***y*** ∼ N (**0, I**_*q*_) with factor *B* simulated under *H*_1_ and Δ = 1 (see Methods). The upper tail of the distribution is zoomed-in. **b)** Relative bias of asymptotic *p* values *vs n/q* ratio. Relative difference between asymptotic (*p*_*A*_) and permutation-based (*p*_*P*_, 10^5^ permutations) *p* values for the interaction term (*AB*) as a function of the ratio between the total sample size and the number of dependent variables (*n/q*). We considered values of *n* ranging from 20 to 300, and values of *q* ranging from 2 to 20. For visualization purposes, we show values of *n/q* ∈ [0,50] and relative biases ∈ [-1,0.5]. The horizontal solid red line marks the 0. The horizontal dashed red lines mark the 5% relative bias. A polynomial was fitted to the points using local fitting (LOESS), in order to describe the trend (fit in green, 95% confidence interval in grey). **c)** Comparison of asymptotic and permutation-based *p* values when the asymptotic null holds (*n* = 300, *q* = 3). **d** Empirical running time as a function of sample size (*n*) for the asymptotic and permutation-based approaches. Each point corresponds to the mean running time across 5 runs with different input data (see Methods). Error bars represent the standard error of the mean (i.e. mean ± SEM). Axes are in log_10_ scale. Dashed lines represent running time growth rates of *n, n*^2^ and *n*^3^.

Provided that our result in (3) only holds asymptotically, the accuracy of the proposed asymptotic *p* values will depend on the total sample size (*n*). In addition, other factors such as the number of response variables (*q*) or their correlation structure may also have an impact. Thus, we evaluated the relative difference between asymptotic and permutation-based *p* values across a broad range of values of *n* and *q* (Figure 1b). We considered independent response variables. Other situations with increasing degrees of correlation (in absolute value) between the response variables would be equivalent to scenarios with fewer independent response variables. When the *n/q* ratio is small, asymptotic and permutation-based *p* values may differ substantially. As the *n/q* ratio increases, this bias converges to 0. Overall, when the asymptotic null does not hold, asymptotic *p* values tend to be conservative. As a result, they can still be used without inflated type I error rates (Supplementary Figure S4). When the asymptotic null holds (note that this occurs even for relatively small values of the *n/q* ratio, i.e. *n/q* ≈ 20), we observe an almost perfect correlation between asymptotic and permutation-based *p* values (Pearson’s *r >* 0.9999), even for small *p* values (Figure 1c). Analogous results were obtained with different distributions of **Y** and data transformations (Supplementary Figure S5).

Overall, our asymptotic approach produces highly accurate *p* values, while dramatically reducing the computational cost with respect to the permutation test (Figure 1d). In addition, it allows to compute *p* values down to a precision limit of 10^−14^, difficult to achieve through permutations when the number of observations is relatively large (the smallest *p* value that can be achieved via permutation is 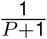, where *P* is the number of permutations). It is also particularly advantageous in complex designs where selecting the permutation schema that ensures an exact test is not trivial, or even not possible (see Methods).

### Simulation study

We designed a comprehensive set of simulations to evaluate the performance of our method in the context of phenotype-genotype association testing. We obtained genotype data from the 1000 Genomes Project (1KGP, Phase 3)^35^, and generated cohorts with different structures (unrelated individuals, population stratification, relatedness, actual 1KGP structure)^36^. We simulate phenotypes as the sum of the contribution of the effect of a genetic variant, population structure and residual noise, using an additive model. We consider a variety of scenarios, modifying the number of traits, their distribution and correlation structure, the minimum allele frequency (MAF) of the genetic variant, the structure of the population, and the fraction of phenotypic variance explained by each term in the model (see Methods). For a given scenario, we obtain 10,000 phenotype-genetic variant pairs and evaluate type I error and power of asymptotic PERMANOVA (as implemented in MANTA), and compare with those of MANOVA and multivariate LMMs (as implemented in GEMMA^18^). To ensure highly parallel, portable and reproducible simulations, we embedded our simulation framework in a containerized Nextflow^37^ computational workflow, available at https://github.com/dgarrimar/manta-sim.

### Type I error

We simulated phenotypes from a cohort of unrelated individuals using model (8) in Methods, without a causal variant effect term. We set the fraction of variance explained by the genetic background, 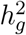, to 0.2 (see Methods). MANTA displays controlled type I error across different numbers of traits (Figure 2a). As the test does not assume any probabilistic distribution for the residuals, this does not affect type I error (Supplementary Figure S6). However, like its parametric counterparts, MANTA is sensitive to heterogeneity in multivariate dispersion^25^. We observed that type I error rates can be substantially inflated when there are differences in the trait variances between genotype groups (Figure 2c, upper panel). This is particularly problematic for lower MAFs. MANOVA and GEMMA display an analogous behaviour, with slightly larger type I errors. However, in contrast to the parametric approaches, MANTA controls well type I error when the trait correlation structure differs between genotype groups^25^ (Figure 2c, lower panel). Heterogeneity in variances or correlations may arise, for instance, in the presence of epistasis or gene by environment effects^38^. In addition, several studies have reported genetic variants associated with differences in phenotype variances (i.e. variance QTLs)^38,39^. We also simulated a scenario with strong outliers (see Methods), known to be challenging for type I error control when testing low MAF variants with methods assuming normally distributed traits^17,20^. Although outliers affect all the methods, across different numbers of traits, MANTA is more robust to its presence than MANOVA or GEMMA, specially when many traits are analyzed and trait-to-trait correlations are relatively large (Supplementary Figure S7).

**Figure 2.**
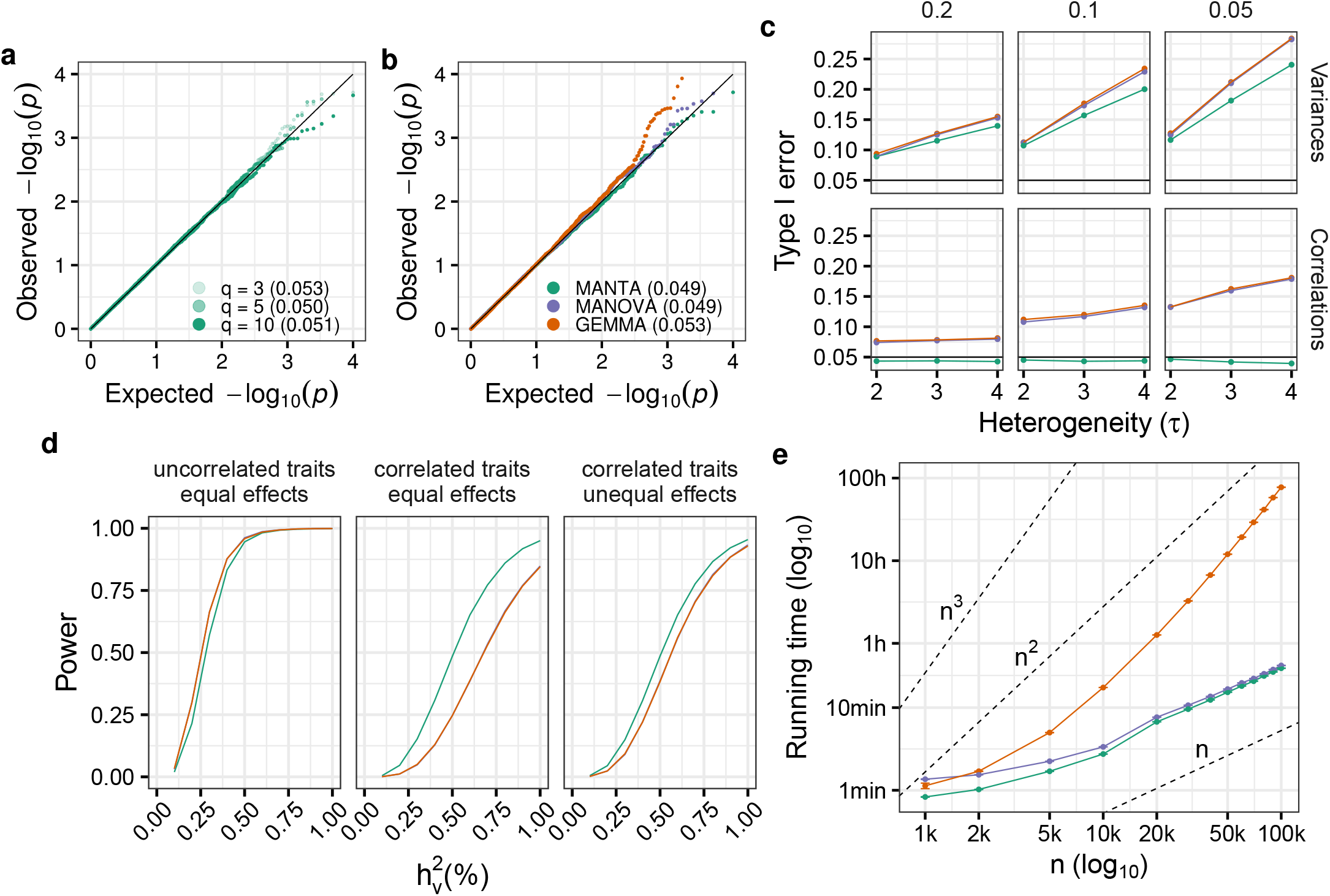
**a)** QQ-plot of *p* values obtained with MANTA for 3, 5 and 10 traits simulated in unrelated individuals (*n* = 1,000). Type I errors are shown between parentheses. Simulation details: 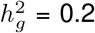, multivariate normal residuals. **b)** QQ-plot of *p* values obtained with MANTA (green), MANOVA (purple) and GEMMA (orange) on the actual 1000 Genomes Project (1KGP) genotypes (*n* = 2,504), with 5 genotype principal components (PCs) included as covariates in the model (in the case of MANTA and MANOVA). Simulation details: 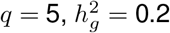, multivariate normal residuals. Colors are maintained for figures c-e. **c)** Type I error of the three methods when simulating binomial SNPs with a certain minimum allele frequency (MAF) and different degrees of heterogeneity (given by *τ*, see Methods) between genotype groups at the level of trait variances and correlations. **d)** Power of the three methods as a function of the percentage of variance explained by the causal variant 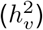, using the 1KGP genotypes, across different scenarios: the causal variant has equal effects on uncorrelated (left) and correlated (middle) traits, and unequal effects on correlated traits (right). Simulation details: actual 1KGP dataset (*n* = 2,504), *q* = 5 traits, 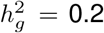, trait-to-trait correlation = 0 (left) and 0.5 (middle, right), effect size ratio = 1 (left, middle) and 2 (right), 5 genotype principal components (PCs) included as covariates in the model in the case of MANTA and MANOVA, Bonferroni corrected *p* values. **e)** Empirical running time as a function of sample size (*n*) for the three methods. Each point corresponds to the mean running time across 5 runs with different input data (see Methods). Error bars represent the standard error of the mean (i.e. mean ± SEM). Axes are in log_10_ scale. Dashed lines represent running time growth rates of *n, n*^2^ and *n*^3^.

In the context of GWAS studies, population stratification (systematic differences in ancestry) and relatedness (either family structure or cryptic relatedness among individuals with no known family relationships) are known to result in inflated type I errors^40^. When simulating phenotypes from a synthetic cohort formed by a small number of distinct populations, or from the actual 1KGP dataset (see Methods), MANTA generates calibrated *p* values, across different values of 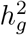, by including the top principal components (PC) of the genotype matrix as covariates in the model (Figure 2b and Supplementary Figure S8a). MANOVA behaves similarly. When simulating strongly related individuals, however, only LMM-based approaches such as GEMMA yield calibrated *p* values (Supplementary Figure S8a). Still, when the fraction of variance explained by relatedness is relatively low, our approach (MANTA+PC) performs better than parametric MANOVA+PC (Supplementary Figure S8b).

Given our previous work with multivariate proportions in alternative splicing analyses^26,28,29^, we were also interested in evaluating the performance of MANTA, MANOVA and GEMMA in this scenario. With this aim, we dropped the genetic background term in model (8), simulating trait vectors as points in the simplex (see Methods). However, while MANTA can deal directly with linearly dependent phenotypes, this is not the case of MANOVA, which requires an additional transformation. Similarly, numerical optimization in GEMMA tends to fail with this type of traits, which deviate substantially from multivariate normality (the software produces an error and does not generate any result; this is particularly striking when very few traits are analyzed, e.g. with *q* = 3, only 83 out of 10,000 tests completed correctly), or results in markedly deflated *p* values. After transforming the traits (square root), the three methods control well type I error, although GEMMA still displays slightly deflated *p* values (Supplementary Figure S9).

### Power

To evaluate power, we simulated phenotypes from the actual 1KGP dataset, given that MANTA+PC, MANOVA+PC and GEMMA displayed controlled type I error in this scenario. Figure 2d displays power curves in different settings. First, we considered uncorrelated traits with unit variance, and causal variants affecting any number of traits and explaining, on average, a fraction of the phenotypic variance 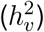 ranging from 0.1% to 1%. Overall, the three methods behave similarly across trait distributions and numbers of traits, and display large power to detect small differences (e.g. a causal variant explaining 0.5% of the variance of 1 out of 5 uncorrelated normal traits can be detected with power 0.95 after Bonferroni correction, see Supplementary Figures S10 and S11). In addition, we simulated different trait variances, correlation structures and types of effects. As previously reported, the power of multivariate methods is highly sensitive to the specific combination of genetic effects and phenotype correlations^10^. When the causal variant has similar effects across traits, as trait-to-trait correlation increases, MANTA outperforms MANOVA and GEMMA (Supplementary Figure S12). Although the scenario of concordant genetic effects and trait correlations seems more likely^10^, we also simulated different effects across traits (see Methods). In this case, MANOVA and GEMMA present higher power than MANTA with increasing trait-to-trait correlations (Supplementary Figure S13).

We also observed power differences regarding the variances of the traits, with MANTA providing higher power when the trait variances are relatively similar, and MANOVA and GEMMA displaying the opposite behaviour (Supplementary Figure S14). For each method, we barely observed differences in power when increasing the proportion of phenotypic variance explained by the genetic background at low trait-to-trait correlations. However, a drop in power was observed (more marked for MANOVA and GEMMA) when both the proportion of phenotypic variance explained by the genetic background and the trait-to-trait correlations are large (Supplementary Figure S15). In the context of multivariate proportion traits, when simulating genetic effects using a different approach (vectors of traits are points in the simplex with different locations, depending on the genotype at the causal variant and on a parameter Δ, see Methods), MANTA performs well with both untransformed and transformed (square root) traits. As stated above, MANOVA and GEMMA present problems with linearly dependent traits. Nevertheless, the three methods show similar power with transformed traits (Supplementary Figure S16).

Overall, MANTA presents high power to detect genetic effects on multiple traits, comparable to MANOVA and GEMMA, and outperforms them in several scenarios, particularly when genetic effects and trait-to-trait correlations are concordant.

### Running time and RAM usage

We evaluated the running time and RAM usage of MANTA with respect to the sample size (up to *n* = 100,000 individuals) and the number of traits analyzed (up to *q* = 32 traits), in comparison to MANOVA and GEMMA (see Methods). Results are summarized in Figure 2e and Supplementary Figure S17. The running time and RAM usage of MANTA scale approximately linearly with the number of individuals analyzed, and sublinearly with the number of responses. MANOVA displays a similar behaviour, with slightly increased running times for small sample sizes. However, GEMMA’s running time and RAM usage increase dramatically with both *n* and *q*. As a result, its application needs to be restricted to the analysis of modest numbers of phenotypes in cohorts of moderate size.

## Real data applications

### Population-biased splicing QTL mapping

Alternative splicing (AS), the process through which multiple transcript isoforms are produced from a single gene, is subject to a tight regulation, often tissue-, cell-type- or condition-specific^41^. However, while population-biased genetic effects on gene expression have been explored^2^, little is known about the population-dependent genetic control of AS across human tissues. AS can be treated as a multivariate phenotype, built from the relative abundances of a gene’s transcript isoforms^29^. Thus, here we apply asymptotic PERMANOVA to identify *cis* population-biased splicing quantitative trait loci (pb-sQTLs), across a panel of human tissues. The complete pb-sQTL catalog generated is available as Supplementary Data 1.

We obtained transcript expression data from the V8 release of the GTEx Project^2^, corresponding to up to 54 tissues from 838 deceased donors, and matched genotypes. We restricted our analyses to 818 individuals of European or African ancestry (referred to as European American (EA) and African American (AA), respectively), and to 31 tissues with more than 20 individuals of each ancestry (see Methods and Supplementary Figure S18). For *cis* pb-sQTLs mapping, we developed sQTLseekeR2.int, a slightly modified version of sQTLseekeR2, which implements asymptotic PERMANOVA to assess the significance of the effect on AS of the interaction between the donor’s genotype and ancestry (see Methods). At a 0.1 false discovery rate (FDR), we identified a total of 7,719 *cis* pb-sQTLs affecting 938 genes (i.e. pb-sGenes: 913 protein-coding genes and 25 long intergenic non-coding RNAs, lincRNAs) (Supplementary Table S1). Among them, only one (*NQO2*) is also a pb-eGene in GTEx^2^.

These numbers are substantially smaller than the ones reported for regular *cis* sQTLs in the same dataset^29^. This is explained by the more stringent pre-processing applied here (e.g. at least 10 individuals per level of the interaction are required, see Methods), which resulted in a smaller number of variant-gene pairs tested (see Supplementary Table S1), as well as by the larger statistical power required to detect significant interactions^42^. Our observation is consistent with the numbers reported by the GTEx Consortium on regular and population-biased eQTLs^2^.

As expected, the ratio between the number of genes with pb-sQTLs and the number of tested genes grows with the tissue sample size (Spearman’s *ρ* = 0.89, Supplementary Figure S19). Skin (sun-exposed) and skeletal muscle present the largest values of this ratio. Both tissues are known to have structural and functional differences between EA and AA individuals^43,44^. In addition, skin (not sun-exposed), displays a comparable –or even larger– value of this ratio with respect to other tissues with larger sample sizes (e.g. lung, thyroid, nerve). We observed that pb-sQTLs are enriched, with respect to matched non pb-sQTLs (see Methods), in functional annotations related to AS, such as splice sites or RNA-binding protein (RBP) binding sites, as well as in GWAS hits and transcription factor binding sites (Supplementary Figure S20a). pb-sQTLs are also closer to splice sites than non pb-sQTLs (two-sided Wilcoxon Rank-Sum test *p* value *<* 10^−16^, Supplementary Figure S20b). Altogether, this suggests that our pb-sQTLs are indeed capturing the biology underlying AS.

Among the pb-sQTLs identified, we found rs2739412, a pb-sQTL for the Kallikrein related peptidase 5 (*KLK5*) gene in skin (both sun-exposed and not sun-exposed). rs2739412 affects the relative abundances of the three main AS isoforms of *KLK5* in EA individuals, but not in AA individuals. In EA individuals, the abundance of isoform *KLK5-203* increases with the number of copies of the C allele, while isoforms *KLK5-201* and *KLK5-202* display the opposite behaviour (Figure 3a). The three isoforms differ in the structure of the 5’ untranslated region (5’ UTR), with *KLK5-203* having a shorter 5’ UTR than *KLK5-201* and *KLK5-203* (Figure 3b). We specifically analyzed the reads mapping in this region across genotype groups and ancestries, which confirmed our results (Figure 3c, Supplementary Table S2).

**Figure 3.**
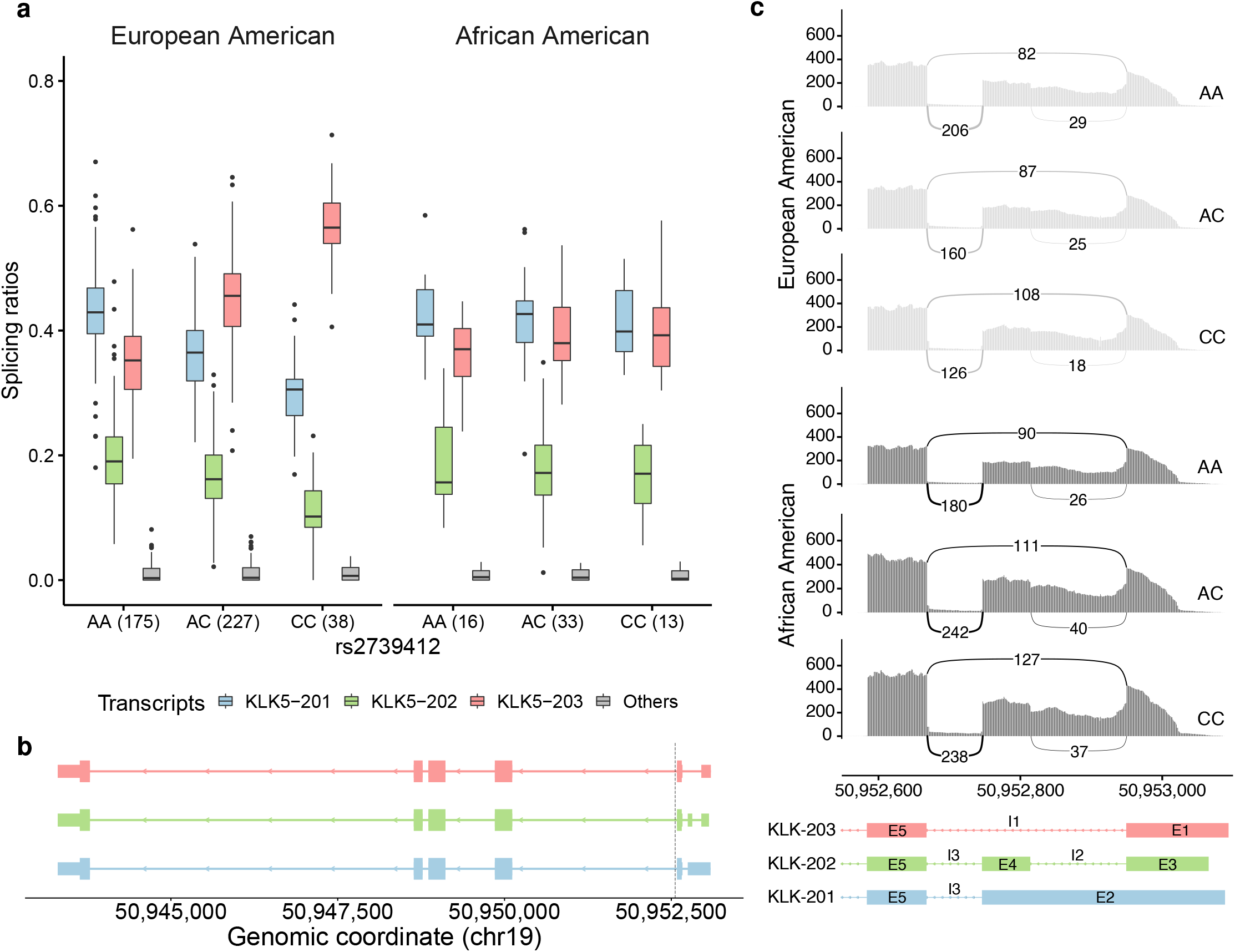
**a)** Relative abundances of the three most expressed isoforms in skin (not sun-exposed) from the gene *KLK5* (chr19:50,943,303-50,953,093, reverse strand, *KLK5-201, KLK5-202* and *KLK5-203*, all protein-coding), for each ancestry (European American, EA, African American, AA) and genotype group at the rs2739412 locus (chr19:50,952,558, A/C), represented as boxplots. In boxplots, the box represents the first to third quartiles and the median, while the whiskers indicate ± 1.5 × interquartile range. The least abundant isoforms are grouped in Others. The number of individuals in each genotype group is shown between parentheses. European American individuals that are homozygous for the reference allele (AA) at the SNP locus, present larger levels of *KLK5-201* (blue), with respect to *KLK5-202* (green) and *KLK5-203* (red). In contrast, CC homozygous express preferentially *KLK5-203* (red). Heterozygous individuals exhibit intermediate abundances. This is not observed for African American individuals. **b**) Exonic structure of the isoforms of *KLK5*, which differ in the structure of the 5’ untranslated region (5’ UTR). The dotted vertical line marks the location of the SNP. **c)** Sashimi plot corresponding to the 5’ UTR, which represents the median coverage and splice junction counts across all skin (not sun-exposed) samples of each ancestry and genotype group at rs2739412, obtained using ggsashimi^45^

KLK5 is a serine protease that plays a central role in normal stratum corneum shedding. High KLK5 activity has been related to pathological desquamation in Netherton syndrome and atopic dermatitis^46^. Given its function in extracellular matrix degradation, it has also been proposed as a candidate biomarker for epithelial-cell carcinomas^47,48^. Specifically, higher expression of *KLK5* isoforms with short 5’ UTRs, including *KLK5-203*, has been reported in ovarian cancer with respect to normal ovarian cells^47,49^. Since the 5’ UTR is an important region for post-transcriptional regulation, and its sequence is a key determinant of translation efficiency^50^, we investigated the impact of isoform usage on the levels of the KLK5 protein. Using a convolutional neural network model (https://optimus5.cs.washington.edu/MRL), trained on the results of a massively parallel reporter assay^51^, we predicted the mean ribosomal load (MRL) of the 5’ UTR of *KLK5* isoforms. We found that isoform *KLK5-203* presents almost a 60% larger predicted MRL than *KLK5-201* and *KLK5-202* (6.42 *vs* 4.10 and 3.99, respectively), suggesting that its preferential usage could lead to higher levels of the KLK5 protein.

In GTEx v8 skin tissues, rs2739412 is not identified as an eQTL for *KLK5*, but it is an sQTL for intron I3 (chr19:50,952,668-50,952,747) according to LeafCutter + FastQTL^2^, with the C allele associated to a decreased intron-excision ratio (Supplementary Figure S21). This is consistent with our results, given that this intron is present in the 5’ UTR of isoforms *KLK5-201* and *KLK5-202*, but absent from isoform *KLK5-203*. However, the LeafCutter + FastQTL pipeline did not identify other introns as significant, such as I1 (chr19:50,952,668-50,952,950), which displays clear differences between genotype groups in EA individuals (Figure 3c, Supplementary Table S2). In addition, the differences in I3 between the heterozygous and CC-homozygous individuals at rs2739412 are obscured (Supplementary Figure S21), likely by the effect of pooling EA and AA individuals, ignoring the effect of ethnicity. This highlights the increased power of our multivariate approach over univariate strategies, as well as the value of the analysis of context-dependent genetic effects on AS.

We show additional examples of pb-sQTLs identified by our approach in Supplementary Figure S22, affecting the lincRNA *SNHG8* and the protein-coding gene *RPUSD4*. Regular eQTLs and sQTLs have been identified for both genes in multiple tissues^2,29^, but no population-biased effects were reported.

### GWAS of hippocampal subfield volumes

The hippocampus is a critical structure for memory, spatial navigation and cognition^52^, which has been related to several major brain disorders, including Alzheimer’s disease^53^ and schizophrenia^54^. Multiple GWAS have identified genetic variants associated with whole hippocampal volume^55,56^. However, the hippocampus is a heterogeneous structure, with different subregions that carry out distinct functions^52^ and may be differentially affected by disease^57^. While a recent large GWAS (*n* = 21,297) reported genetic associations with individual subfield volumes^4^, this study analyzed each subfield separately, likely resulting in reduced statistical power. In contrast, here we apply asymptotic PERMANOVA (as implemented in MANTA) to identify genetic variants affecting jointly the volumes of hippocampal subfields in the UK Biobank cohort (*n* = 41,974). To the best of our knowledge, this is the largest GWAS of hippocampal subfield volumes performed to date. Summary statistics are provided as Supplementary Data 2.

Briefly, we first obtained Magnetic Resonance Imaging (MRI)-derived volumes of 12 hippocampal subfields (from anterior to posterior, approximately: parasubiculum, presubiculum, subiculum, cornu ammonis (CA) 1, 2/3 and 4, granule cell layer of the dentate gyrus (DG), hippocampus-amygdala transition area (HATA), fimbria, molecular layer of the DG, hippocampal fissure and hippocampal tail) and genotype data from the UK Biobank, corresponding to 41,974 individuals (we summed the corresponding volumes in left and right brain hemispheres, given that they are highly correlated, Supplementary Figure S23). Then, we used MANTA, embedded in a Nextflow pipeline for multivariate GWAS (available at https://github.com/dgarrimar/mvgwas-nf), to test for association between hippocampal subfield volumes and a total of 8,440,588 genetic variants (see Methods).

Our GWAS identified 45 genome-wide significant loci associated to hippocampal subfield volumes (Figure 4, Supplementary Table S4). Overlap with the GWAS Catalog revealed that, out of them, only 10 (22%) have been previously related to hippocampal volumes. 32 (71%) have been associated with other MRI-derived brain endophenotypes, and 21 (47%) with cognition, intellectual ability or neurodegenerative and neuropsychiatric disorders. 9 (20%) have not been associated to brain-related traits before. Remarkably, we replicated 8 out of 10 loci associated with individual hippocampal subfields previously identified using a univariate approach in [4]. For most of these loci, the affected subfields were identical in both studies. However, in some cases, our multivariate approach identified associations with subfields that were not captured by the univariate strategy (Supplementary Figure S24).

**Figure 4.**
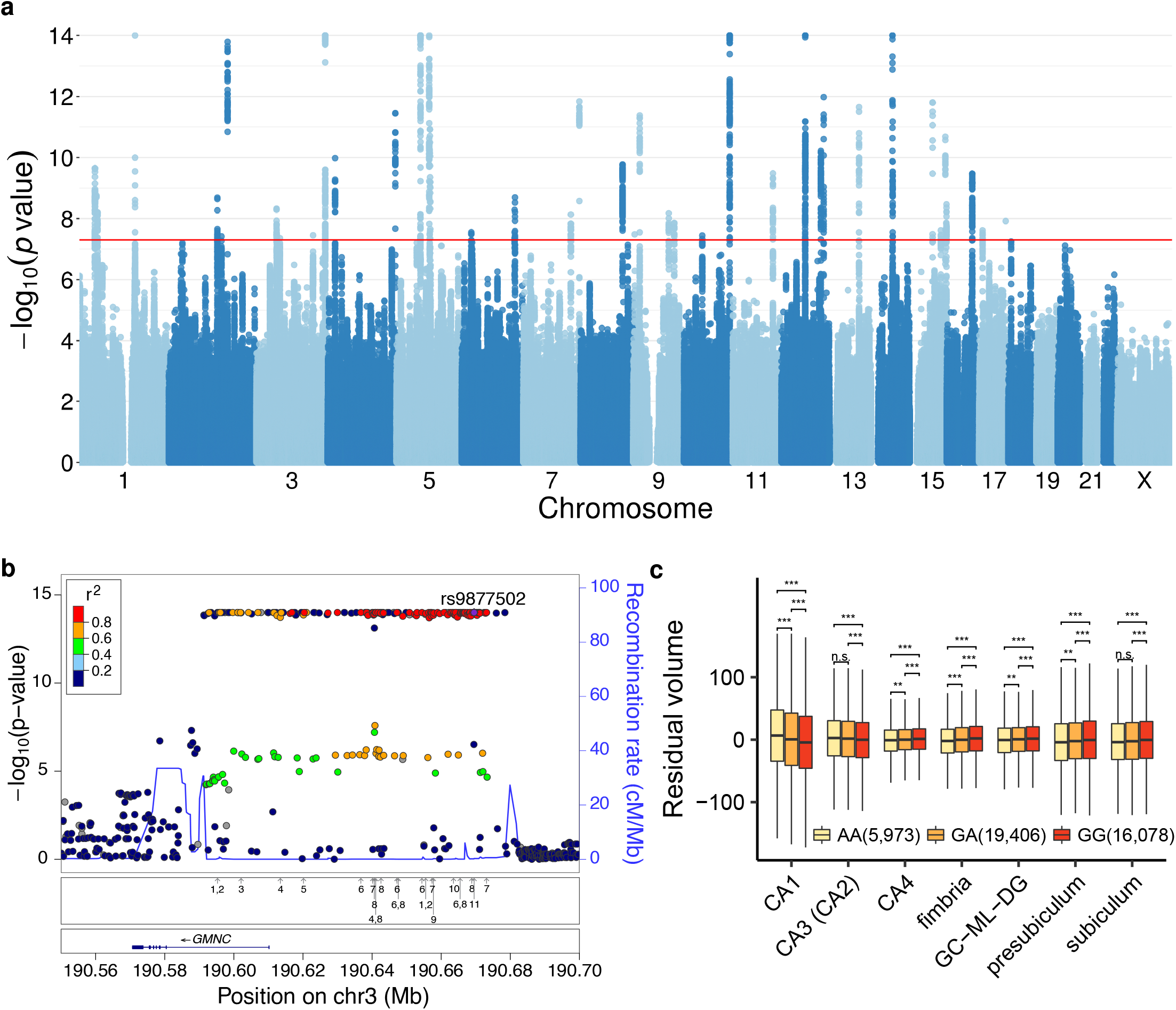
**a)** Manhattan plot showing the − log_10_ *p* value of association (asymptotic PERMANOVA test as implemented in MANTA) between genetic variants and the volumes of 12 hippocampal subfields. The horizontal red line corresponds to the genome-wide significance threshold selected (5 · 10^−8^). **b)** Regional plot of the *GMNC* locus, showing the − log_10_ *p* values for association with hippocampal subfield volumes for all tested genetic variants (asymptotic PERMANOVA test). Linkage disequilibrium patterns (color-coded) and recombination rates are also displayed. The lower panels represent the location of previous associations with brain-related traits and diseases in the GWAS Catalog (shown as arrows), encoded as follows: 1) brain morphology, 2) subcortical volume, 3) nucleus accumbens volume, 4) cortical thickness, 5) lateral ventricular volume, 6) brain region volumes, 7) cortical surface area, 8) white matter microstructure, 9) cerebrospinal P-tau181p levels, 10) cerebrospinal T-tau levels and 11) Alzheimer’s disease biomarkers. **c)** Covariate-adjusted volumes (mm^3^) of the most changing hippocampal subfields across genotype groups at rs9877502 (chr3:190,669,518, G/A) are shown as boxplots. In boxplots, the box represents the first to third quartiles and the median, while the whiskers indicate ± 1.5 × interquartile range. The number of individuals on each group is shown between parentheses. FDR adjusted *p* values (Wilcoxon Rank-Sum test) for each pairwise comparison are also displayed, encoded as follows: *** (*p* ≤ 0.001), ** (0.001 *< p* ≤ 0.01), * (0.01 *< p* ≤ 0.05), n.s. (*p >* 0.05). For visualization purposes, outliers are not shown.

Overall, the Experimental Factor Ontology (EFO) terms which correspond to the most commonly associated traits with the SNPs in the identified loci are related to brain morphology and cognition (Supplementary Figure S25). In addition, GO enrichment of genes across the 45 loci revealed functions related to neuronal development and differentiation, as well as processes such as locomotion and exploratory behaviour (Supplementary Figure S26). The former observation has also been reported in GWAS of cortical and subcortical brain measurements^23^, while the latter was recently described for genes prioritized from loci associated with the volume of the whole hippocampus and the hippocampal tail^4^. We also analyzed our results jointly with summary statistics from GWAS of Alzheimer’s disease^58^ and schizophrenia^59^, two diseases for which the hippocampus is known to be relevant^53,54^. In both cases, we observed stronger associations with the disease among the SNPs affecting hippocampal volumes (Supplementary Figure S27). This highlights the importance of analyzing multi-dimensional MRI-derived endophenotypes to gain insight into complex brain disorders^30^.

As an example of the identified loci, Figure 4b displays the chr3:190,587,740-190,836,742 locus, which contains the *GMNC* gene and its upstream region. *GMNC*, encoding the Geminin coiled-coil domain-containing protein, plays an important role in the generation of neural stem cells, which are responsible for proliferation and neurogenesis in the adult brain^60^. We show the effect of rs9877502 (chr3:190,669,518, G/A), a SNP located in this locus, across several hippocampal subfields (Figure 4c). In particular, the G allele is associated to increased CA1-3 volumes and decreased volumes of other subfields, including subiculum and presubiculum. rs9877502 has not been previously related to hippocampal volumes, but it is associated with Alzheimer’s disease risk, tangle pathology and cognitive decline^61^. Another example is the chr16:70,658,224-70,692,574 locus. The most associated SNPs overlap the *IL34* gene, which encodes interleukin 34 (IL-34). This cytokine acts as a cell type-specific ligand for the colony stimulating factor 1 receptor. In the brain, *IL34* is key for the maintenance and differentiation of the microglia^62^. Indeed, modulating IL-34 levels has been proposed as a selective approach to control microglial proliferation in neurodegenerative disorders^63^. IL-34 seems also to play a relevant role in the clearance of amyloid *β*-protein, a major hallmark of Alzheimer’s disease^64^. An intronic variant of *IL34* (rs78927322, chr16:70,636,538 C/G, not tested in our analysis due to low MAF), located 22 Kb upstream from the locus that we defined, had been previously associated to hippocampal volume in Alzheimer’s disease patients (Alzheimer Disease Neuroimaging Initiative –ADNI– cohort)^65^. Supplementary Figure S28 shows the regional plot corresponding to the *IL34* locus, as well as the effects of the lead SNP rs12923477 (chr16:70,667,804, A/G), mainly affecting CA1 volume.

In addition to the identification of genetic associations, as we used age as an independent variable in our MANTA model, we further recapitulated the well-known effect of aging^53^ across hippocampal subfields (*p* value for age 1 · 10^−14^, Supplementary Figure S29).

## Discussion

In this work, we obtain the limiting distribution of the PERMANOVA test statistic under the null hypothesis, for complex designs and Euclidean distances. Our result also holds for relatively small values of the ratio between the sample size and the number of dependent variables, and after any data transformation that preserves the independence of the observations. We provide an efficient approach to compute asymptotic *p* values for the association between any quantitative multivariate response and a set of predictors of interest, that we have implemented in the MANTA R package (available at https://github.com/dgarrimar/manta). MANTA produces highly accurate *p* values, down to a precision limit of 10^−14^, while dramatically reducing the running time with respect to the permutation test. This also avoids having to select the appropriate permutation schema, which may not be straightforward^66^. Our comprehensive simulation study in the context of genotype-phenotype association testing, and our analyses using real datasets, demonstrate that asymptotic PERMANOVA is a valuable non-parametric alternative to identify genetic effects on multiple traits in the context of GWAS and QTL mapping.

In extensive simulations, we show that the asymptotic test yields calibrated *p* values independently of the number and distribution of the traits of interest. In contrast to parametric approaches (i.e. MANOVA, multivariate LMMs), the test is robust to differences in the trait correlation structure between genotype groups, and displays smaller type I error rates in the presence of outliers and heterogeneity in trait variances. In addition, it presents large power to identify genetic associations, comparable to the parametric tests, and outperforms them in several scenarios. Particularly, our test seems more powerful when genetic effects and trait correlations are concordant. Regarding empirical running time and RAM usage as a function of the number of observations and traits analyzed, our method behaves similarly to MANOVA, and performs orders of magnitude better than multivariate LMMs. Among the three methods evaluated, only asymptotic PERMANOVA can deal with untransformed linearly dependent phenotypes, such as proportions. We also highlight the value of providing the community with a reproducible and portable simulation pipeline to evaluate multivariate methods in the context of phenotype-genotype association studies (available at https://github.com/dgarrimar/manta-sim).

To illustrate the versatility and power of asymptotic PERMANOVA, we have employed it specifically to identify genetic associations with intrinsically multivariate phenotypic traits, First, we used it to generate the first catalog of population-biased *cis* genetic effects on alternative splicing across human tissues. In particular, we identified 7,719 pb-sQTLs in the GTEx V8 cohort, mainly in tissues with known differences between individuals of European and African ancestry (i.e. skin^43^ or muscle^44^), but also in others such as thyroid or blood. As with population-biased eQTLs, identifying these associations with GTEx sample sizes is challenging^2^. However, in a scenario with highly correlated traits that depart substantially from normality (i.e. relative transcript abundances), our multivariate, non-parametric method offers increased power. Despite its modest size, the generated catalog can help to gain insights into the population-specific regulation of alternative splicing, both in health and disease contexts, as we show for the *KLK5* gene. In this case, we have identified a genetic variant that is associated with the alternative usage of *KLK5* isoforms in European Americans, but not in African Americans. Since this alternative usage may be linked to disease risk^47,49^, the responsible allele would be a risk factor only in individuals of European ancestry. This is an example of how the European bias of genetic association studies could lead to inaccurate risk assessment in non-European populations, and strongly argues for increasing the diversity of such studies, by including individuals of under-represented ancestries^67^.

Second, we employed our asymptotic approach to perform the largest GWAS of hippocampal subfield volumes to date (UK Biobank cohort, *n* = 41,974). Due to the highly shared genetic architecture of brain-related traits and disorders, this is a paradigmatic scenario where multivariate approaches are better tailored to capture underlying biology than the conventional univariate strategy^23^. Indeed, we identified 45 (35 novel) genome-wide significant loci, over four times more than the largest previous study on individual subfields^4^. Subsequent functional enrichment revealed a large overlap with loci associated with a variety of brain-related traits, gene sets with functions in neurodevelopment and links to disease (schizophrenia, Alzheimer’s disease). We also identified some candidate genes, such as *GMNC* or *IL34*, with key roles in brain pathophysiology. Overall, these results contribute to understand the mechanisms through which genetic variants impact complex organismal traits, via their effects on molecular and higher order endophenotypes, such as organ morphology.

Beyond the specific case-studies above, a wide variety of traits in Biology are intrinsically multivariate (e.g. size and connectivity of brain regions, cardiometabolic traits, facial and allometric measurements, neuropsychiatric symptoms, composition of the gut microbiota, gene networks, single-cell multi-omics, etc.) and –often– not normally distributed. Consider also the information that can be automatically extracted using deep learning techniques, such as features obtained from histological images via convolutional autoencoder networks^68^. Hence, these traits are well-suited for the analysis with asymptotic PERMANOVA. Although in this work we have focused on testing the association between genetic variants and biological traits, note that our approach, as implemented in MANTA, can be applied to any set of predictor and (quantitative) response variables of interest, expanding its usage to multiple fields (Biology, Chemistry, Economics, Epidemiology, Psychology, Physics, etc.).

Nevertheless, our method also presents some limitations. First, while it does not make any assumption on the distribution of the traits, it still requires homoscedasticity (homogeneity of trait dispersions) and independence between the observations. Both are common assumptions in most linear modelling strategies, although they can be relaxed in generalized linear models or mixed models. However, these require either defining *a priori* the variance structure, which can be particularly difficult in large and complex biological datasets, or inferring it from the actual data, which is often highly inefficient. Our assumption of independence can be violated in the presence of population stratification or genetic relatedness between individuals^40^. Although we can account for the former by including the top genotype PCs as covariates in the model, the latter can only be corrected via mixed models. Nevertheless, as we show here, multivariate LMMs can have prohibitive running times even when analyzing few traits in medium-sized cohorts.

In addition, asymptotic PERMANOVA seems more robust than MANOVA to this asumption. Recently, PERMANOVA was modified to incorporate information of genetic relatedness in a mixed model manner^69^. Evaluating whether our asymptotic result can be applied in this context would be a potential avenue for future research. A second limitation is that asymptotic PERMANOVA, as with most multivariate methods, does not provide an estimate of the effect size that can be directly employed in downstream analyses, unlike univariate approaches. Finally, our statistical framework is currently limited to Euclidean distances, which may be too restrictive for some applications.

Overall, we propose asymptotic PERMANOVA as a fast, powerful exploratory method, that can be followed up by more detailed analyses to further characterize the relationship between predictors and dependent variables. We provide an efficient, user-friendly implementation in the MANTA R package, and a containerized Nextflow pipeline for highly parallel, portable and reproducible multivariate GWAS analyses. As the size, the multi-dimensionality and the overall complexity of biological data continues to grow, our approach will enable non-parametric, multivariate analyses of millions of individuals in reasonable running times.

## Methods

### The PERMANOVA test statistic

Following the notation of the Results section, the *n* × *q* matrix of response variables, **Y** = (*y*_*ij*_) collects *n* independent observations of a vector of *q* random variables, and the *n* × *p* matrix **X**, the values of *p* predictor variables. Anderson proposed a geometric, permutation-based method (permutational multivariate analysis of variance or PERMANOVA)^25^ in order to study the effects of **X**. This approach uses a *n* × *n* suitable distance matrix between the *n* individuals based on the **Y** outcomes, allowing the computation of a pseudo-F statistic. If the Euclidean distance is used, some properties can be studied in the context of the standard multivariate multiple linear regression (MMR) (see Supplementary Note 1 for further details). The aim of MMR is to regress **Y** on **X** following the model in (1) and generalizes some of the multiple regression results (that is, when *q* = 1). For instance, the ordinary least squares (OLS) estimation of the 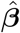 parameters is 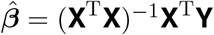, provided that **X** has full rank. 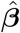 is the solution of the *q* simultaneous multiple linear regressions on each column of **Y**. Each column in 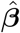 corresponds exactly to the individual multiple regression of the associated column in **Y**.

In the Euclidean distance case, if the null hypothesis of interest is ***β*** = **0** (all the parameters of every predictor are null, i.e., the omnibus test), the PERMANOVA and MMR test statistics are equivalent with expression:

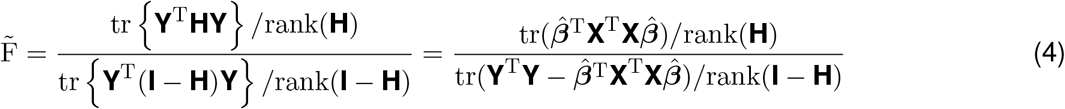

where **H** is the usual projection matrix in linear models and tr denotes the trace. Expression (2) in the Results section shows the test statistic used when testing hypotheses on a subset of parameters.

### Null distribution of the test statistic under permutation

The empirical null distribution of the PERMANOVA test statistic 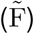 can be characterized using permutations, that is, by recomputing 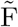after random shuffling of the data. Then, *p* values are obtained by comparing the observed value of 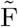 to the distribution of permuted 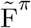 values. The only assumption of the permutation test is that the observations are exchangeable under the null hypothesis (*H*_0_). In complex designs, however, it is unclear how to ensure this in order to obtain an exact test (i.e. a test with a type I error rate exactly equal to the significance level selected *a priori*)^66^.

In the case of a model with two main factors (i.e. *A* and *B*) and an interaction term (i.e. *AB*), observations are exchangeable between the different levels of *A* and *B* only under the global null hypothesis. However, in the presence of main effects (*A* or *B* under the alternative hypothesis, *H*_1_) observations are exchangeable only within levels of other main factors. For example, if *B* is under *H*_1_, an exact permutation test for *A* that controls for the effect of *B* requires permutations to be restricted to the levels of *B*. In this scenario, unrestricted permutation of raw data yields an approximate test. See [66] for a detailed discussion. Notably, there is no exact test for the interaction term controlling for the effect of both main factors, as in this case the only possible value of the permuted test statistic is the one obtained on the original data.

### Computation of asymptotic *p* values and MANTA R package

As we describe in this work, the null distribution of the test statistic converges to a weighted sum of independent chi-square variables (see Results). To compute asymptotic *p* values, we can rely on its cumulative distribution function (CDF). Although such distribution does not have a closed form, it can be approximated with high accuracy, and several approaches are available. We focused on three of these algorithms: Imhof^31^, Davies^32^ and Farebrother^33^, as implemented in the CompQuadForm R package^70^. While the first two rely on the numerical inversion of the characteristic function, the third takes advantage of the fact that the CDF can be expressed as an infinite series of central chi-square distributions^70^.

To compare their performance, we simulated uniform sets of weights (*λ*_*j*_ ∼ *U* (*a* = 0, *b* = 1), with *j* ∈ {1, …, *q*}), considering different values of *q* and degrees of freedom for the chi-square distribution. Then, for a range of values of the test statistic we evaluated the obtained *p* values and the computation time. Note that any set of weights can be scaled to obtain values in the interval [0,1], and that scaling both the weights and the test statistic results in identical asymptotic *p* values. The typical behaviour of each algorithm is shown in Supplementary Figure S1.

Overall, we found almost identical *p* values between the three methods down to a precision of 10^−10^. However, while Farebrother *p* values decreased monotonically with the value of the test statistic, down to the precision limit (≈ 10^−14^), Imhof and Davies *p* values below 10^−10^ displayed an erratic behaviour, with values ≤ 0. In addition, regarding speed, Farebrother outperformed Imhof and Davies in the majority of scenarios. Hence, we selected the Farebrother method for *p* value calculation. Only when 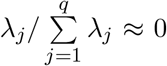 for one or more *j* in {1, …, *q*}, this approach displayed longer running times, especially for large values of the test statistic. To solve this problem, we dropped the weights for which 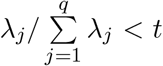. We tried several *j*=1values of *t*, and found that *t* = 10^−3^ provides a good balance between speed and accuracy.

We have implemented the asymptotic PERMANOVA test in the MANTA R package, available at https://github.com/dgarrimar/manta. MANTA enables asymptotic *p* value calculation for the predictors in user-defined MMR models, using the Farebrother method. It allows to select different types of sums of squares (i.e. I, II or III), as well as logarithm and square root data transformations.

### Simulations to compare asymptotic and permutation tests

#### Model

We considered the following MMR model, in which the response variables are regressed on two categorical predictors (i.e. factors *A* and *B*) and their interaction (*AB*):

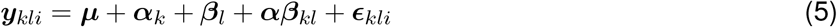

where ***y***_*kli*_ is the *q*-dimension vector corresponding to the *k, l, i* observation, ***µ*** the vector of means, ***α***_*k*_ the *q* vector of parameters (one component per response variable) associated to level *k* of factor *A* and, similarly, ***β***_*l*_ the vector of parameters of level *l* of factor *B*, ***αβ***_*kl*_ the vector corresponding to level *k, l* of the interaction *AB*, and ***ϵ***_*kli*_ the vector of random errors. We selected 2 and 3 levels for *A* and *B*, respectively, in a completely crossed, balanced design.

### Data generation

Observations of the vector of response variables (***y***, a row of **Y**) were generated by random sampling from a given multivariate distribution. We considered several distributions, varying the total sample size (*n*) and the number of response variables (*q*). We simulated both the null hypothesis (*H*_0_) of no association between predictors and responses and the alternative hypothesis (*H*_1_) of factor *B* associated with all the responses. To illustrate that transformations of **Y** that preserve the independence of the observations result in the same limiting distribution, in some scenarios we applied square root and logarithm transformations.

- **Multivariate normal**. We considered first this scenario, as it is assumed in many multivariate linear modeling approaches. Under *H*_0_, the vector of responses was simulated as ***y*** ∼ 𝒩 (**0, I**_*q*_), where **I**_*q*_ is the *q* × *q* identity matrix. Under *H*_1_, we generated ***y*** in the first and second levels of *B* with means **1** and −**1**, respectively, ensuring that *E*(***y***) = **0**.
- **Vectors of proportions**. Our interest in this scenario is related to our previous work with multivariate proportion data for the study of alternative splicing^26,28,29^, and corresponds to the generation of points in the *q* − 1 simplex. Here, ***y*** ∼ S(***p***, *σ*_*g*_), where ***p*** is a given point in the simplex and *σ*_*g*_ is the standard deviation of the generator model. We obtained ***p*** so that 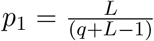 and 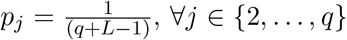 corresponds to the center of the simplex, while *L >* 1 to locations that range from the center of the simplex to one of its vertices, ***e***_**1**_ = (1, 0, …, 0). Unless stated otherwise, we set *L* = 1. To generate observations in the *q* − 1 simplex with certain variability (given by *σ*_*g*_) around ***p***, ensuring that *E*(***y***) = ***p***, we implemented an approach that performs small random displacements from ***p*** towards the simplex vertices (see Supplementary Note 2). Under *H*_1_, we generated observations of the responses in the first level of factor *B* as ***y*** ∼ *S*(***p***_Δ_, *σ*_*g*_), where ***p***_Δ_ is obtained from ***p*** advancing along the geodesic that joins ***p*** with the simplex vertex ***e***_1_ = (1, 0 …, 0). This displacement depends on a parameter Δ (see Supplementary Note 2). Analogously, in the second level of *B*, we obtained ***y*** ∼ *S*(***p***_−Δ_, *σ*_*g*_) to ensure that *E*(***y***) = ***p***. Once **Y** was obtained, we applied a square root transformation. This is equivalent to using the Hellinger distance on the untransformed data, as pointed out in the results section.
- **Gaussian copula with uniform marginals**. In this scenario, ***y*** ∼ 𝒞 (***R***), where ***R*** is the correlation matrix of ***y***. We set ***R*** to **I**_*q*_. We used the normalCopula function from the copula R package^71^, with uniform *U* (*a* = 0, *b* = 1) marginals, to generate **Y**, which was eventually centered and scaled. We also simulated observations of the responses under *H*_1_ by adding (subtracting) **1** to the rows of **Y** corresponding to the first (second) level of factor *B*.
- **Multinomial**. In this scenario, ***y*** ∼ ℳ(*N*, ***p***), where *N* and ***p*** are the number of trials and the vector of event probabilities, respectively. We set *N* to 1,000, and simulated ***p*** as in the multivariate proportion scenario (see above). Under *H*_1_, we simulated observations of the response variables in the first level of *B* as ***y*** ∼ 𝓂 (*N*, ***p***_Δ_), where ***p***_Δ_ is obtained from ***p*** as in the multivariate proportion scenario (see Supplementary Note 2). Likewise, in the second level of *B*, ***y*** ∼ 𝓂 (*N*, ***p***_−Δ_) to ensure that *E*(***y***) = *N* ***p***. Once **Y** was obtained, we applied a logarithm transformation.

### Evaluation of running time

To compare the running time of asymptotic *vs* standard PERMANOVA, we simulated multivariate normal data with increasing size (from 1,000 to 10,000 observations, *q* = 3) following model (5), under *H*_0_ (see above). We measured the running time to perform a single test, repeated 5 times with different input data. In the case of standard PERMANOVA, we considered 10^3^, 10^4^ and 10^5^ permutations. All running times were measured on a single core of an Intel Xeon Platinum 8160 CPU (2.10GHz).

### Simulations in the context of genotype-phenotype association studies

#### Model

We employed the following MMR model, in which multiple phenotypes are regressed on a set of continuous covariates, in addition to the genetic variant of interest:

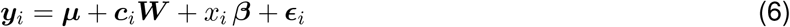

where ***y***_*i*_ is the vector of *q* traits measured in the *i*-th individual, ***µ*** the vector of intercepts, ***c***_*i*_ the vector of covariates corresponding to the *i*-th individual, ***W*** the *k* × *q* matrix of covariate effects on the *q* traits, *x*_*i*_ the genotype (scalar, i.e. 0, 1 or 2) of variant *X* in this individual, ***β*** the vector of genetic effects of *X* on each of the *q* traits, and ***ϵ***_*i*_ the vector of random errors.

We further considered a second model, without additional covariates, which incorporates a random term to account for population structure:

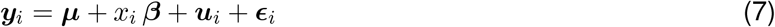

where ***u***_*i*_ is the vector of random effects due to population structure.

Here we refer to population structure, not only as large-scale, systematic differences in ancestry, but also as other forms of relatedness between individuals. Note that the former may be also accounted for by including the top *k* principal components of the genotype matrix as covariates in model (6)^40^.

#### Data generation

##### Genotypes and population structure

We obtained genotype data from the 1000 Genomes Project (1KGP, Phase 3). We considered 8,046,946 biallelic SNPs and short indels with minimum allele frequency (MAF) ≥ 0.05, measured in 2,504 individuals. In addition to the real 1KGP dataset, we simulated cohorts of *n* = 1,000 individuals with different population structures (unrelated individuals, population stratification, relatedness) as described in [36]. In short, we first assigned randomly *A* ancestors from the real 1KGP dataset to each new individual. Then, we simulated the individual’s genotype as a mosaic of blocks of 1,000 variants, each coming from one of the ancestors selected at random. Genetic relatedness in the simulated cohort depends on the number of ancestors *A* (e.g. *A* = 2 for highly related individuals, *A* = 10 for approximately unrelated individuals). Population stratification can also be simulated by sampling the ancestors of an individual from the same sub-population. To simulate unrelated and related individuals we set *A* = 10 and *A* = 2, respectively, with ancestors sampled from all European populations (CEU, FIN, GBR, IBS, TSI). To simulate population stratification, we set *A* = 10, with each individual’s ancestors sampled from the same European population.

For running time and RAM usage evaluation (see below), we simulated cohorts mimicking the 1KGP data structure (*A* = 10, each individual’s ancestors sampled from the same sub-population, considering all sub-populations available), with sample sizes in the range 1,000 − 100,000. To study the impact of heteroscedasticity and outliers on type I error as a function of MAF (see below), we simulated biallelic SNPs with a given MAF under a binomial model, with *x*_*i*_ ∼ *B*(*N, p*), where *N* = 2 and *p* = MAF. We considered MAFs ∈ {0.2, 0.1, 0.05, 0.01, 0.005}.

##### Phenotypes

We simulated different numbers (*q*) of phenotypes, as the sum of the contribution of the effect of a causal variant, population structure and residual noise, following an additive model analogous to (7):

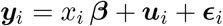

equivalently, in matrix form:

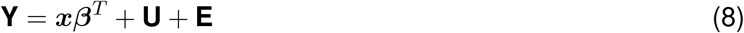

where **Y** is the *n* × *q* matrix of phenotype values, ***x*** the vector of genotypes at variant *X* for *n* individuals, ***β*** the vector of genotype effects on the *q* traits, **U** the *n* × *q* matrix of random effects due to population structure, and **E** the *n* × *q* matrix of random errors.

Random errors were simulated by random sampling from a multivariate distribution:

- **Multivariate normal**. Here ***ϵ*** ∼ 𝒩 (**0, Σ**). We simulated structured covariance matrices as follows. First, we set all pairwise trait correlations to *r*, where *r* ∈ {0, 0.2, 0.5, 0.8}. Then, we obtain 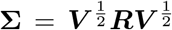, where ***R*** is the trait correlation matrix and ***V*** a *q* × *q* diagonal matrix containing the trait variances. We considered either unit variances or equally spaced variances in the range 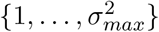, with 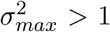 We also generated random covariance matrices as **Σ** = **AA**^*T*^, where **A** is a *q* × *q* matrix with elements *a*_*jk*_ ∼ 𝒩 (0, 1). Note that we obtained **Σ** so that it is positive definite. In this scenario, we simulated heteroscedasticity by setting covariance matrices to **Σ**, *τ/*2**Σ** and *τ* **Σ**, respectively, depending on whether the genotype at the variant is 0, 1 or 2. We further simulated differences in the correlation structure (but not in the trait variances) between genotype groups, by scaling covariances accordingly to obtain correlation matrices ***R***, 2*/τ* ***R*** and 1*/τ* ***R***, respectively, depending on whether the genotype at the variant is 0, 1 or 2. We set *τ* ∈ {2, 3, 4}.
- **Multivariate *t***. Here ***ϵ*** ∼ *t*_*ν*_(**0, Σ**). We set *ν* = 3 degrees of freedom. This distribution is similar to multivariate normal but presents very heavy tails. We simulated unit variances and different trait correlations as in the multivariate normal case.
- **Vectors of proportions**. Here ***ϵ*** ∼ 𝒮(***p***, *σ*_*g*_), where ***p*** is a given point in the simplex and *σ*_*g*_ is the standard deviation of the generator model. Details on the simulation are given in a previous section.
- **Gaussian copulas with different marginals**. Here ***ϵ*** ∼ 𝒞 (***R***), where ***R*** is the trait correlation matrix. We considered unit variances and different trait correlation structures as in the multivariate normal case. We simulated different marginal distributions: either uniform *U* (*a* = 0, *b* = 1), beta *Beta*(*α* = 0.5, *β* = 0.5) or gamma Γ(*k* = 1, *θ* = 10). Details on the simulation are given in a previous section.
- **Multinomial**. Here ***ϵ*** ∼ ℳ(*N*, ***p***), where *N* and ***p*** are the number of trials and the vector of event probabilities, respectively. Details on the simulation are given in a previous section.

Population structure effects were simulated by random sampling from a matrix normal distribution^18,36^:

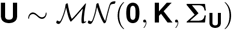

where **K** is the known *n* × *n* relatedness matrix, obtained as 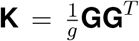 where **G** is the centered and scaled genotype matrix corresponding to *g* genome-wide variants observed in *n* individuals. **Σ**_**U**_ is the trait-to-trait covariance matrix due to population structure, obtained as **Σ**_**U**_ = **AA**^*T*^, where **A** is a *q* × *q* matrix with elements *a*_*jk*_ ∼ 𝒩 (0, 1).

We simulated the null hypothesis (*H*_0_) of no association between the genetic variant and the traits, by setting null genetic effects (***β*** = **0**, i.e. dropping the variant term in (8)). Additionally, we simulated the alternative hypothesis (*H*_1_) where a causal variant affects *t* ≤ *q* traits with equal (***β*** = **1**) or different effects (*β*_*j*_ equally spaced in the range {1, …, *β*_*max*_}, with *β*_*max*_ *>* 1). Note that the former and the latter correspond, respectively, to concordant and discordant genetic effects and trait-to-trait correlations. When generating multivariate proportion or multinomial traits, we simulated concordant genetic effects as *β*_1_ = 1, 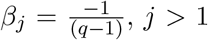. We re-scaled matrices **x*β***^*T*^, **U** and **E** so that the fractions of variance explained by the (*q−*1) genetic variant and population structure were 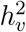 and 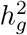, respectively. We selected 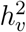 ranging from 0.001 *g* to 0.01, and 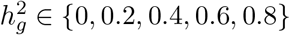.

Finally, to generate traits (rather than residuals), that are vectors of proportions (i.e. sum up to 1), we followed the steps depicted in a previous section to obtain points in the simplex. To simulate *H*_1_ (i.e. a causal variant associated to the traits), we generated values of the traits in individuals with genotypes 0, 1 and 2 at the causal variant as ***y*** ∼ *S*(***p***_−Δ_, *σ*_*g*_), ***y*** ∼ *S*(***p***, *σ*_*g*_) and ***y*** ∼ *S*(***p***_Δ_, *σ*_*g*_), respectively (see above for details). Note that in this case population structure is not taken into account.

### Evaluation of type I error and power

We selected a significance level of *α* = 0.05. For each combination of conditions, we simulated *m* = 10,000 phenotype-genotype pairs (**Y, *x***). Under *H*_0_, we evaluated the type I error for asymptotic PERMANOVA test, MANOVA (Pillai’s trace), and the multivariate LMM –implemented in GEMMA^18^– (Wald test), as follows:

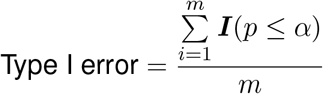

where *p* is the *p* value of the association and ***I*** the indicator function. We employed an analogous setting to estimate power when simulating phenotype-genotype pairs under *H*_1_. We evaluated power after adjusting *p* values for multiple hypothesis testing (Bonferroni).

### Evaluation of running time and RAM usage

To evaluate the running time and RAM usage of MANTA, MANOVA and GEMMA as a function of sample size, we simulated cohorts mimicking the population structure of the 1KGP dataset (see above), with increasing size (from 1,000 to 100,000 individuals). We fixed the fraction of variance explained by population structure at 0.2 and the number of traits at 3. We measured the running time to perform 10,000 tests on the same number of phenotype-genotype pairs, simulated under *H*_0_ (no causal variant effects). In the case of MANTA and MANOVA, 20 genotype principal components (PC) were included as covariates in the model. In the case of GEMMA, we also considered the time required to build the relatedness matrix. Each run was repeated 5 times with different input data. Analogously, we evaluated the running time of the three methods as a function of the number of traits (from 2 to 32), simulating cohorts with sample size 1,000. All running times were measured on a single core of an Intel Xeon Platinum 8160 CPU (2.10GHz). Regarding RAM usage, we report the peak RSS (resident set size) for the corresponding processes, estimated by Nextflow via `ps -o rss`.

### Implementation

We used the implementation of standard PERMANOVA available in the vegan R package (`adonis` method). Asymptotic *p* values were computed using our implementation of asymptotic PERMANOVA test, available at https://github.com/dgarrimar/manta (MANTA v1.0.0). MANOVA *p* values were computed using the `manova` method in the stats R package, with default parameters. We employed the implementation of the multivariate LMM available in GEMMA^18^ v0.98.3 (https://github.com/genetics-statistics/GEMMA), which relies on Wald test for *p* value calculation (default). Model (5) was assessed by MANTA with default parameters. Both MANTA (type I sums of squares, *p* values computed only for the genotype term via the option `subset`) and MANOVA assessed model (6). GEMMA assessed model (7). Numerical optimization in GEMMA failed (and consequently, the software produced an error and did not generate any result) for a small fraction of the tests performed. High error rates were found only in certain scenarios, e.g. when the trait distribution deviated substantially from multivariate normality or the number of traits analyzed was relatively large. All the simulations were performed using R v3.5.2 and Python v3.5.3. For parallelization and portability purposes, we embedded our code in a pipeline (available at https://github.com/dgarrimar/manta-sim) built using Nextflow (v20.04.1), a framework for computational workflows^37^. We also used Docker container technology (https://www.docker.com) to ensure the reproducibility of our results.

### Population-biased splicing QTL mapping

#### GTEx data and population definition

Transcript expression (transcripts per million, TPM) and variant calls were obtained from the V8 release of the GTEx Project (dbGaP accession phs000424.v8.p2 [https://www.ncbi.nlm.nih.gov/projects/gap/cgi-bin/study.cgi?study_id=phs000424.v8.p2]). These correspond to 15,253 samples from 838 deceased donors with both RNA-seq in up to 54 tissues and Whole Genome Sequencing (WGS) data available. Metadata at donor and sample level was also retrieved. GTEx V8 uses the hg38/GRCh38 human reference genome assembly and the GENCODE v26 annotation (https://www.gencodegenes.org/human/release_26.html). Further details on GTEx data preprocessing and quality control (QC) pipelines can be found in [2].

We considered individuals of European and African ancestry (97.6% of all individuals). We defined European Americans (EA, n = 715) and African Americans (AA, n = 103) as the subset of self-reported White and Black individuals, respectively (field ‘RACE’ in the GTEx metadata). Self-reported ancestry was confirmed via genotype PCA (Fig. S18). Following [2], we restricted our analysis to 31 tissues with sample sizes *>*20 in both populations.

#### pb-sQTL mapping

For *cis* population-biased sQTL (pb-sQTL) mapping, we developed a slightly modified version of sQTLseekeR2, named sQTLseekeR2.int, which implements asymptotic PERMANOVA test to assess the significance of the association between alternative splicing (AS) on one side, and the genotype, the condition of interest (i.e. ancestry), and the interaction between the two on the other. In sQTLseekeR2.int, as in sQTLseekeR2, AS is modeled as a multivariate outcome, composed by the relative abundances of the alternative transcript isoforms of a gene (splicing ratios), after a square root transformation. sQTLseekeR2.int was run using a containerized Nextflow pipeline. The software and the pipeline are available at the *interaction* branch of the https://github.com/guigolab/sQTLseekeR2 and https://github.com/guigolab/sqtlseeker2-nf repositories, respectively.

Under the assumption that most variants with *cis* effects on alternative splicing are likely to be carried on the sequence of the primary transcript or its close vicinity^29^, the *cis* window was defined as the gene body plus 5 Kb upstream and downstream the gene boundaries. We considered protein conding genes and long intergenic non-coding RNAs (lincRNAs) expressed ≥ 1 TPM in at least 80% of the samples (samples with lower gene expression were removed from the analysis of the gene), with at least two isoforms and a minimum isoform expression of 0.1 TPM (transcripts with lower expression in all samples were removed). These filters correspond to the default parameters of sQTLseekeR2.int. We analyzed only biallelic SNPs and short *indels* (autosomal + X) with MAF ≥ 0.01. We required at least 10 samples per observed level of the interaction term. Both genotype and population were treated as categorical variables. Donor ischemic time, gender and age, as well as the sample RIN (RNA integrity number) and genotyping platform were regressed out from the splicing ratios prior to association testing.

In total, 687,737 variants and 14,122 genes were analyzed. To correct for the fact that multiple variants are tested per gene, we used eigenMT^72^. eigenMT estimates the effective number of independent tests (*M*_*eff*_) per gene, considering the LD structure among the tested variants. *M*_*eff*_ is then used instead of the total number of tests (*M*) in Bonferroni correction. We set *α* = 0.05. As our test statistic is sensitive to the heterogeneity of the splicing ratios’ variability between the levels of the interaction term, a permutation-based (10^4^ permutations) multivariate homoscedasticity test^73^ was also performed for each gene-variant pair. Pairs failing this test after multiple testing correction by eigenMT were not reported as significant pb-sQTLs. eigenMT allows to compute a gene-level *p* value (corresponding to the smallest –corrected– *p* value per gene). To account for the fact that multiple genes are tested genome-wide, we applied Benjamini-Hochberg false discovery rate (FDR) to gene-level *p* values^72^. We set a FDR threshold of 0.1.

#### Functional enrichment of pb-sQTLs

We obtained eCLIP peaks in HepG2 and K562 cell lines for 113 RBPs^74^ from the ENCODE Portal (https://www.encodeproject.org, accessed 2019-07-04). For each RBP, we selected the peaks significant at FDR *<* 0.01 and with a fold-change (FC) with respect to the mock input ≥ 2. We further required a minimum overlap between replicates (50% of the length of the union of a given pair of peaks). Splice donor and acceptor sites from protein-coding and lincRNA genes were derived from the GENCODE v26 annotation (https://www.gencodegenes.org/human/release_26.html). Disease and complex-trait associated variants were retrieved from the GWAS Catalog (https://www.ebi.ac.uk/gwas, accessed 2021-01-29), extended to include GTEx variants in high linkage disequilibrium (*r*^2^ *>* 0.8) with the GWAS hits. We obtained ChIP-seq peaks for 66 transcription factors from the Ensembl Regulation dataset (ftp://ftp.ensembl.org/pub/release-86/regulation/homo_ sapiens/AnnotatedFeatures.gff.gz). The coordinates of these functional elements were overlapped with all the tested variants (either pb-sQTLs or not) to obtain a functional annotation per variant. Then, pb-sQTLs were compared to a null distribution of 1,000 sets of randomly sampled variants not identified as pb-sQTLs (i.e. non pb-sQTLs), of the same size of the pb-sQTL set. Non pb-sQTLs were matched to pb-sQTLs in terms of relative location within the gene and MAF. The enrichment was calculated as the odds ratio (OR) of the frequency of a certain annotation among pb-sQTLs to the mean frequency of the same annotation across the 1,000 non pb-sQTLs sets. The significance of each enrichment was assessed using a two-sided Fisher’s exact test.

### GWAS of the volumes of hippocampal subfields

#### UK Biobank data

We obtained Magnetic Resonance Imaging (MRI)-derived hippocampal subfield volumes, corresponding to 41,974 individuals with genotype data available, from the UK Biobank. Hippocampal sub-segmentation from T1-weighted structural images was performed with FreeSurfer v6.0 (http://surfer.nmr.mgh.harvard.edu), based on the atlas described in [75], in the framework of an image-processing pipeline developed and run on behalf of the UK Biobank^76^. We also obtained metadata at individual level. Further details on image acquisition, quality control and analysis are available at Resource 1977 (https://biobank.ctsu.ox.ac.uk/crystal/refer.cgi?id=1977). Additional information on genotyping, genotype QC and imputation can be found at Category 100319 (https://biobank.ndph.ox.ac.uk/ukb/label.cgi?id=100319) and at [77] (human reference genome assembly: hg19/GRCh37). The complete list of Data-Fields retrieved can be found in Supplementary Table S3.

#### Genome-wide association analysis

The traits of interest were the volumes of 12 hippocampal subfields from anterior to posterior, approximately: parasubiculum, presubiculum, subiculum, cornu ammonis (CA) 1, 2/3 and 4, granule cell layer of the dentate gyrus (DG), hippocampus-amygdala transition area (HATA), fimbria, molecular layer of the DG, hippocampal fissure and hippocampal tail. We obtained the total volume of each subfield by summing its corresponding volume in i) hippocampal tail and body and ii) left and right brain hemispheres. The correlation of subfield volumes between hemispheres was large (Supplementary Figure S23). We considered individual’s age, gender, total hippocampal volume (sum of all subfields minus the hippocampal fissure), and the first five genotype principal components (PCs) as relevant covariates. We analyzed only biallelic SNPs and short *indels* (autosomal + X) with MAF ≥ 0.01, missingness *<* 0.05 and at least 10 individuals per genotype group.

We used MANTA (https://github.com/dgarrimar/manta) (type I sums of squares) to test for association between genetic variants and the volumes of the 12 hippocampal subfields. We defined a model that included the covariates plus the genotype. Except for gender, all predictors were treated as continuous variables. *p* values were computed only for the genotype term via the option `subset` in MANTA. In total, 8,440,588 variants were tested for association. The analysis was run within a containerized Nextflow pipeline, available at https://github.com/dgarrimar/mvgwas-nf. We adopted the common 5 · 10^−8^ threshold for genome-wide significance.

#### Functional analysis

Loci definition was carried out by the Functional Mapping and Annotation of Genome-Wide Association Studies (FUMA) platform^78^, with default parameters. The identified loci, as well as the actual genome-wide significant variants, were overlapped with the GWAS Catalog (including the Experimental Factor Ontology (EFO) annotations for the GWAS terms, https://www.ebi.ac.uk/gwas, accessed 2021-01-29). We identified the closest genes to genome-wide significant variants, and performed hypergeometric tests to assess Gene Ontology (GO) Biological Process term over-representation. We selected as gene universe all protein-coding genes, and set a FDR threshold of 0.05. GWAS summary statistics for Alzheimer’s disease –or family history of Alzheimer’s disease–^58^ and schizophrenia^59^ were also retrieved and overlapped with our set of genome-wide significant variants, extended to include variants in high linkage disequilibrium (*r*^2^ ≥ 0.8).

## Data availability

All the data employed in this study is publicly available. GTEx data was obtained from dbGaP (https://www.ncbi.nlm.nih.gov/gap), accession phs000424.v8.p2 [https://www.ncbi.nlm.nih.gov/projects/gap/cgi-bin/study.cgi?study_id=phs000424.v8.p2]. Brain MRI and genotype data were obtained from the UK Biobank (https://www.ukbiobank.ac.uk/) under Application Number 50141. The list of Data-Fields retrieved is available in Supplementary Table S3. ENCODE eCLIP data was obtained from the ENCODE Portal (https://www.encodeproject.org). The Ensembl Regulation dataset was obtained from ftp://ftp.ensembl.org/pub/release-86/regulation/homo_sapiens/AnnotatedFeatures.gff.gz. The GWAS Catalog, including the Experimental Factor Ontology (EFO) annotations for the GWAS terms, was obtained from https://www.ebi.ac.uk/gwas. The GENCODE v26 annotation was obtained from https://www.gencodegenes.org/human/release_26.html. A detailed description of the data can be found in Methods. The pb-sQTL catalog generated and the hippocampal subfield GWAS summary statistics are provided as Supplementary Data 1 and 2, respectively (available at https://public-docs.crg.es/rguigo/Data/dgarrido/Garrido-Martin_et_al_2022.zip).

## Code availability

The MANTA R package is available at https://github.com/dgarrimar/manta. The Nextflow pipelines employed to perform the simulations are available at https://github.com/dgarrimar/manta-sim. The Nextflow pipeline for multivariate GWAS is available at https://github.com/ dgarrimar/mvgwas-nf.sQTLseekeR2-int and its companion Nextflow pipeline for condition-biased sQTL mapping are available at the interaction branch of the https://github.com/guigolab/sQTLseekeR2 and https://github.com/guigolab/sqtlseeker2-nf repositories, respectively.

## Acknowledgements

We thank Manuel Muñoz, Emilio Palumbo, Natalia Vilor and Juan D. Gispert for useful discussions. We thank Romina Garrido for administrative support. The Genotype-Tissue Expression (GTEx) project is supported by the Common Fund of the Office of the Director of the National Institutes of Health (https://commonfund.nih.gov/GTEx). This research has been conducted using the UK Biobank Resource (https://www.ukbiobank.ac.uk) under Application Number 50141. This work has been possible in part thanks to the grant CZF2019-002436 from the Chan Zuckerberg Initiative. We also acknowledge support of the Spanish Ministry of Economy, Industry and Competitiveness (MEIC) to the EMBL partnership, ‘Centro de Excelencia Severo Ochoa’, the CERCA Programme / Generalitat de Catalunya and the European Regional Development Fund (ERDF).

## Author information

### Contributions

D.G-M., M.C. and R.G. conceived and designed the study. M.C. developed the theoretical framework, with the help of D.G-M. and F.R. D.G-M. implemented the software, performed the simulations and analyzed the data. M.C. and F.R. contributed ideas and statistical advice, helping with the design of the software. D.G-M. and M.C. wrote the original draft. All the authors reviewed the final manuscript.

