## Supplementary Material for "A fast non-parametric test of association for multiple traits"

D. Garrido-Martín et al.

| Tissue | EA | AA | Variants | Genes | pb-sQTLs | pb-sGenes |
|---|---|---|---|---|---|---|
| Adipose - Subcutaneous | 492 | 71 | 325,445 | 9,697 | 1,277 | 176 |
| Adipose - Visceral (Omentum) | 402 | 51 | 162,236 | 8,618 | 197 | 30 |
| Adrenal Gland | 200 | 25 | 5,855 | 1,462 | 0 | 0 |
| Artery - Aorta | 338 | 38 | 45,281 | 5,455 | 12 | 6 |
| Artery - Coronary | 180 | 25 | 6,219 | 1,553 | 0 | 0 |
| Artery - Tibial | 489 | 76 | 345,493 | 9,464 | 1,114 | 139 |
| Breast - Mammary Tissue | 337 | 47 | 123,365 | 8,004 | 160 | 34 |
| Cells - Cultured fibroblasts | 416 | 58 | 189,829 | 8,142 | 331 | 56 |
| Cells - EBV-transformed lymphocytes | 116 | 29 | 6,753 | 1,386 | 6 | 1 |
| Colon - Sigmoid | 273 | 34 | 22,573 | 3,751 | 17 | 6 |
| Colon - Transverse | 305 | 50 | 141,357 | 8,100 | 97 | 28 |
| Esophagus - Gastroesophageal Junction | 281 | 36 | 33,299 | 4,538 | 10 | 4 |
| Esophagus - Mucosa | 424 | 60 | 223,860 | 9,237 | 473 | 69 |
| Esophagus - Muscularis | 396 | 55 | 206,883 | 9,056 | 195 | 32 |
| Heart - Atrial Appendage | 322 | 42 | 70,406 | 6,333 | 19 | 6 |
| Heart - Left Ventricle | 334 | 42 | 53,526 | 5,338 | 36 | 2 |
| Lung | 444 | 58 | 241,674 | 9,871 | 385 | 47 |
| Minor Salivary Gland | 118 | 23 | 4,224 | 1,129 | 5 | 3 |
| Muscle - Skeletal | 602 | 86 | 327,757 | 8,439 | 1,499 | 199 |
| Nerve - Tibial | 449 | 67 | 329,639 | 10,187 | 619 | 108 |
| Ovary | 140 | 21 | 2,298 | 730 | 0 | 0 |
| Pancreas | 252 | 39 | 40,302 | 4,908 | 5 | 4 |
| Prostate | 186 | 27 | 13,958 | 2,171 | 57 | 3 |
| Skin - Not Sun Exposed (Suprapubic) | 440 | 62 | 249,241 | 9,648 | 602 | 100 |
| Skin - Sun Exposed (Lower leg) | 518 | 73 | 333,619 | 10,090 | 1,938 | 270 |
| Small Intestine - Terminal Ileum | 144 | 25 | 5,919 | 1,509 | 0 | 0 |
| Spleen | 185 | 34 | 20,486 | 3,638 | 6 | 3 |
| Stomach | 269 | 44 | 80,945 | 6,776 | 14 | 7 |
| Testis | 277 | 34 | 40,283 | 5,036 | 5 | 5 |
| Thyroid | 494 | 62 | 284,363 | 10,085 | 616 | 105 |
| Whole Blood | 574 | 80 | 155,988 | 5,664 | 555 | 60 |
| | | | | **Total (unique)** | 7,719 | 938 |

**Table S1.** Number of individuals of each ancestry (European American, EA, and African American, AA), variants and genes tested; pb-sQTLs and pb-sGenes (genes with at least one pb-sQTL) identified across tissues after multiple testing correction (see Methods).

| | European American | | | African American | | |
|---|---|---|---|---|---|---|
| | **AA** | **AC** | **CC** | **AA** | **AC** | **CC** |
| I1 in *KLK5-203* (chr19:50,952,668-50,952,950) | 82 (0.26) | 87 (0.32) | 108 (0.42) | 90 (0.31) | 111 (0.28) | 127 (0.32) |
| I2 in *KLK5-202* (chr19:50,952,814-50,952,950) | 29 (0.09) | 25 (0.09) | 24 (0.09) | 26 (0.09) | 40 (0.10) | 37 (0.09) |
| I3 in *KLK5-201* and *KLK5-202* (chr19:50,952,668-50,952,747 ) | 206 (0.65) | 160 (0.59) | 126 (0.49) | 180 (0.61) | 242 (0.62) | 238 (0.59) |

**Table S2.** Median number (proportion) of reads supporting three alternative *KLK5* 5' UTR introns, across individuals with different ancestries (European American, African American) and genotypes at rs2739412 (chr19:50,952,558, A/C)

| Data-Field | Description |
|---|---|
| 31 | Sex |
| 21022 | Age at recruitment |
| 22828 | Imputation from genotype (WTCHG) |
| 26620 | Volume of Hippocampal-tail (left hemisphere) |
| 26621 | Volume of subiculum-body (left hemisphere) |
| 26622 | Volume of CA1-body (left hemisphere) |
| 26623 | Volume of subiculum-head (left hemisphere) |
| 26624 | Volume of hippocampal-fissure (left hemisphere) |
| 26625 | Volume of presubiculum-head (left hemisphere) |
| 26626 | Volume of CA1-head (left hemisphere) |
| 26627 | Volume of presubiculum-body (left hemisphere) |
| 26628 | Volume of parasubiculum (left hemisphere) |
| 26629 | Volume of molecular-layer-HP-head (left hemisphere) |
| 26630 | Volume of molecular-layer-HP-body (left hemisphere) |
| 26631 | Volume of GC-ML-DG-head (left hemisphere) |
| 26632 | Volume of CA3-body (left hemisphere) |
| 26633 | Volume of GC-ML-DG-body (left hemisphere) |
| 26634 | Volume of CA4-head (left hemisphere) |
| 26635 | Volume of CA4-body (left hemisphere) |
| 26636 | Volume of fimbria (left hemisphere) |
| 26637 | Volume of CA3-head (left hemisphere) |
| 26638 | Volume of HATA (left hemisphere) |
| 26639 | Volume of Whole-hippocampal-body (left hemisphere) |
| 26640 | Volume of Whole-hippocampal-head (left hemisphere) |
| 26641 | Volume of Whole-hippocampus (left hemisphere) |
| 26642 | Volume of Hippocampal-tail (right hemisphere) |
| 26643 | Volume of subiculum-body (right hemisphere) |
| 26644 | Volume of CA1-body (right hemisphere) |
| 26645 | Volume of subiculum-head (right hemisphere) |
| 26646 | Volume of hippocampal-fissure (right hemisphere) |
| 26647 | Volume of presubiculum-head (right hemisphere) |
| 26648 | Volume of CA1-head (right hemisphere) |
| 26649 | Volume of presubiculum-body (right hemisphere) |
| 26650 | Volume of parasubiculum (right hemisphere) |
| 26651 | Volume of molecular-layer-HP-head (right hemisphere) |
| 26652 | Volume of molecular-layer-HP-body (right hemisphere) |
| 26653 | Volume of GC-ML-DG-head (right hemisphere) |
| 26654 | Volume of CA3-body (right hemisphere) |
| 26655 | Volume of GC-ML-DG-body (right hemisphere) |
| 26656 | Volume of CA4-head (right hemisphere) |
| 26657 | Volume of CA4-body (right hemisphere) |
| 26658 | Volume of fimbria (right hemisphere) |
| 26659 | Volume of CA3-head (right hemisphere) |
| 26660 | Volume of HATA (right hemisphere) |
| 26661 | Volume of Whole-hippocampal-body (right hemisphere) |
| 26662 | Volume of Whole-hippocampal-head (right hemisphere) |
| 26663 | Volume of Whole-hippocampus (right hemisphere) |

**Table S3.** List of Data-Fields retrieved from the UK Biobank. Further details on each Data-Field are available at the UK Biobank data showcase (`https://biobank.ndph.ox.ac.uk/showcase`).

| Locus | chr | start | end | Lead SNP | Position | *p* value |
|---:|---:|---:|---:|---|---:|---|
| 1 | 1 | 43,760,236 | 43,949,718 | rs1004291 | 43,858,630 | $2.23 \cdot 10^{-10}$ |
| 2 | 1 | 47,974,123 | 47,980,916 | rs6658111 | 47,980,916 | $1.77 \cdot 10^{-9}$ |
| 3 | 1 | 50,828,316 | 51,480,258 | rs11584472 | 51,114,503 | $6.24 \cdot 10^{-9}$ |
| 4 | 1 | 155,033,308 | 155,087,188 | rs11589479 | 155,033,308 | $1.00 \cdot 10^{-14}$ |
| 5 | 2 | 134,320,098 | 134,533,762 | rs3936142 | 134,471,302 | $2.07 \cdot 10^{-9}$ |
| 6 | 2 | 145,755,449 | 145,799,709 | rs7589308 | 145,761,254 | $3.85 \cdot 10^{-8}$ |
| 7 | 2 | 162,796,517 | 162,891,848 | rs2909455 | 162,843,078 | $1.62 \cdot 10^{-14}$ |
| 8 | 3 | 55,535,699 | 55,560,144 | rs73077403 | 55,558,889 | $4.73 \cdot 10^{-9}$ |
| 9 | 3 | 55,943,930 | 56,273,743 | rs17825124 | 56,052,060 | $1.57 \cdot 10^{-8}$ |
| 10 | 3 | 64,554,254 | 64,626,922 | rs11923848 | 64,602,886 | $4.34 \cdot 10^{-8}$ |
| 11 | 3 | 156,715,450 | 156,789,474 | rs11719102 | 156,786,544 | $3.54 \cdot 10^{-8}$ |
| 12 | 3 | 190,587,740 | 190,836,742 | rs61500084 | 190,636,749 | $1.00 \cdot 10^{-14}$ |
| 13 | 4 | 20,088,468 | 20,239,195 | rs13150272 | 20,121,121 | $1.04 \cdot 10^{-10}$ |
| 14 | 4 | 187,366,735 | 187,390,722 | 4:187389308_TA_T | 187,389,308 | $3.53 \cdot 10^{-12}$ |
| 15 | 5 | 65,923,428 | 66,219,202 | rs258260 | 66,076,217 | $1.00 \cdot 10^{-14}$ |
| 16 | 5 | 90,796,839 | 91,040,975 | rs7724031 | 90,808,954 | $1.00 \cdot 10^{-14}$ |
| 17 | 5 | 92,182,752 | 92,361,481 | rs888814 | 92,186,429 | $5.50 \cdot 10^{-11}$ |
| 18 | 5 | 92,683,607 | 93,548,292 | 5:92797041_TG_T | 92,797,041 | $1.19 \cdot 10^{-11}$ |
| 19 | 6 | 25,786,226 | 27,730,334 | rs34073492 | 26,739,452 | $2.80 \cdot 10^{-8}$ |
| 20 | 6 | 146,976,037 | 147,008,552 | rs9497602 | 146,990,571 | $5.17 \cdot 10^{-9}$ |
| 21 | 6 | 148,041,598 | 148,095,291 | rs9399619 | 148,056,480 | $2.03 \cdot 10^{-9}$ |
| 22 | 7 | 132,045,957 | 132,141,199 | rs7803579 | 132,087,061 | $7.42 \cdot 10^{-9}$ |
| 23 | 7 | 155,797,978 | 155,808,233 | rs56354640 | 155,799,049 | $1.46 \cdot 10^{-12}$ |
| 24 | 8 | 116,464,988 | 116,636,719 | rs2049867 | 116,588,570 | $1.68 \cdot 10^{-10}$ |
| 25 | 9 | 3,924,447 | 3,944,778 | rs7861718 | 3,924,447 | $3.29 \cdot 10^{-8}$ |
| 26 | 9 | 18,006,202 | 18,109,156 | rs12337268 | 18,086,410 | $4.16 \cdot 10^{-12}$ |
| 27 | 9 | 98,216,876 | 98,280,656 | rs28536201 | 98,276,753 | $6.68 \cdot 10^{-9}$ |
| 28 | 9 | 113,651,047 | 113,679,463 | rs4246883 | 113,674,955 | $1.38 \cdot 10^{-8}$ |
| 29 | 10 | 50,210,710 | 50,384,525 | 10:50252571_GA_G | 50,252,571 | $3.60 \cdot 10^{-8}$ |
| 30 | 10 | 126,302,936 | 126,564,695 | rs7099316 | 126,469,005 | $1.00 \cdot 10^{-14}$ |
| 31 | 11 | 110,945,939 | 111,083,785 | rs58066679 | 111,056,158 | $3.32 \cdot 10^{-10}$ |
| 32 | 12 | 65,372,996 | 66,065,136 | rs61921502 | 65,832,468 | $1.00 \cdot 10^{-14}$ |
| 33 | 12 | 109,029,333 | 109,260,895 | rs7484851 | 109,134,193 | $6.07 \cdot 10^{-11}$ |
| 34 | 12 | 117,309,440 | 117,338,504 | rs146607495 | 117,319,202 | $1.06 \cdot 10^{-12}$ |
| 35 | 13 | 81,164,609 | 81,468,042 | rs9545540 | 81,298,245 | $2.23 \cdot 10^{-12}$ |
| 36 | 14 | 59,064,730 | 59,092,581 | rs160459 | 59,074,136 | $1.00 \cdot 10^{-14}$ |
| 37 | 14 | 59,629,611 | 59,804,999 | rs1252992 | 59,665,286 | $2.01 \cdot 10^{-12}$ |
| 38 | 15 | 63,274,450 | 63,297,719 | rs1489315 | 63,292,446 | $1.57 \cdot 10^{-12}$ |
| 39 | 15 | 86,004,091 | 86,079,115 | rs1531854 | 86,022,084 | $2.44 \cdot 10^{-8}$ |
| 40 | 15 | 98,429,818 | 98,446,241 | rs5814830 | 98,431,818 | $2.07 \cdot 10^{-11}$ |
| 41 | 15 | 101,200,873 | 101,210,973 | rs8036545 | 101,210,641 | $1.38 \cdot 10^{-8}$ |
| 42 | 15 | 101,713,433 | 101,794,473 | 15:101778449_ACCAT_A | 101,778,449 | $3.43 \cdot 10^{-9}$ |
| 43 | 16 | 70,658,224 | 70,692,574 | rs12923477 | 70,667,804 | $3.37 \cdot 10^{-10}$ |
| 44 | 17 | 9,137,688 | 9,155,926 | rs80100171 | 9,151,754 | $2.44 \cdot 10^{-8}$ |
| 45 | 17 | 73,439,428 | 73,532,213 | rs77001351 | 73,498,652 | $1.21 \cdot 10^{-8}$ |

**Table S4.** Whole-genome significant loci for hippocampal subfield volumes identified by MANTA. Locus definition was performed by the FUMA platform (default parameters). Coordinates correspond to the hg19/GRCh37 human reference genome assembly.

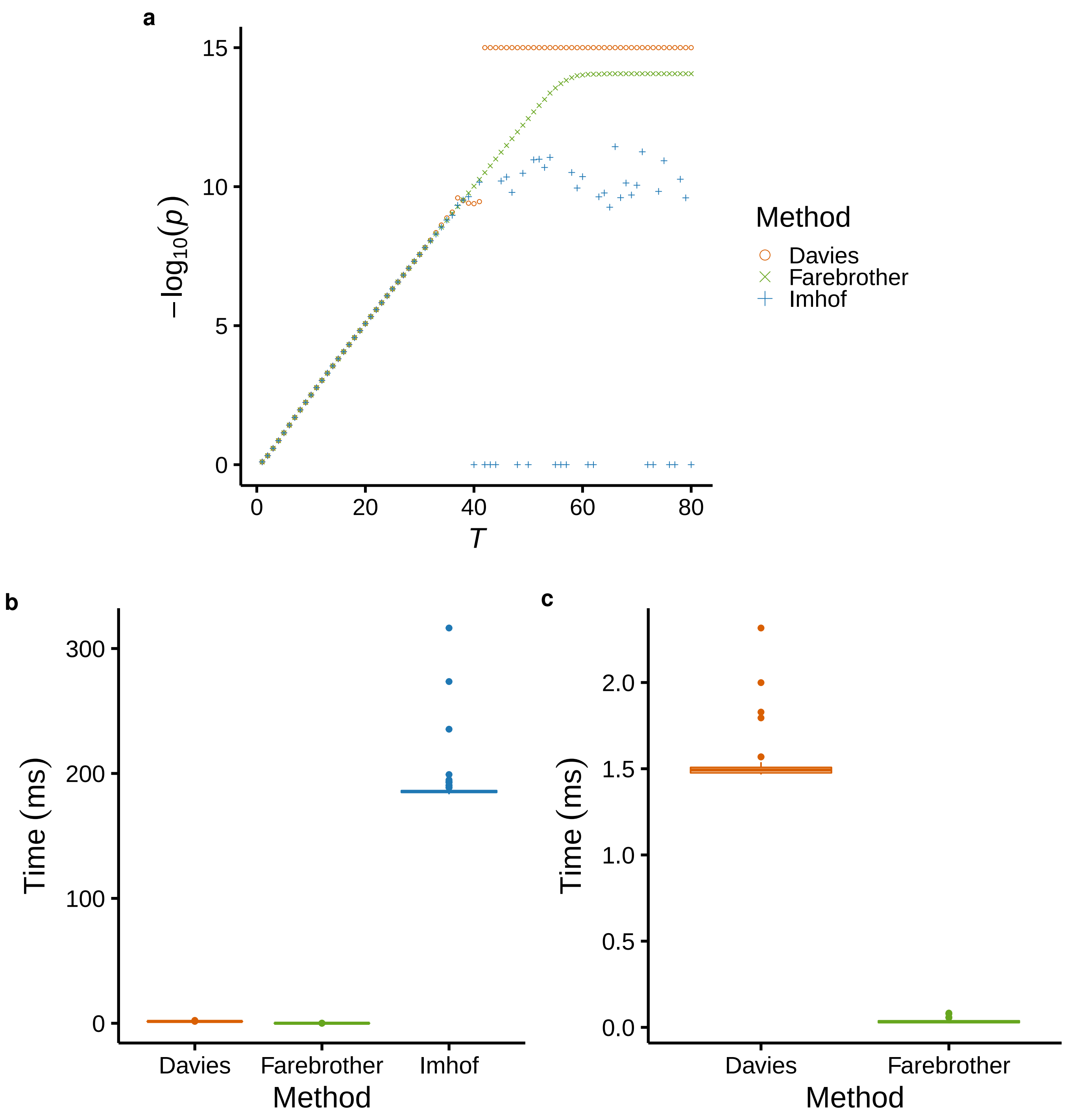


**Figure S1.** **a)** Behaviour of the *p* values obtained by Davies, Farebrother and Imhof algorithms, as implemented in the CompQuadForm R package[70], as a function of different values of the test statistic (labelled as $T$, x-axis). The data shown correspond to a simulation in which $q = 5$ weights are sampled from a uniform distribution ($\lambda_j \sim U(a = 0, b = 1)$, $j \in \{1, \ldots, 5\}$), $T \in \{1, 2, \ldots, 80\}$, and chi-square variables have 1 degree of freedom. While Farebrother *p* values decrease monotonically with the value of $T$, down to the precision limit ($\approx 10^{-14}$), Imhof and Davies generated *p* values of 0 (here displayed as $p = 10^{-15}$) or even negative (here displayed as $p = 1$). **b)** Running time of the three methods (in milliseconds), computed across 100 runs with $T = 30$ using the microbenchmark R package[79]. **c)** Zoom of b).

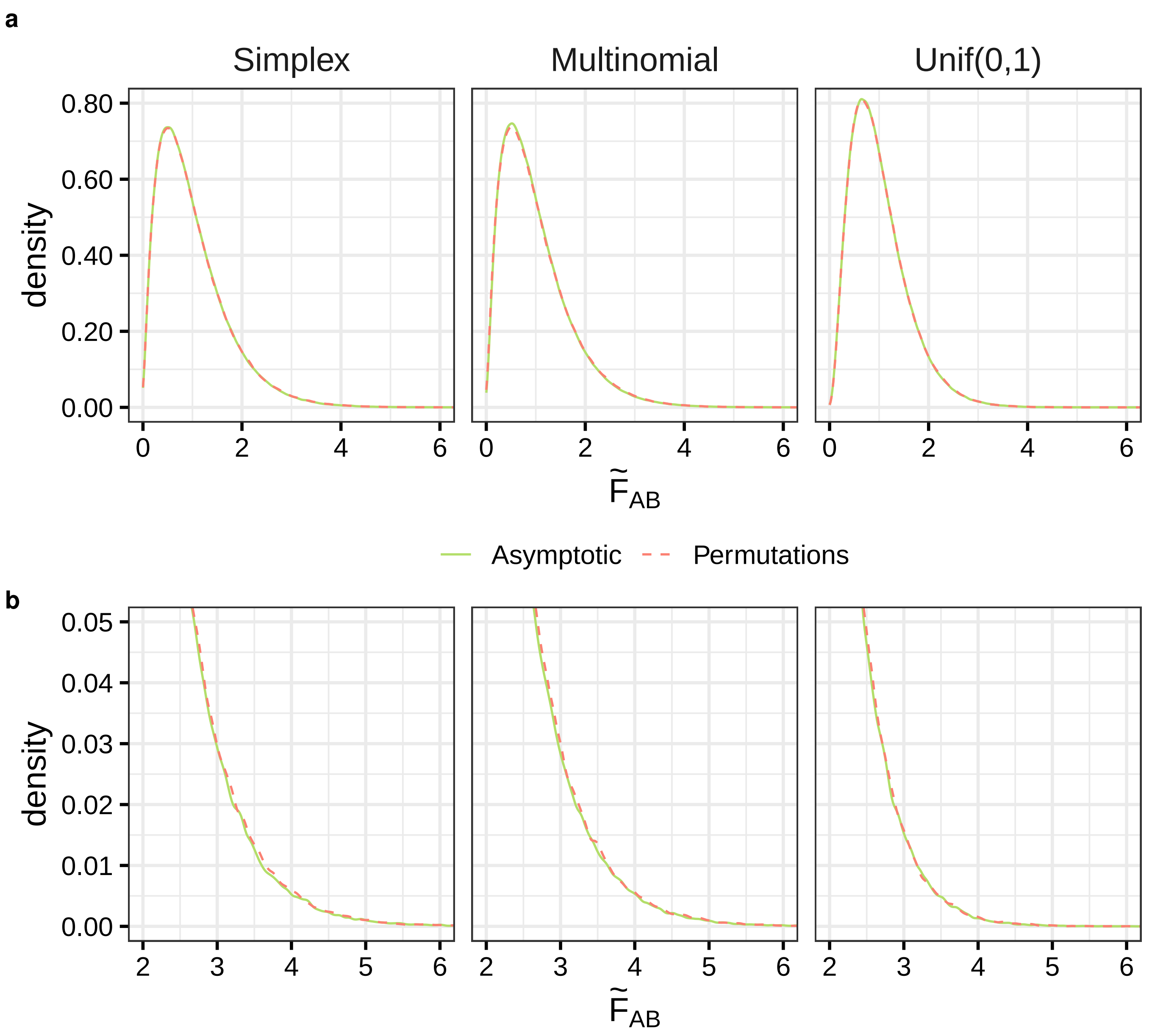


**Figure S2. a)** Asymptotic null distribution of the PERMANOVA test statistic for the interaction term (green solid line), compared to the empirical null distribution obtained using $10^6$ permutations (red dashed line), for different distributions of the response variables: multivariate proportions (left), multinomial (middle), and Gaussian copula with uniform marginals (right). **b)** Zoom of the upper tail of the distribution. Simulation details: model (5), $n = 300$ observations of $q = 3$ response variables, with $B$ generated under $H_1$, $\Delta$ set to 0.1 (simplex, multinomial) and 1 (Gaussian copula) (see Methods).

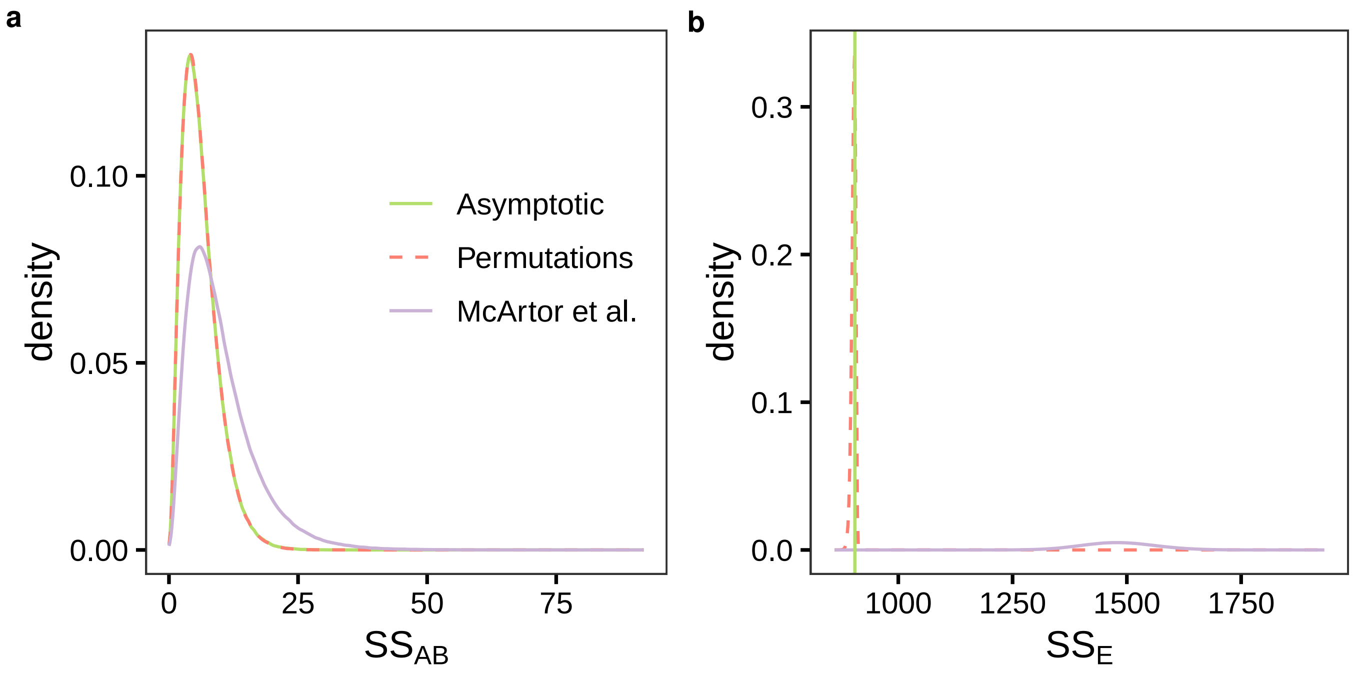


**Figure S3.** Null distribution of **a)** the numerator ($SS_{AB}$) and **b)** the denominator ($SS_E$) sums of squares of the PERMANOVA test statistic in (2). We simulated a scenario analogous to the one employed for Figure 1a and studied the test statistic for the interaction term (AB). We compared our asymptotic distribution derived from (3) (green solid line), with the proposal of McArtor et al.[34] (purple solid line) and the distribution obtained using $10^6$ permutations (red dashed line).

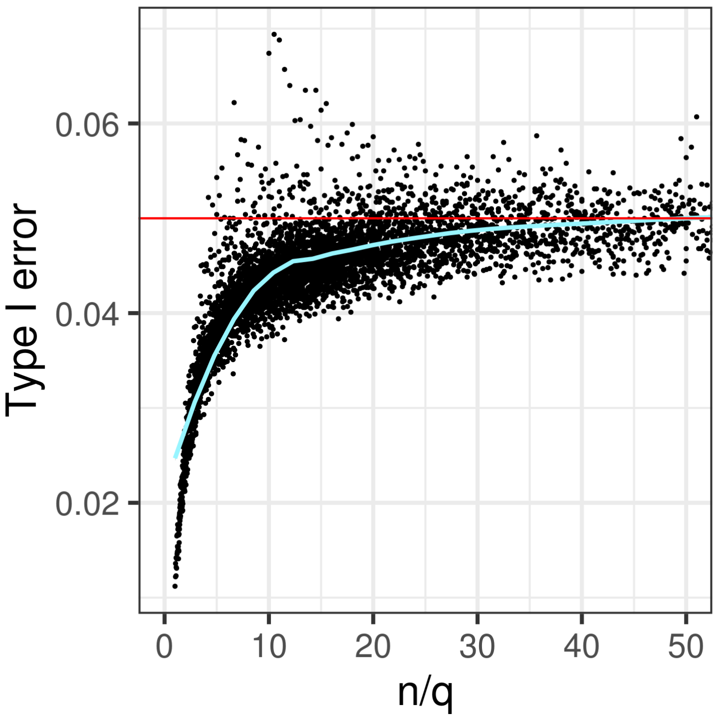


**Figure S4.** Type I error of asymptotic *p* values for the interaction term ($AB$) as a function of the $n/q$ ratio. We considered values of $n$ ranging from 20 to 300, and values of $q$ ranging from 2 to 20. For visualization purposes, we show the window $n/q \in$ [0,50]. Simulation details: $\boldsymbol{y} \sim \mathcal{N}(\mathbf{0}, \mathbf{I}_q)$, model (5) with factor $B$ simulated under $H_1$ and $\Delta =$ 1 (see Methods). The horizontal red line marks the 0. A polynomial has been fitted to the points using local fitting (LOESS), in order to describe the trend (fit shown in blue, 95% confidence interval in grey).

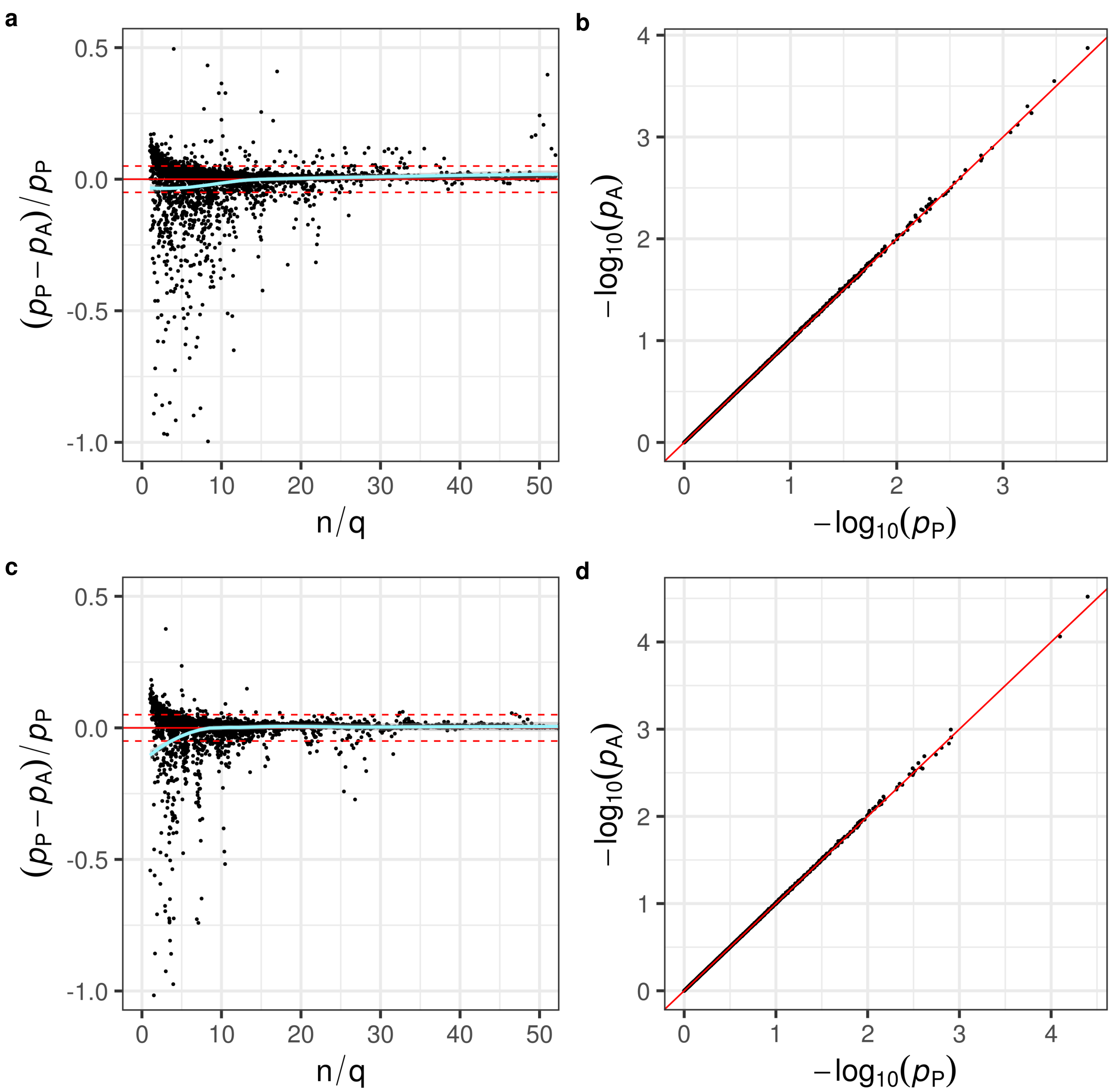


**Figure S5. a, c)** Relative bias of asymptotic *p* values *vs* $n/q$ ratio, for different distributions of the response variables: multivariate proportions (simplex) and Gaussian copula with uniform marginals, respectively. Relative difference between asymptotic ($p_A$) and permutation-based ($p_P$, $10^5$ permutations) *p* values for the interaction term ($AB$) as a function of the ratio between the total sample size and the number of dependent variables ($n/q$). We considered values of $n$ ranging from 20 to 300, and values of $q$ ranging from 2 to 20. For visualization purposes, we show values of $n/q \in$ [0,50] and relative biases $\in$ [-1,0.5]. The horizontal solid red line marks the 0. The horizontal dashed red lines mark the 5% relative bias. A polynomial has been fitted to the points using local fitting (LOESS), in order to describe the trend (fit shown in green, 95% confidence interval in grey). **b, d)** Comparison of asymptotic and permutation-based *p* values when the asymptotic null holds ($n = 300$, $q = 3$), for different distributions of the response variables: multivariate proportions (simplex) and Gaussian copula with uniform marginals, respectively. Simulation details: $B$ generated under $H_1$, with $\Delta$ set to 0.02 (simplex) or 1 (Gaussian copula).

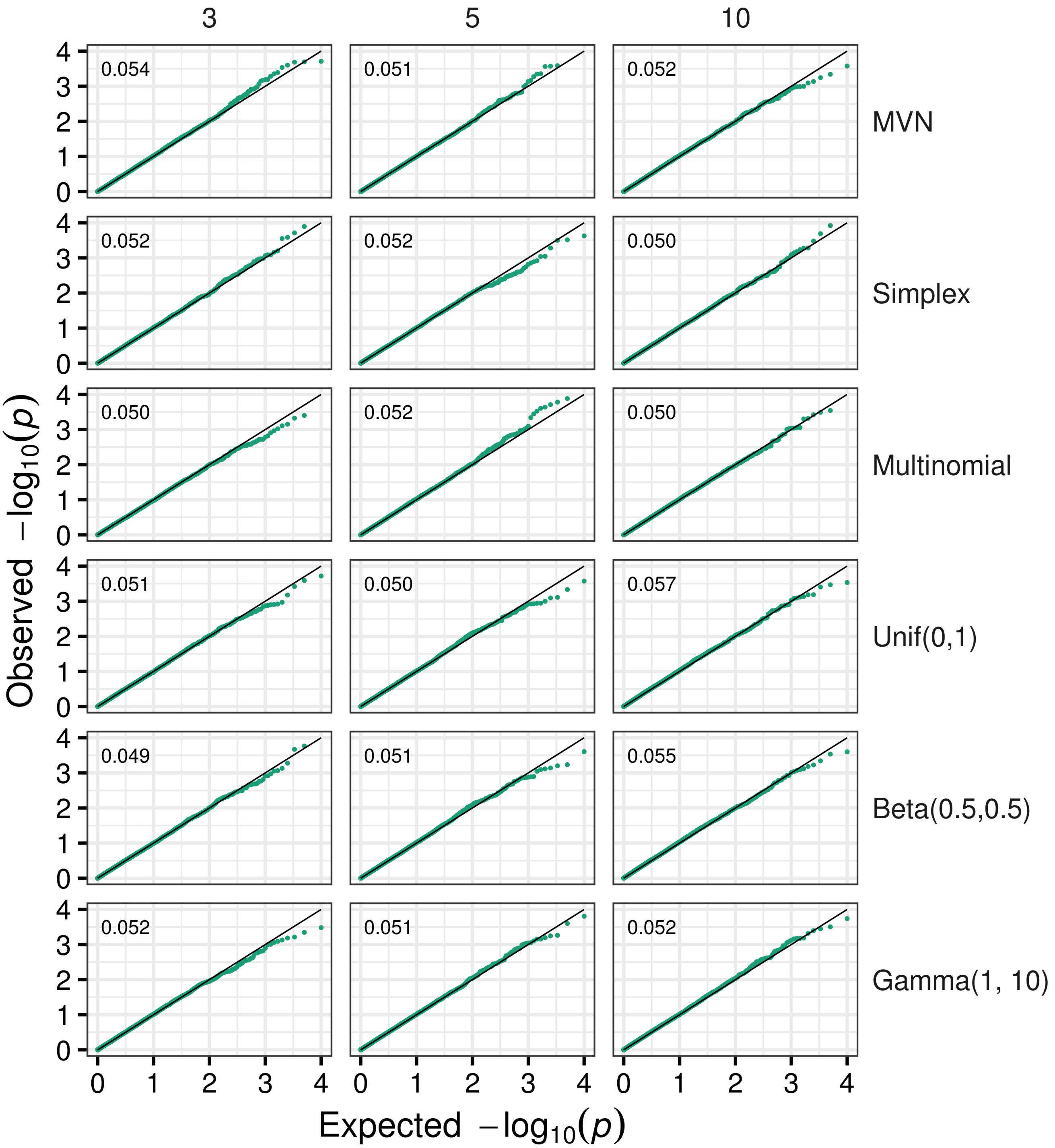


**Figure S6.** QQ-plots of *p* values obtained with MANTA in a cohort of unrelated individuals ($n$ = 1,000), across different numbers of traits (columns) and residual distributions (rows) (see Methods). Type I errors are also shown. In all simulations, $h_g^2$ was set to 0.2.

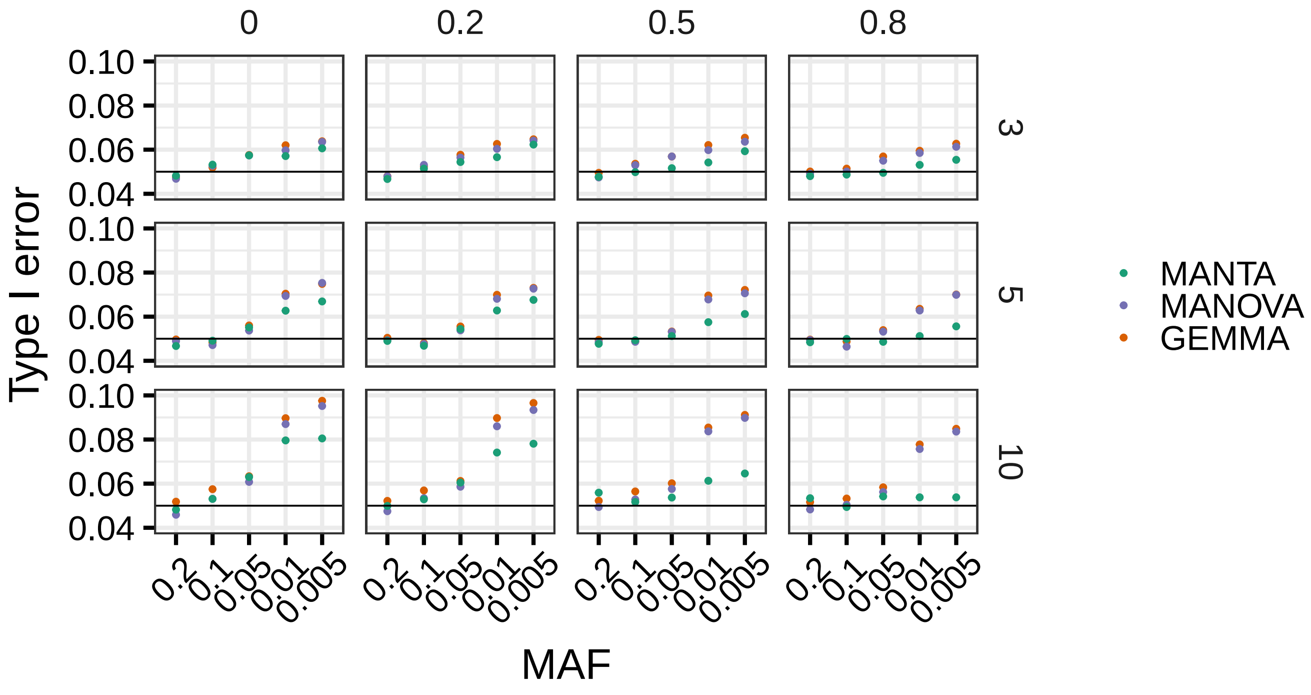


**Figure S7. a).** Type I error of MANTA (green), MANOVA (purple) and GEMMA (orange) as a function of the SNP minimum allele frequency (MAF), across different numbers of traits (rows) and pairwise trait correlations (columns). Simulation details: $n = 1{,}000$, $h_g^2 = 0.2$, multivariate $\boldsymbol{t}$ residuals (3 degrees of freedom), binomial SNPs (see Methods)

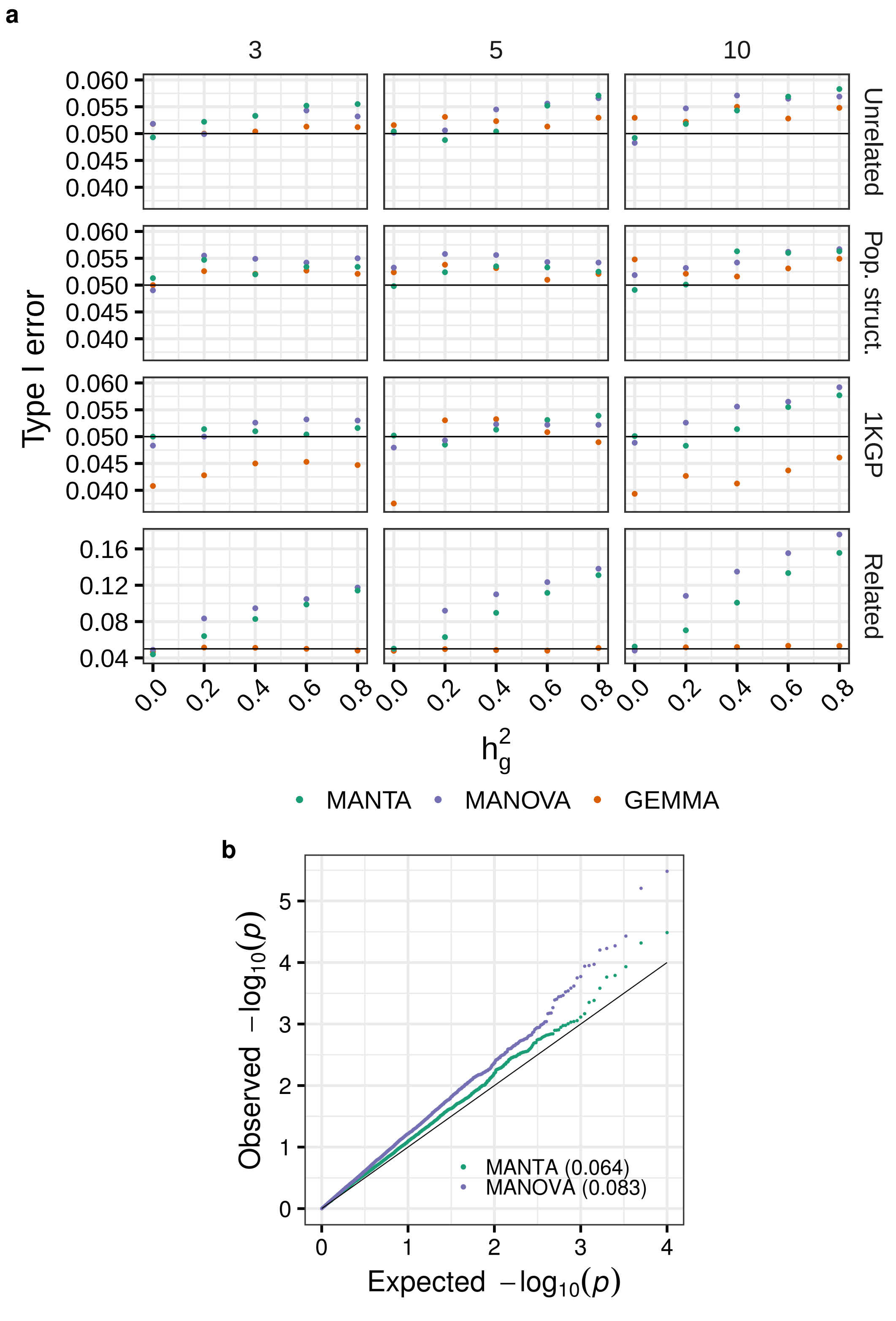


**Figure S8. a)** Type I error of MANTA (green), MANOVA (purple) and GEMMA (orange) as a function of the fraction of variance explained by population structure $h_g^2$, across different numbers of traits (columns) and population structures (rows): unrelated individuals ($n =$ 1,000), population stratification ($n =$ 1,000), actual 1000 Genomes Project dataset (1KGP, $n =$ 2,504) and related individuals ($n =$ 1,000). Simulation details: multivariate normal residuals, 5 genotype principal components (PCs) included as covariates in the model in the case of MANTA and MANOVA. **b)** QQ-plots of *p* values obtained with MANTA (green) and MANOVA (purple) in a cohort of related individuals, when the fraction of variance explained by population structure is small ($h_g^2 =$ 0.2). Type I errors are also shown. Simulation details as above.

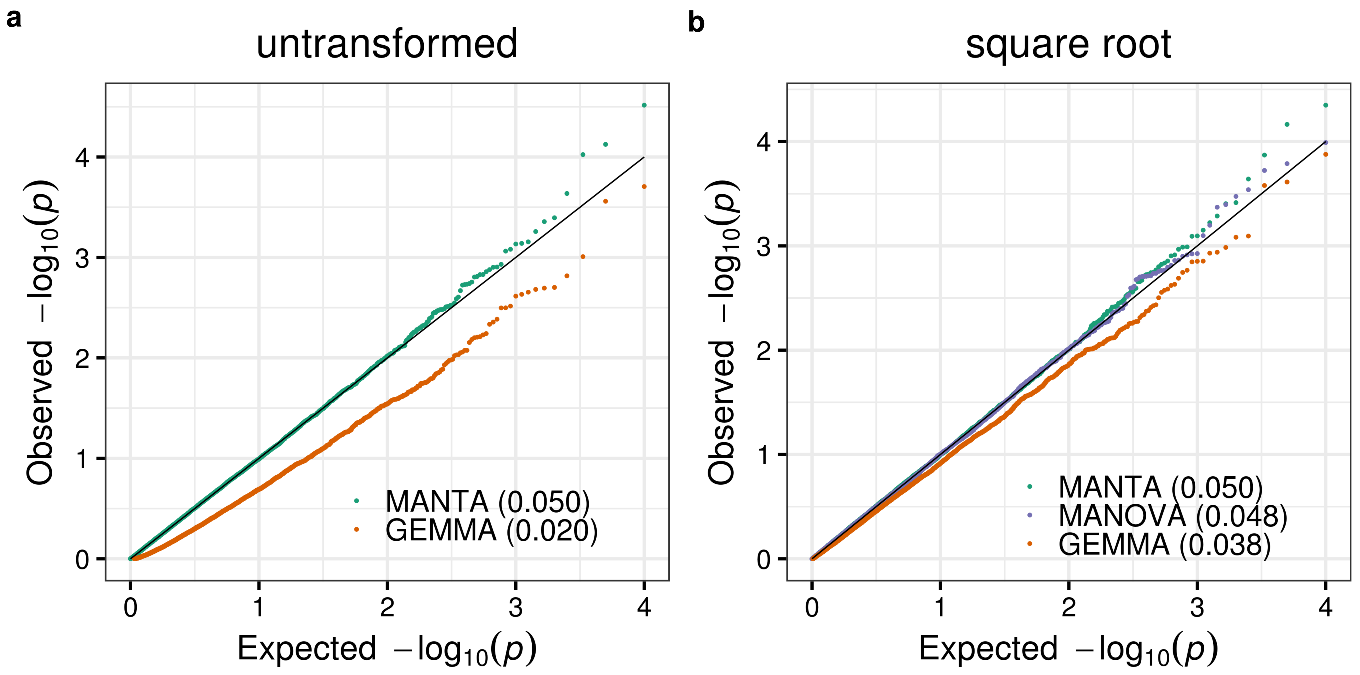


**Figure S9. a)** QQ-plots of $p$ values obtained with MANTA (green) and GEMMA (orange) when simulating multivariate proportion traits (not residuals, see Methods). Type I errors are also shown. Simulation details: actual 1000 Genomes Project genotypes ($n$ = 2,504), $q$ = 5, 5 genotype principal components (PCs) included as covariates in the model (in the case of MANTA). Note that, in this scenario, MANOVA $p$ values cannot be computed. **b)** Analogous representation when the traits are square root transformed. In this case the results of MANOVA are also displayed (purple). Simulation details as above.

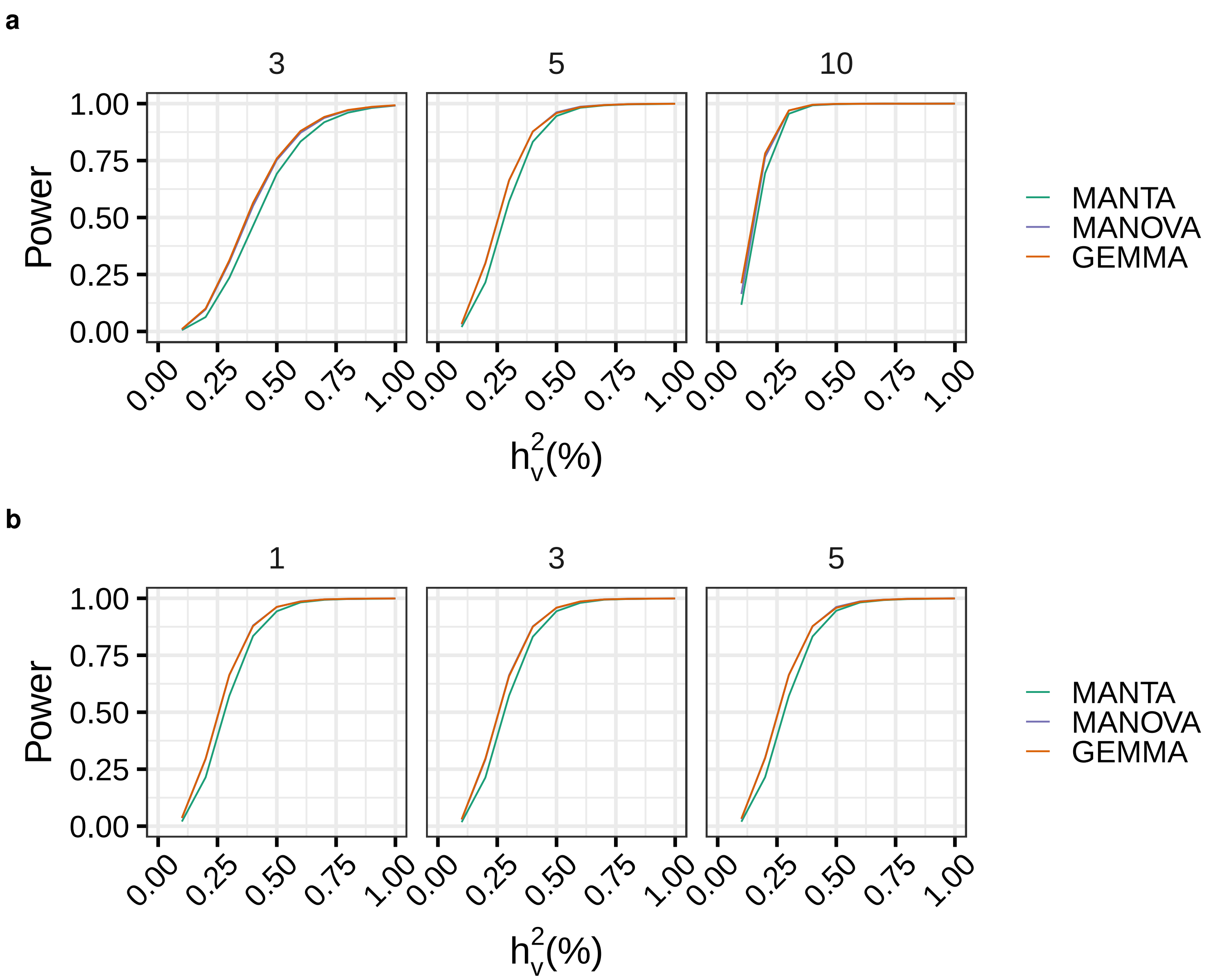


**Figure S10. a)** Power of MANTA (green), MANOVA (purple) and GEMMA (orange) as a function of the percentage of variance explained by the causal variant ($h_v^2$) and the total number of responses ($q$, all affected by the causal variant, columns) **b)** Analogous representation as a function of the number of affected responses, when this is different from $q$ (here $q = 5$). Simulation details: actual 1000 Genomes Project genotypes ($n =$ 2,504), multivariate normal residuals, $h_g^2 =$ 0.2, 5 genotype principal components (PCs) included as covariates in the model in the case of MANTA and MANOVA, Bonferroni corrected *p* values. GEMMA often fails (produces an error) when the number of traits is relatively large: in the $q =$10 scenario shown in a), on average, one out of two GEMMA tests resulted in an error. The fraction of errors in scenarios with fewer traits was substantially smaller (see Methods).

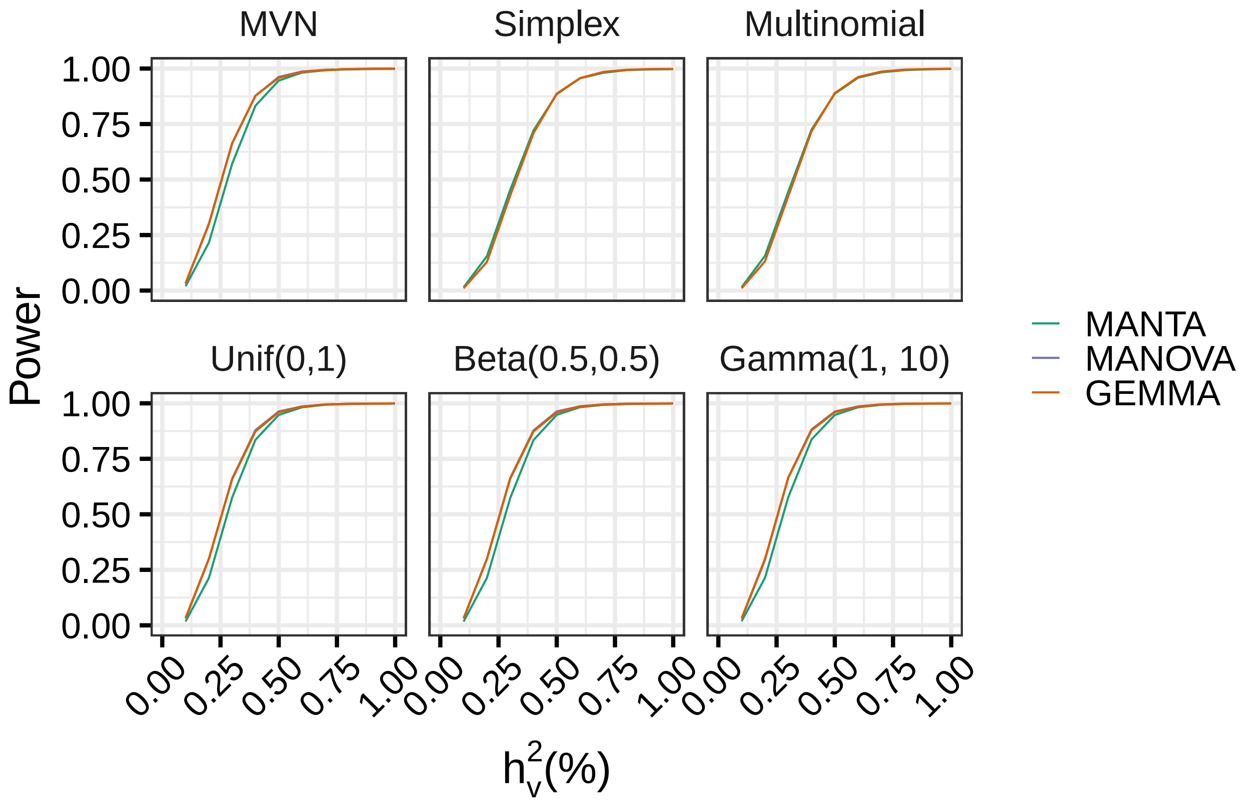


**Figure S11.** Power of MANTA (green), MANOVA (purple) and GEMMA (orange) as a function of the percentage of variance explained by the causal variant ($h_v^2$) and the residual distribution. Simulation details: actual 1000 Genomes Project genotypes ($n =$ 2,504), $q =$ 5 traits, $h_g^2 =$ 0.2, 5 genotype principal components (PCs) included as covariates in the model in the case of MANTA and MANOVA, Bonferroni corrected *p* values. Note that MANOVA *p* values cannot be calculated for linearly dependent traits, as it is the case of the simplex and multinomial scenarios.

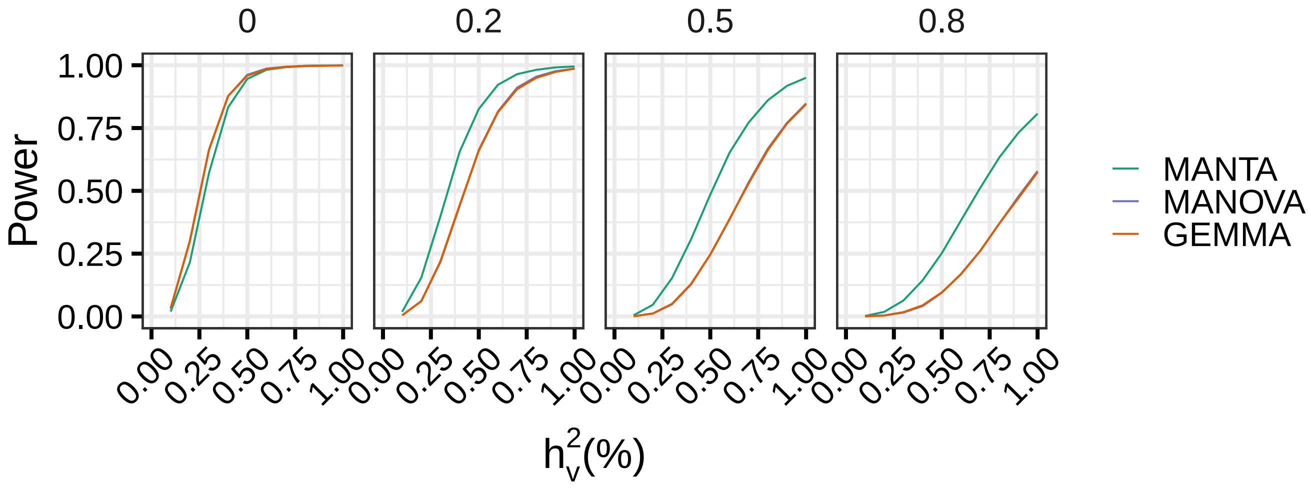


**Figure S12.** Power of MANTA (green), MANOVA (purple) and GEMMA (orange) as a function of the percentage of variance explained by the causal variant ($h_v^2$), for increasing values of trait-to-trait correlations. Simulation details: actual 1000 Genomes Project genotypes ($n =$ 2,504), $q =$ 5 traits, multivariate normal residuals, $h_g^2 =$ 0.2, 5 genotype principal components (PCs) included as covariates in the model in the case of MANTA and MANOVA, Bonferroni corrected *p* values.

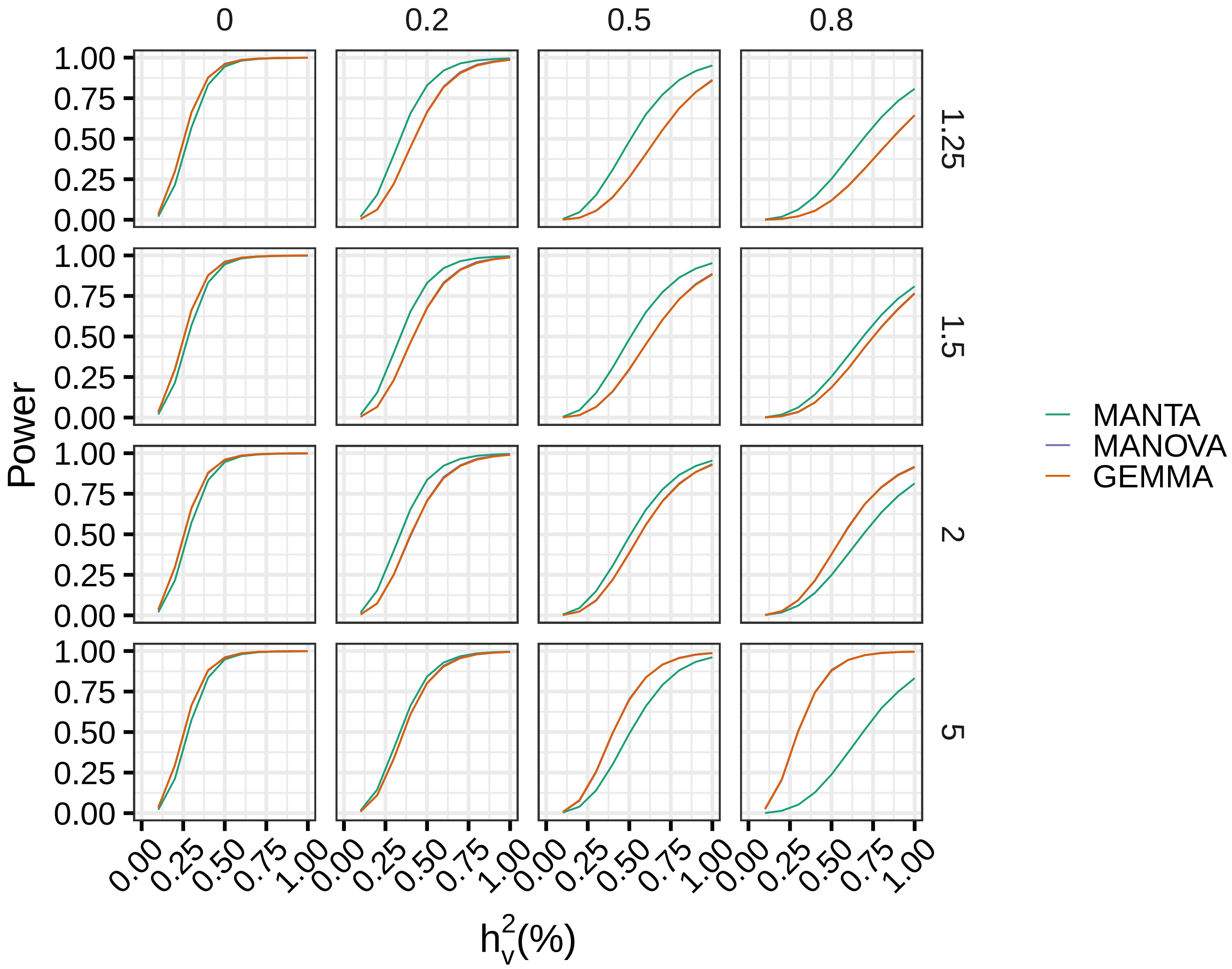


**Figure S13.** Power of MANTA (green), MANOVA (purple) and GEMMA (orange) as a function of the percentage of variance explained by the causal variant ($h_v^2$), across trait-to-trait correlations (columns) and different effect size max/min ratios (rows). A max/min effect size ratio of 1 corresponds to simulate equal (unit) effect sizes across all the traits, that is, effects are concordant with the trait correlation structure. As the max/min effect size ratio increases, the more discordant are genetic effects and trait correlations (see Methods). Simulation details: actual 1000 Genomes Project genotypes ($n =$ 2,504), $q =$ 5 traits, multivariate normal residuals, $h_g^2 =$ 0.2, 5 genotype principal components (PCs) included as covariates in the model in the case of MANTA and MANOVA, Bonferroni corrected *p* values.

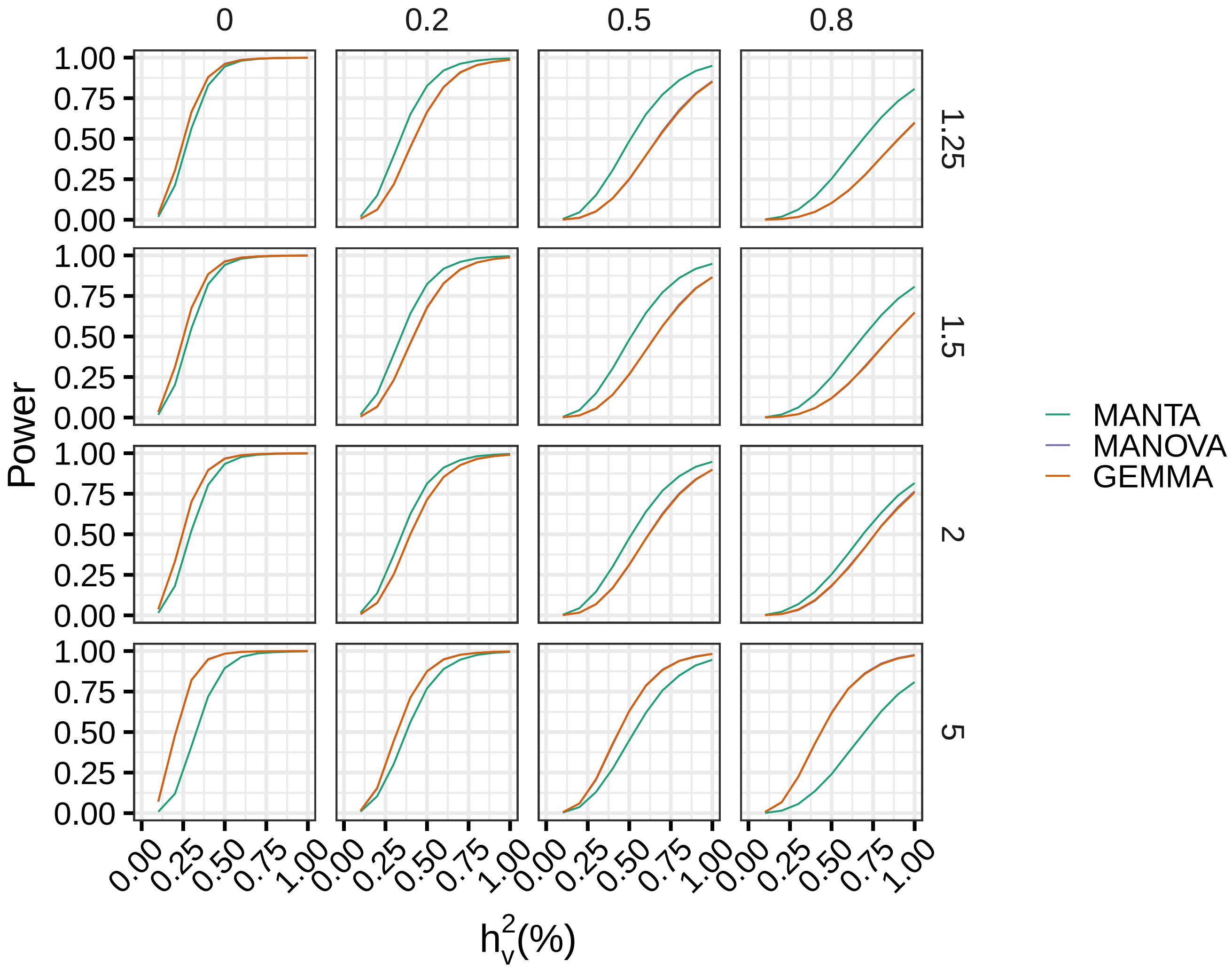


**Figure S14.** Power of MANTA (green), MANOVA (purple) and GEMMA (orange) as a function of the percentage of variance explained by the causal variant ($h_v^2$), across trait-to-trait correlations (columns) and trait max/min variance ratios (rows). A max/min variance ratio of 1 corresponds to equal (unit) variances simulated for all traits (see Methods). Simulation details: actual 1000 Genomes Project genotypes ($n =$ 2,504), $q =$ 5 traits, multivariate normal residuals, $h_g^2 =$ 0.2, 5 genotype principal components (PCs) included as covariates in the model in the case of MANTA and MANOVA, Bonferroni corrected *p* values.

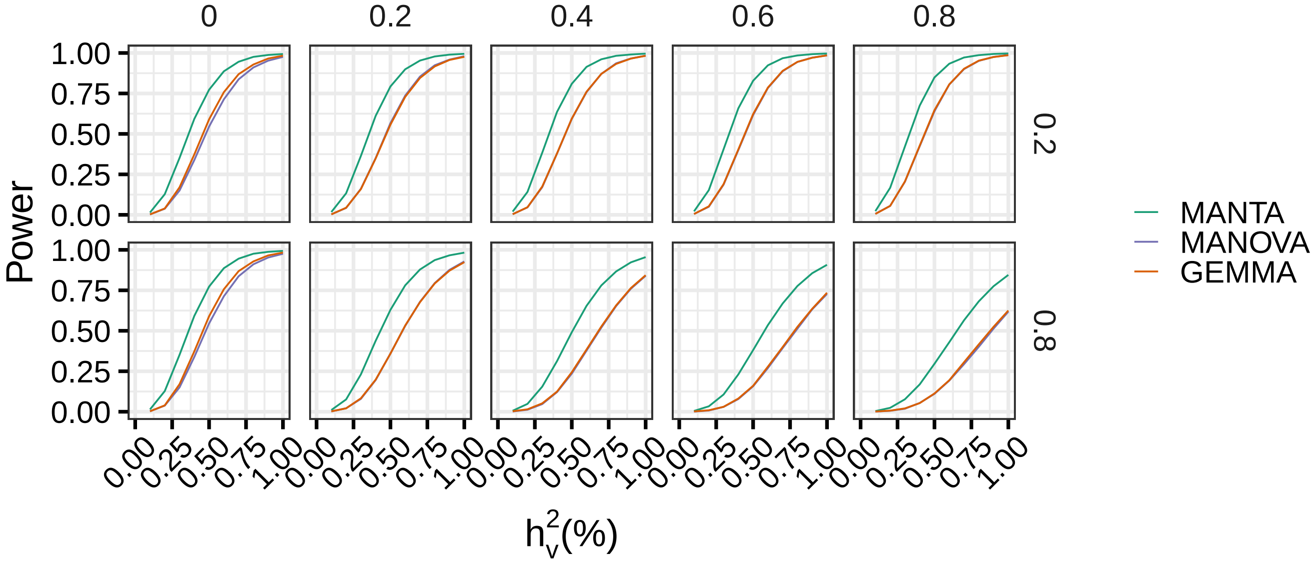


**Figure S15.** Power of MANTA (green), MANOVA (purple) and GEMMA (orange) as a function of the percentage of variance explained by the causal variant ($h_v^2$), for different values of the fraction of variance explained by population structure, $h_g^2$ (columns), and trait-to-trait correlations due to population structure (rows). Simulation details: actual 1000 Genomes Project genotypes ($n$ = 2,504), $q$ = 5 traits, multivariate normal residuals, 5 genotype principal components (PCs) included as covariates in the model in the case of MANTA and MANOVA, Bonferroni corrected *p* values.

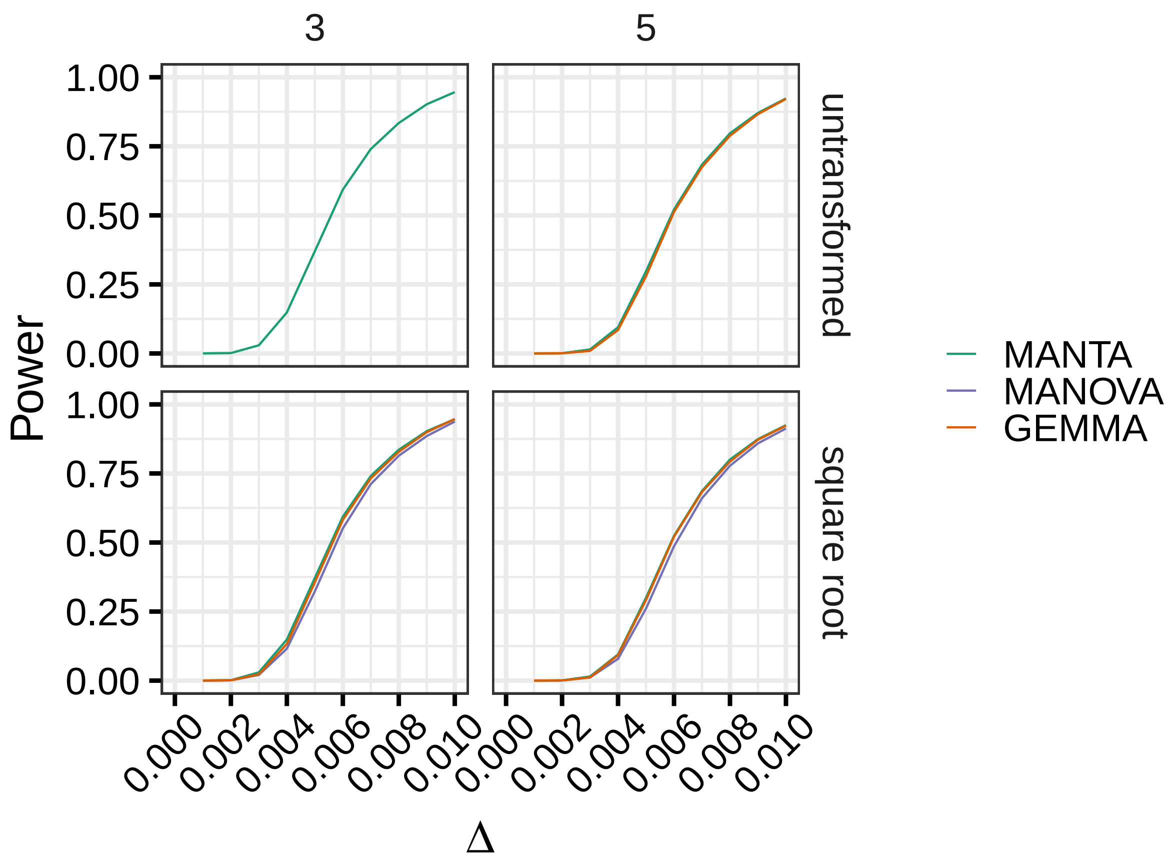


**Figure S16.** Power of MANTA (green), MANOVA (purple) and GEMMA (orange) as a function of $\Delta$, when simulating multivariate proportion traits (not residuals, see Methods), for different number of responses (columns) and trait transformations (rows). Simulation details: actual 1000 Genomes Project genotypes ($n = 2{,}504$), 5 genotype principal components (PCs) included as covariates in the model (in the case of MANTA), Bonferroni corrected $p$ values. Note that MANOVA $p$ values cannot be computed and that GEMMA often fails (produces an error and does not generate any result) with untransformed traits in this scenario (see Methods).

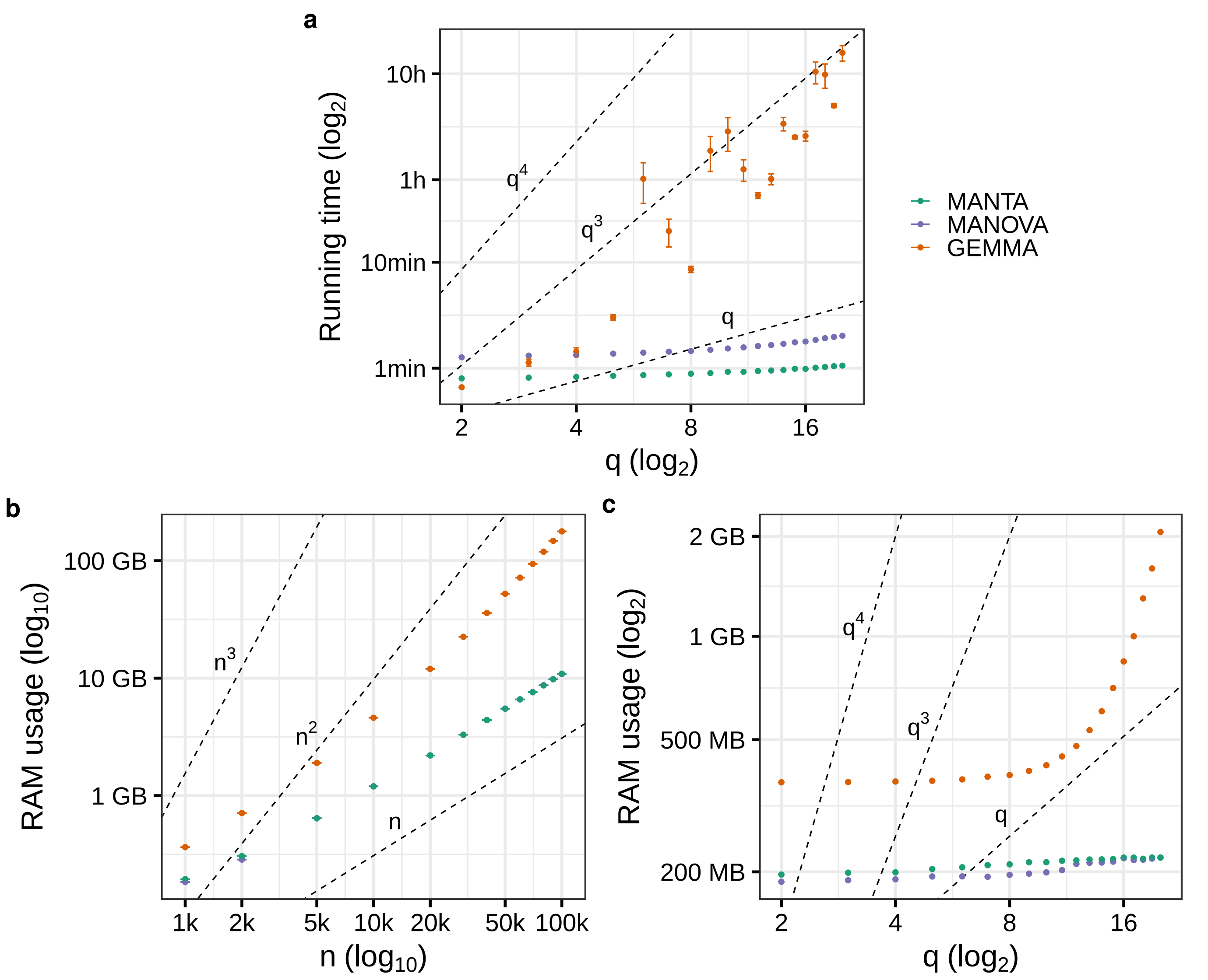


**Figure S17. a)** Empirical running time of MANTA (green), MANOVA (purple) and GEMMA (orange) as a function of the number of traits ($q$). Each point corresponds to the mean running time across 5 runs with different input data (see Methods). Error bars represent the standard error of the mean (i.e. mean $\pm$ SEM). Axes are in $\log_2$ scale. Dashed lines represent running time growth rates of $q$, $q^3$ and $q^4$. **b)**, **c)** Analogous representations for the RAM usage (peak resident set size) of the three methods as a function of the sample size ($n$) and the number of traits ($q$), respectively.

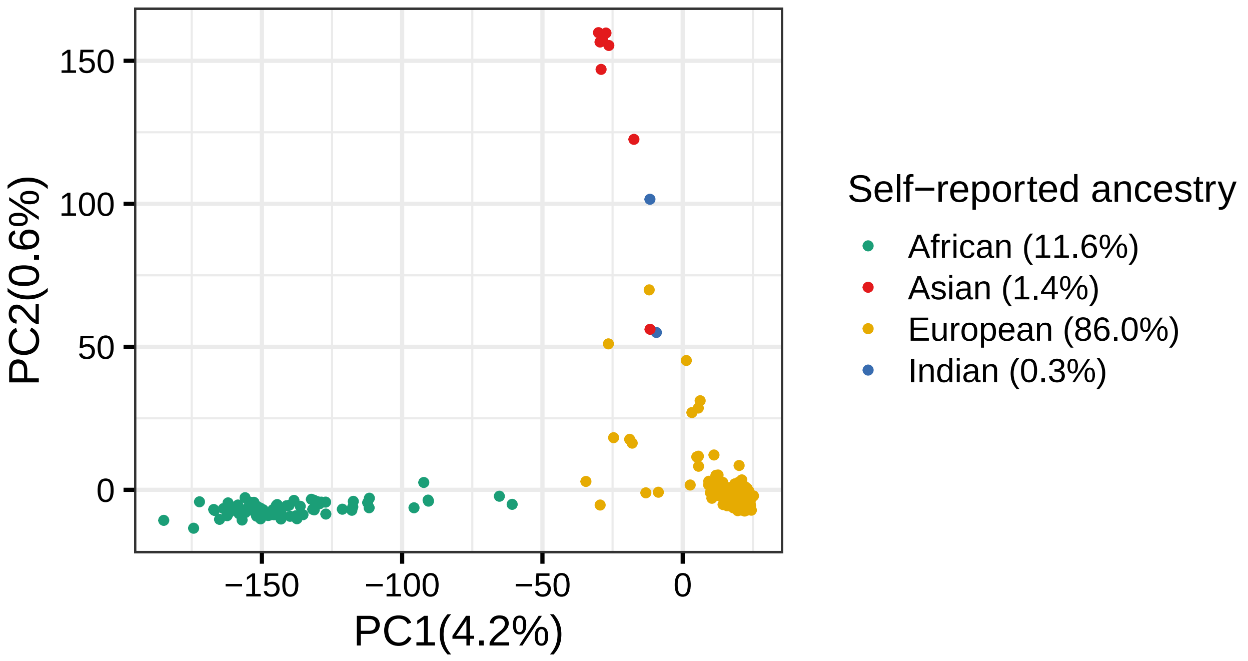


**Figure S18.** Principal Component Analysis (PCA) based on the genotypes of GTEx individuals. Colors represent self-reported ancestry (by the donor, family/next of kin or medical record). Individuals of unknown ancestry (0.6%) are not shown. The percentage of variance explained by the two first PCs and the percentage of individuals of each ancestry appear between parentheses. Self-reported ancestry matches the patterns observed in the PCA. Only individuals of European and African ancestry were used for pb-sQTL mapping.

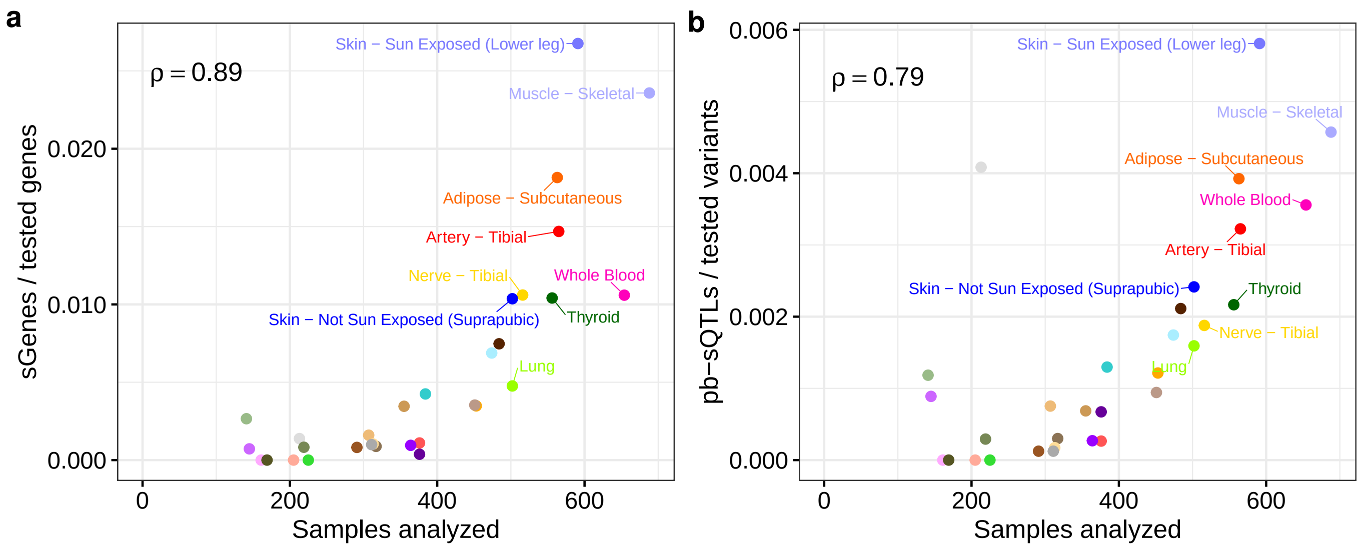


**Figure S19. a)**. Proportion of genes with pb-sQTLs (over tested genes, y-axis) per tissue with respect to the tissue sample size (x-axis). Spearman correlation between both metrics is also displayed. Tissues with sample size $> 500$ are labelled. Tissue color codes are available in Supplementary Table S1. **b)**. Analogous representation for the proportion of tested variants identified as pb-sQTLs.

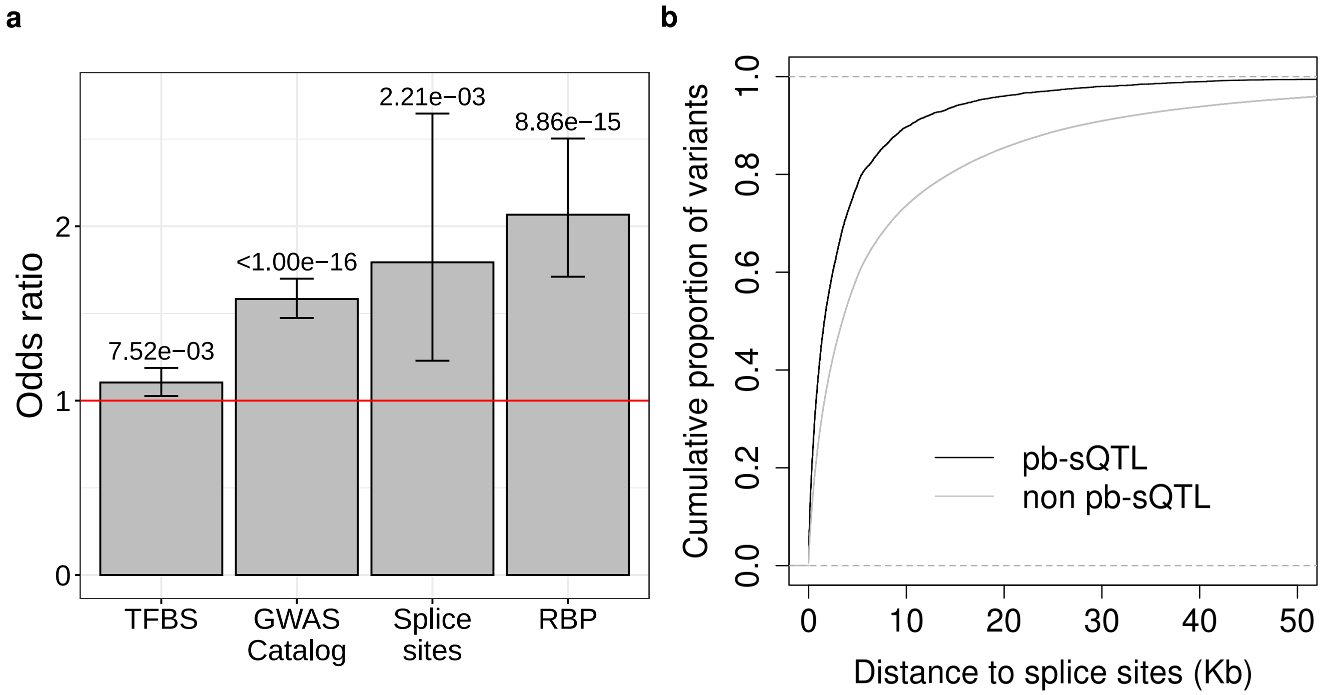


**Figure S20. a)** Enrichment of pb-sQTLs, with respect to a set of matched non pb-sQTLs (two-sided Fisher's exact test FDR $< 0.05$), in a set of functional categories: transcription factor binding sites (TFBS) from the Ensembl Regulatory Build, GWAS Catalog hits, splice sites from protein-coding and lincRNA genes annotated in GENCODE v26 and ENCODE RNA-binding protein (RBP) eCLIP peaks (see Methods). For each functional category, the height of the bar represents the enrichment odds ratio (OR), and the error bar its 95% confidence interval. FDR values are also displayed. **b)** Cumulative distribution of the distance to the closest splice donor or acceptor splice site from genes protein-coding genes and lincRNAs annotated in GENCODE v26, both for pb-sQTLs and non pb-sQTLs.

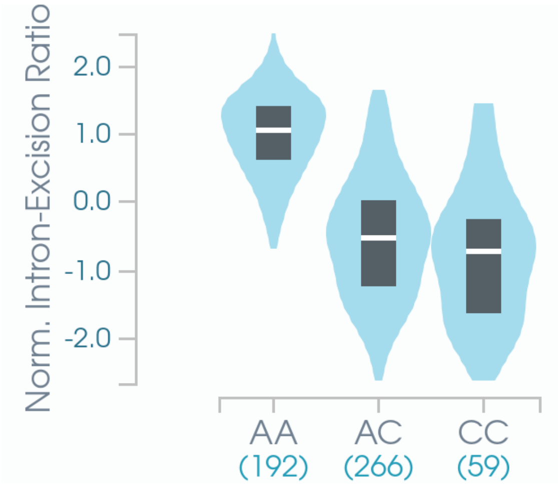


**Figure S21.** Normalized Leafcutter's intron-excision ratio (y-axis) for intron chr19:50,952,668-50,952,747 of *KLK5*, in individuals with different genotype at rs2739412 (chr19:50,952,558, A/C). The number of individuals with each genotype is shown between parentheses. This plot was directly obtained from the GTEx Portal (`https://gtexportal.org`).

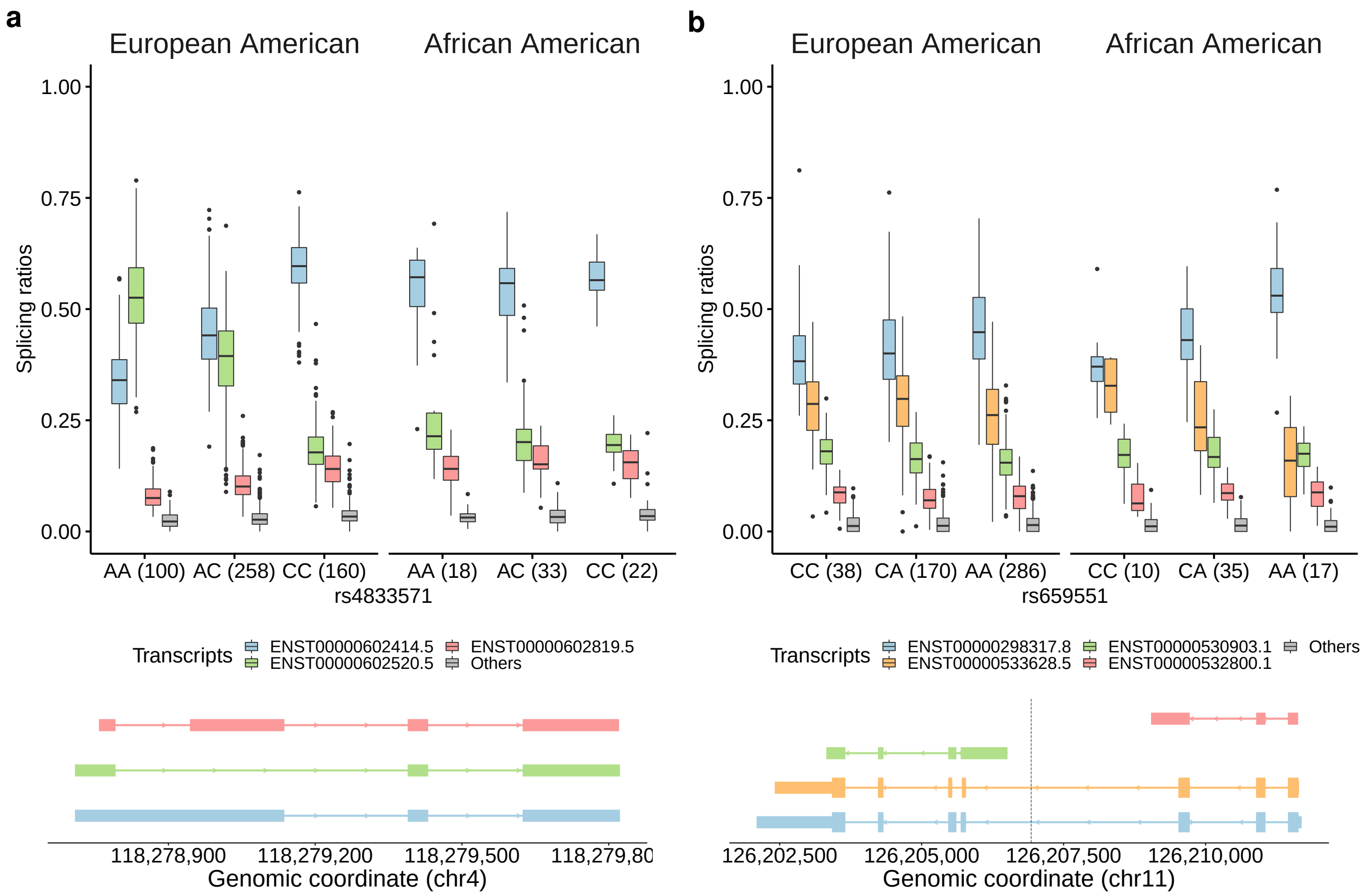


**Figure S22.** **a)** Relative abundances of the most expressed isoforms in skin (sun-exposed) from the lincRNA *SNHG8* (chr4:118,278,709-118,285,316, forward strand), for each ancestry (European American, EA, African American, AA) and genotype group at the rs4833571 locus (chr4:118,283,565 A/C), shown as boxplots. In boxplots, the box represents the first to third quartiles and the median, while the whiskers indicate $\pm$ 1.5 $\times$ interquartile range. The least abundant isoforms are grouped in Others. The number of individuals in each genotype group is shown between parentheses. In European Americans, reference homozygous (AA) individuals express preferentially the *ENST00000602520.5* (green) isoform, while in alternative homozygous (CC) individuals, isoform *ENST00000602414.5* (blue) captures most of the expression of the gene. Heterozygous EA individuals (TC) display an intermediate behaviour. In African Americans, however, isoform expression does not change with the genotype at rs4833571, with *ENST00000602414.5* (blue) being the most expressed isoform in all cases. The exonic structure of the most expressed isoforms of *SNHG8* in skin (sun-exposed) is also displayed. rs4833571 is an pb-sQTL for *SNHG8* in 6 additional tissues. **b)** Analogous representation for rs659551 (chr11:126,206,929, C/A), a pb-sQTL for gene *RPUSD4* (chr11:126,202,096-126,211,692, reverse strand, protein-coding) in thyroid. In this case, in African Americans, the abundance of isoform *ENST00000298317.8* (blue) increases, while the abundance of *ENST00000533628.5* (orange) decreases, with the number of alternative alleles (A) at the rs659551 locus. This behaviour can be barely observed in European Americans. *RPUSD4* is a thyroid-specific pb-sGene. The exonic structure of the most expressed isoforms of *RPUSD4* in thyroid is also shown, together with the location of the SNP (dotted vertical line).

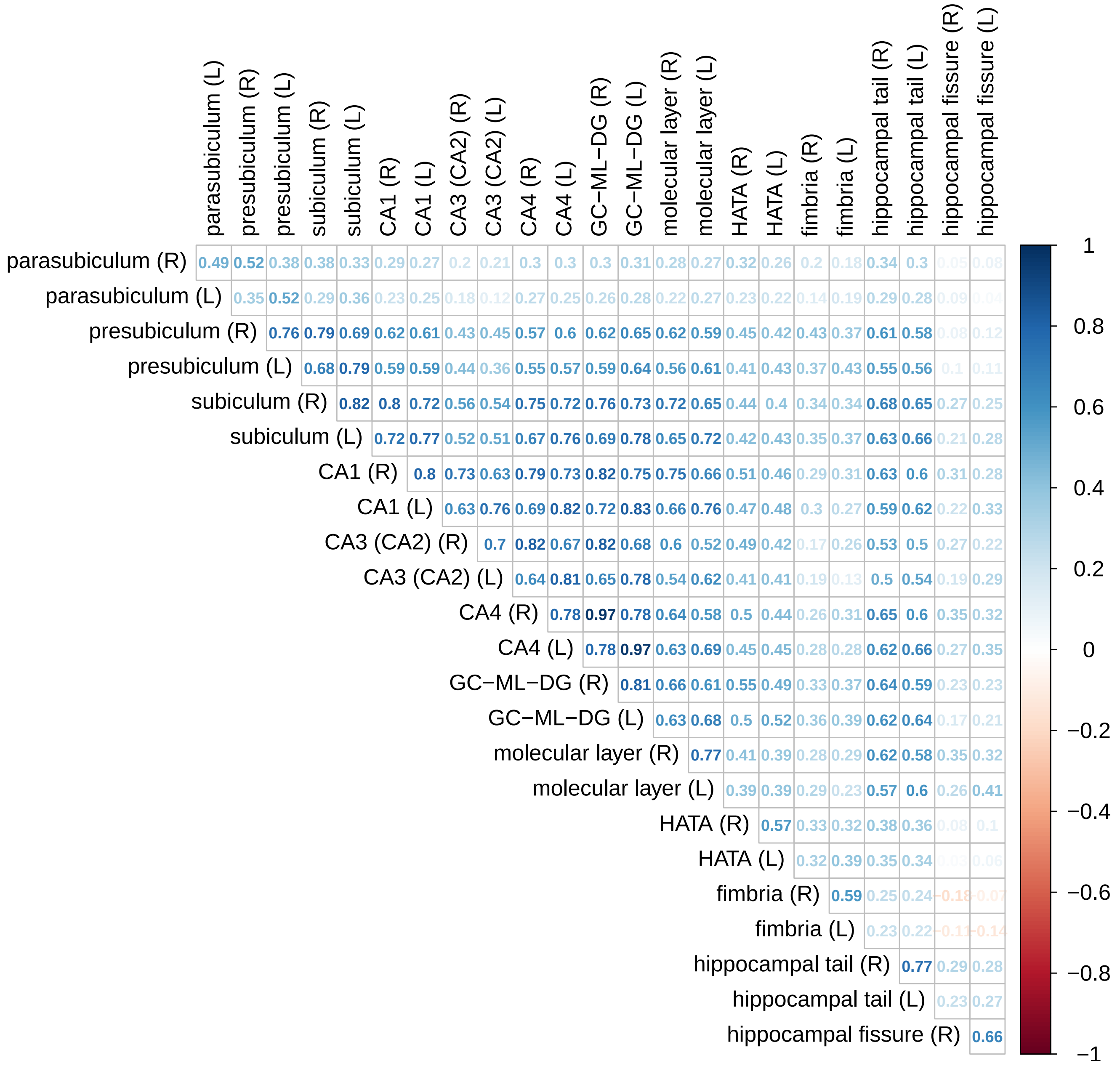


**Figure S23.** Pairwise Pearson correlation of the volume of hippocampal subfields in right (R) and left (L) brain hemispheres.

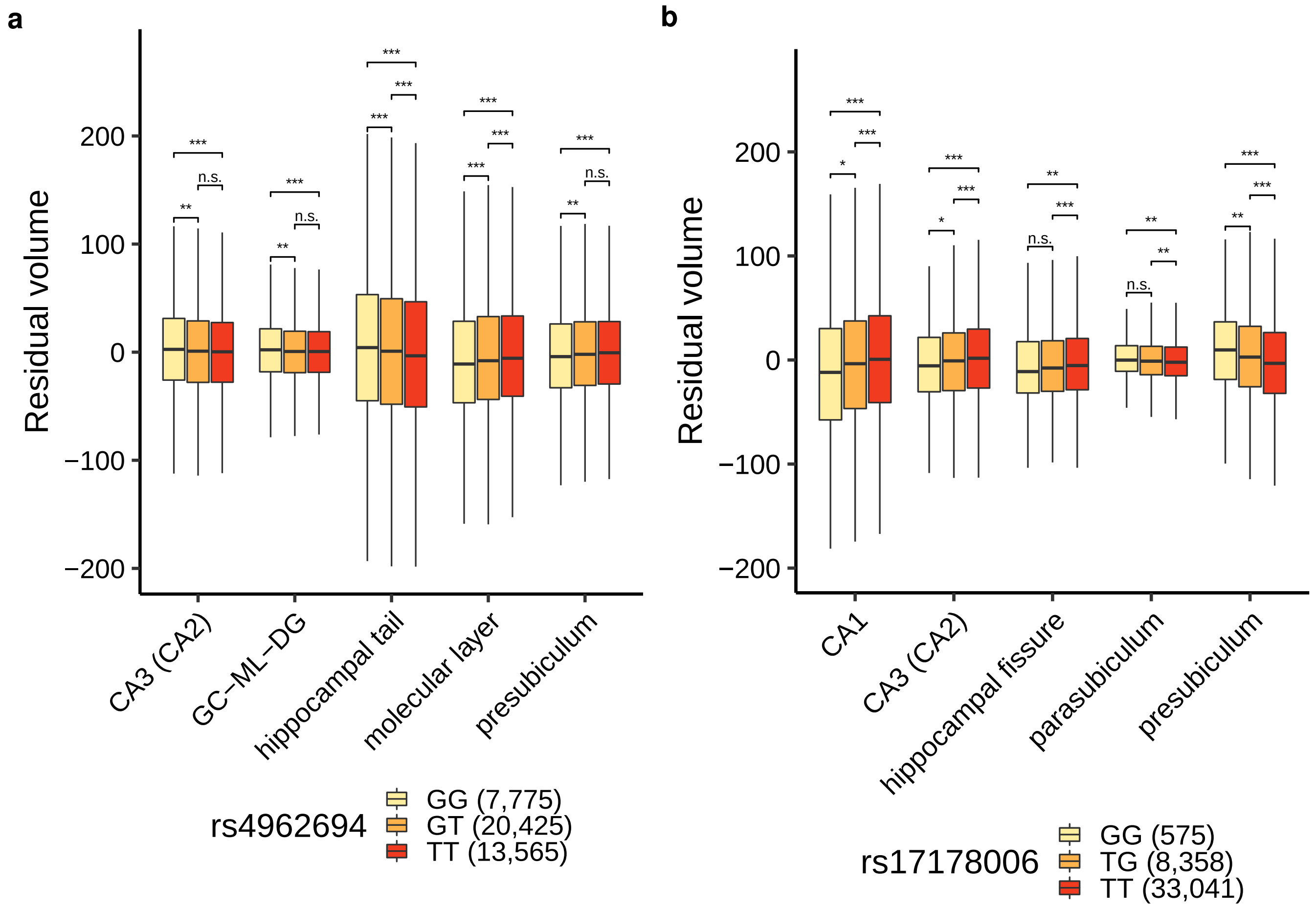


**Figure S24.** **a)** Covariate-adjusted volumes ($mm^3$) of the most changing hippocampal subfields across genotype groups at rs4962694 (chr10:126,436,717, G/T) are shown as boxplots. In boxplots, the box represents the first to third quartiles and the median, while the whiskers indicate $\pm$ 1.5 $\times$ interquartile range. The number of individuals on each group is shown between parentheses. FDR adjusted $p$ values (Wilcoxon Rank-Sum test) for each pairwise comparison are also displayed, encoded as follows: *** ($p \leq 0.001$), ** ($0.001 < p \leq 0.01$), * ($0.01 < p \leq 0.05$), n.s. ($p > 0.05$). For visualization purposes, outliers are not shown. rs4962694 was previously associated with the volume of the molecular layer, but not with the volume of other subfields[4]. **b)** Analogous representation for rs17178006 (chr12:65,718,299, T/G). This SNP was previously associated with the volume of presubiculum and CA1 (although associations with the volumes of CA3 (CA2), CA4, granule cell layer of the DG and hippocampal fissure were also reported when not correcting for total hippocampal volume)[4].

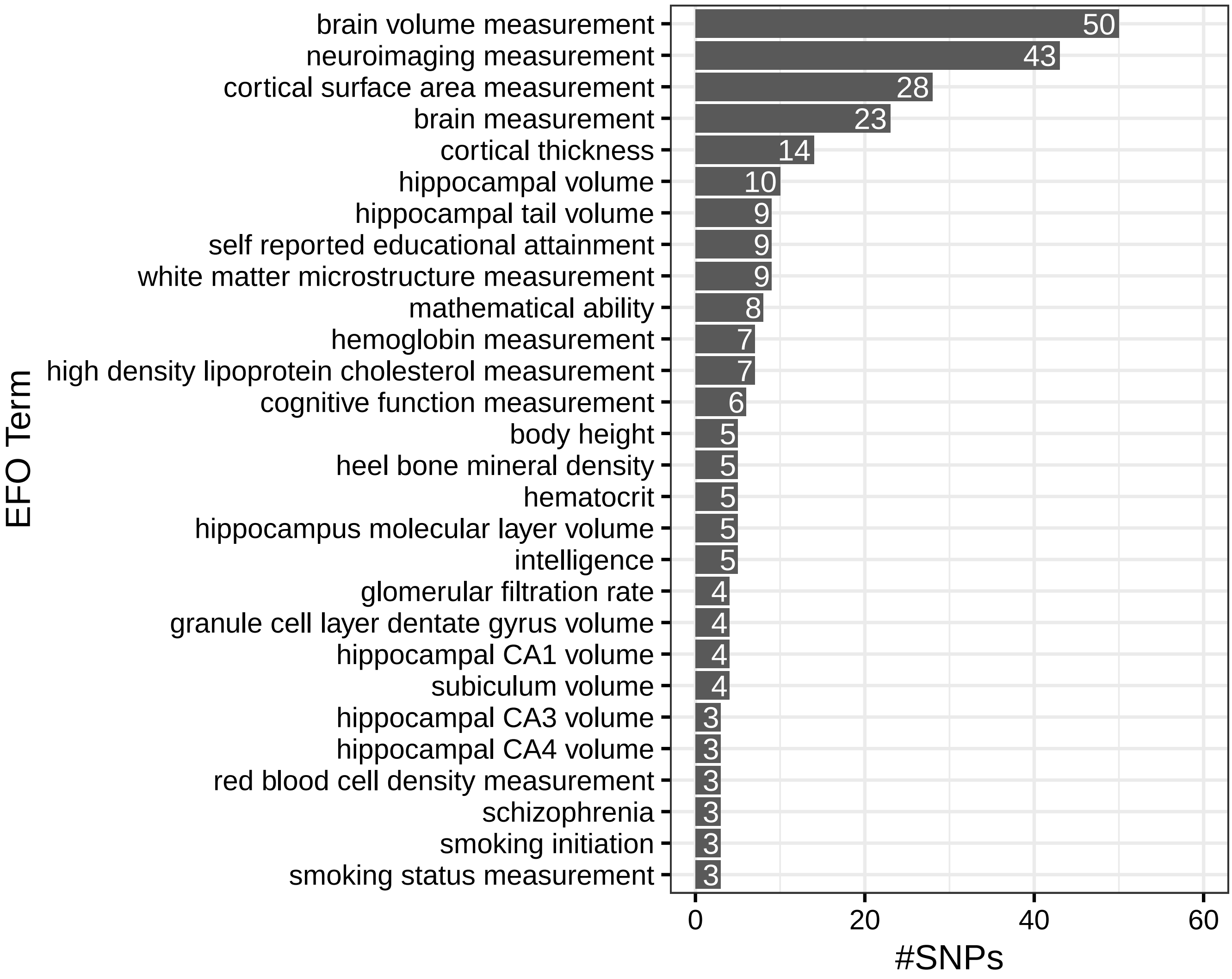


**Figure S25.** Top Experimental Factor Ontology (EFO) terms corresponding to the GWAS traits associated with the genetic variants identified by our approach. The number of genetic variants associated with each EFO term is also displayed. Note that the same genetic variant can be associated with more than one EFO term. Information retrieved from the GWAS Catalog (accessed 2021-01-29).

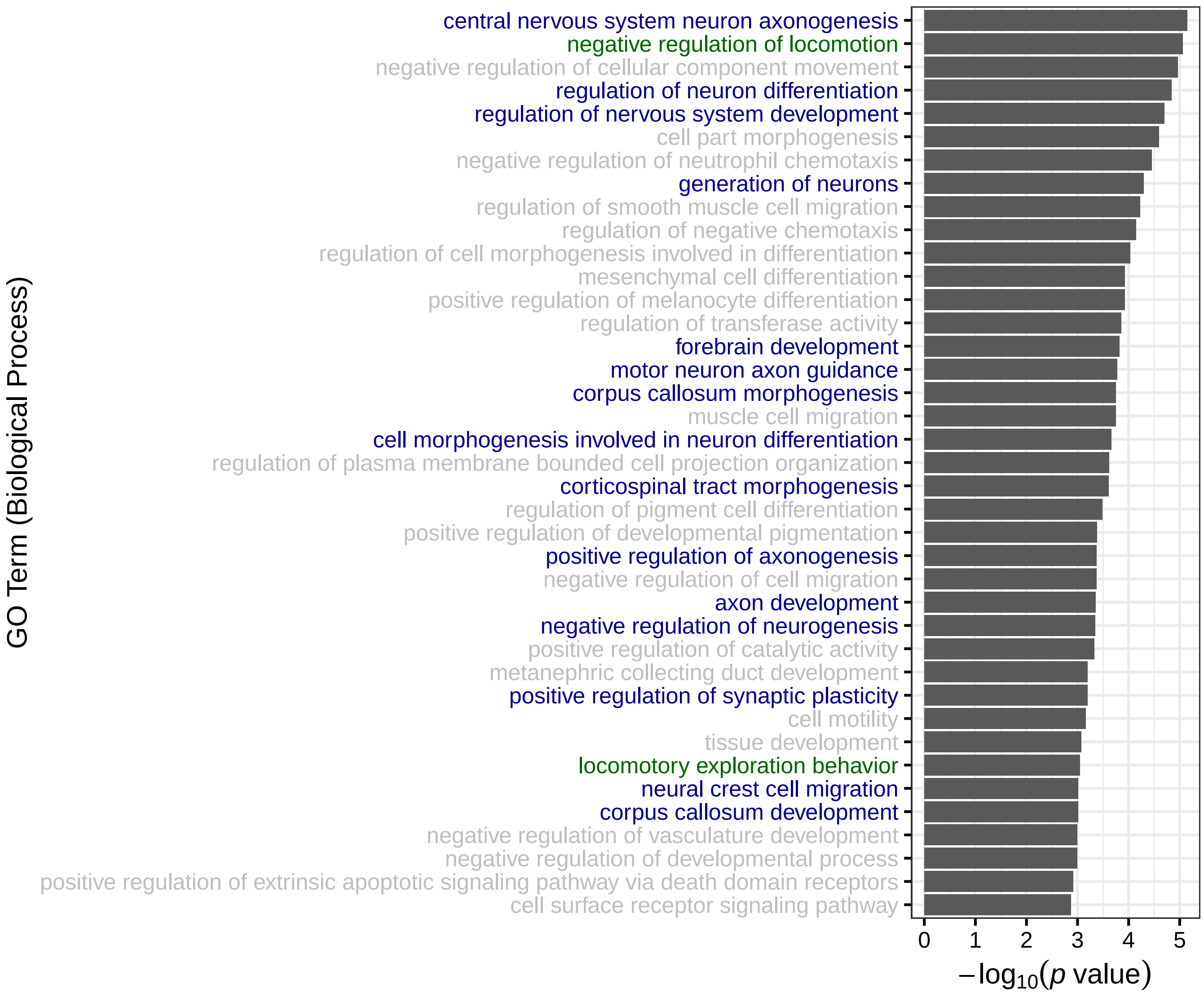


**Figure S26.** Gene ontology (GO) Biological Process terms enriched among the closest genes to the genetic variants associated with hippocampal subfield volumes (FDR $<$ 0.05). For each term (y-axis), the corresponding $-\log_{10}$ $p$ value (hypergeometric test) is shown (x-axis). Terms related to neuronal development and differentiation (blue), as well as to locomotive and exploratory behaviour (green) are highlighted.

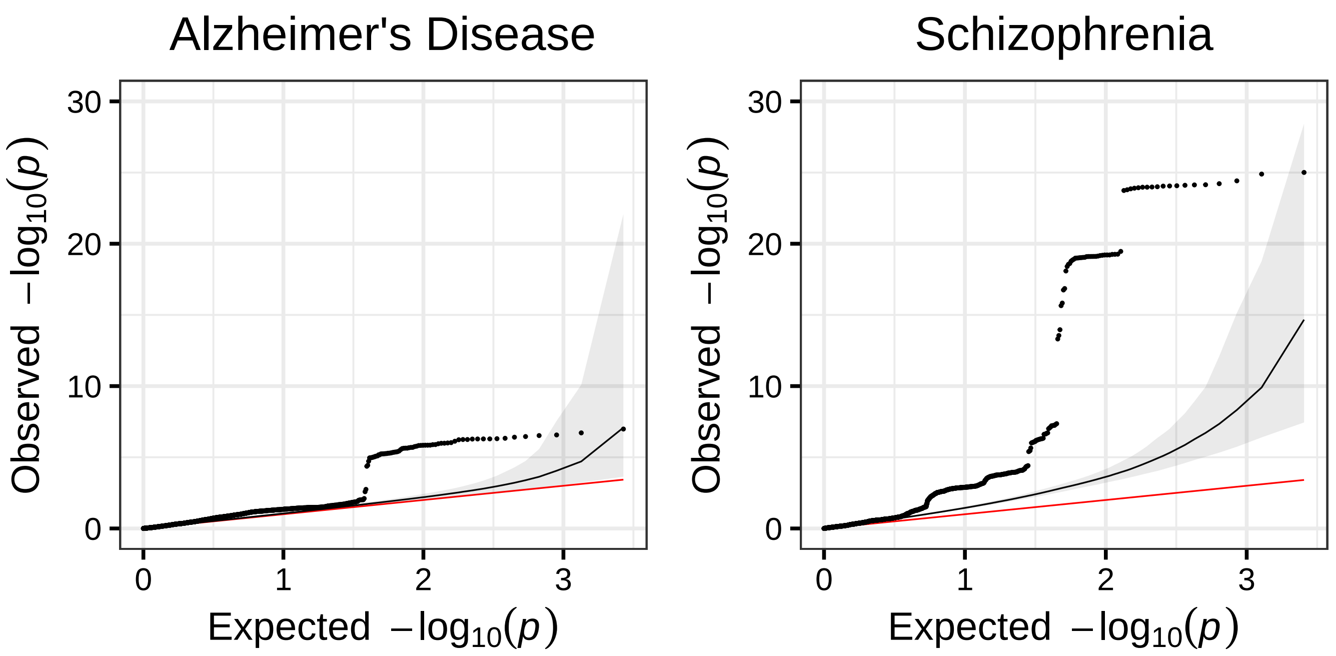


**Figure S27.** Quantile-quantile (QQ) plots of *p* values for association with Alzheimer's disease (left) and schizophrenia (right), for genetic variants associated with hippocampal subfield volumes (asymptotic PERMANOVA test *p* value $< 5 \cdot 10^{-8}$) plus variants in high linkage disequilibrium with them ($r^2 > 0.8$) (black dots), and the rest of variants (black line and grey area). The black line and grey area represent, respectively, the median and middle 95% observed $-\log_{10}$ *p* values across 10,000 random samplings from the latter variant set, with the same size as the former. The identity line is shown in red.

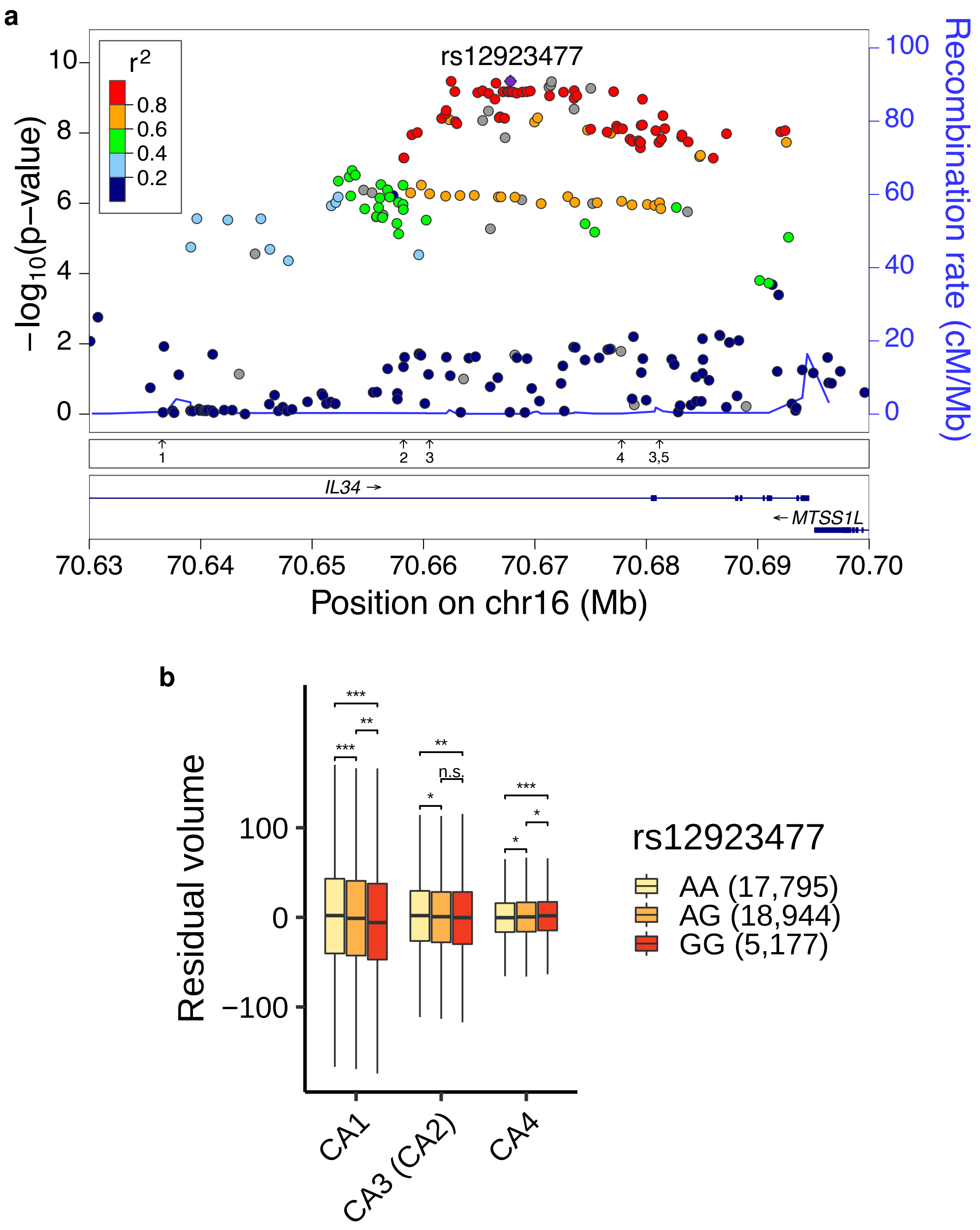


**Figure S28. a)** Regional plot of the *IL34* locus, showing the $-\log_{10}$ *p* values for association with hippocampal subfield volumes for all tested genetic variants (asymptotic PERMANOVA test). Linkage disequilibrium patterns (color-coded) and recombination rates are also displayed. The lower panels represent the location of previous associations with brain-related traits in the GWAS Catalog (shown as arrows), encoded as follows: 1) Hippocampal volume in Alzheimer's disease dementia, 2) Cerebrospinal P-tau181p levels, 3) Brain morphology, 4) Cortical surface area, 5) Subcortical volume. **b** Covariate-adjusted volumes (mm$^3$) of the most changing hippocampal subfields across genotype groups at rs12923477 (chr16:70,667,804, A/G) are shown as boxplots. In boxplots, the box represents the first to third quartiles and the median, while the whiskers indicate $\pm$ 1.5 $\times$ interquartile range. The number of individuals on each group is shown between parentheses. FDR adjusted *p* values (Wilcoxon Rank-Sum test) for each pairwise comparison are also displayed, encoded as follows: *** ($p \leq 0.001$), ** ($0.001 < p \leq 0.01$), * ($0.01 < p \leq 0.05$), n.s. ($p > 0.05$). For visualization purposes, outliers are not shown.

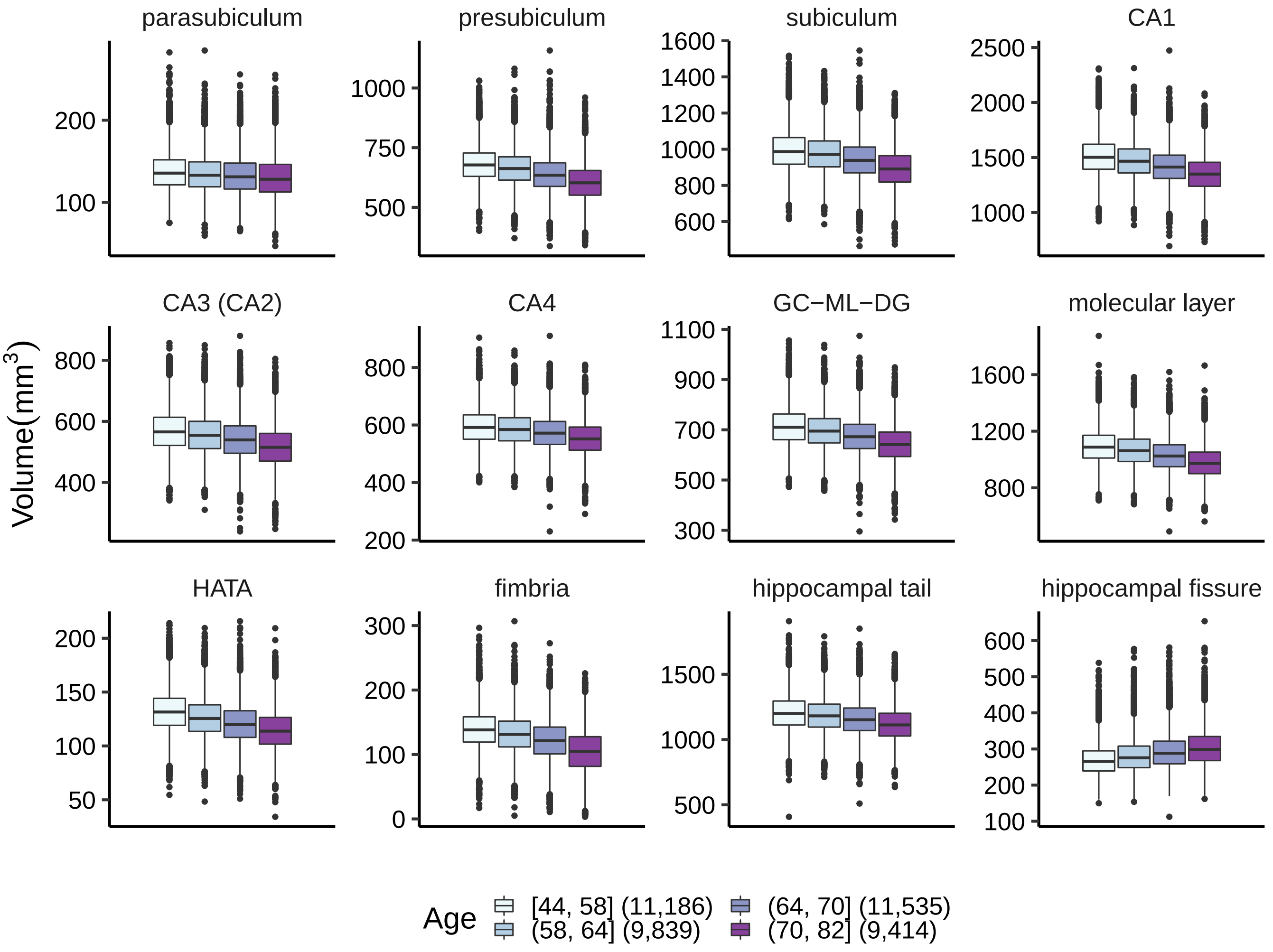


**Figure S29.** Distribution of the volumes (mm$^3$) of 12 hippocampal subfields, represented as boxplots, across four age ranges (corresponding to the quartiles of the age distribution). The number of individuals on each range is shown between parentheses. In boxplots, the box represents the first to third quartiles and the median, while the whiskers indicate $\pm$ 1.5 $\times$ interquartile range.

### Supplementary Note 1: Mathematical proofs

Consider the matrices **Y** and **X**, defined in the main text. Following Anderson, $\mathbf{D} = (d_{ij})$ is the inter-distances matrix between individuals, $\mathbf{A} = (a_{ij}) = \left(-\frac{1}{2}d_{ij}^2\right)$, then **G** in (9) is the matrix introduced by Gower[80]:

$$\mathbf{G} = \left(\mathbf{I} - \frac{1}{n}\mathbf{1}\mathbf{1}^{\mathrm{T}}\right)\mathbf{A}\left(\mathbf{I} - \frac{1}{n}\mathbf{1}\mathbf{1}^{\mathrm{T}}\right)$$

In this general context, Anderson defines the pseudo-F statistic as:

$$\tilde{\mathrm{F}} = \frac{\mathrm{tr}(\mathbf{HGH})/\mathrm{rank}(\mathbf{H})}{\mathrm{tr}((\mathbf{I}-\mathbf{H})\mathbf{G}(\mathbf{I}-\mathbf{H}))/\mathrm{rank}(\mathbf{I}-\mathbf{H})} \tag{9}$$

**Lemma 1.** If **Y** is a centered-column matrix, and **D** is computed using the Euclidean distance, the Anderson statistic in (9) can be expressed as:

$$\tilde{\mathrm{F}} = \frac{\mathrm{tr}(\mathbf{Y}^{\mathrm{T}}\mathbf{HY})/\mathrm{rank}(\mathbf{H})}{\mathrm{tr}(\mathbf{Y}^{\mathrm{T}}(\mathbf{I}-\mathbf{H})\mathbf{Y})/\mathrm{rank}(\mathbf{I}-\mathbf{H})}$$

*Proof.* If $\mathbf{Y} = (y_{ij})$ , any column mean $\bar{\mathbf{y}}_{\cdot j} = 0$. With the Euclidean distance and the definition of **G** it is straightforward to obtain:

$$\mathbf{G} = \mathbf{YY}^{\mathrm{T}}$$

The result is obtained combining the properties of the trace of the matrix product and the idempotence of the hat matrix **H**. □

Bai et al.[81] showed general results on the asymptotics of the m-estimation in the multivariate regression field. For our purposes, the main result in [81] is Theorem 2.4, which in our context can be simplified to:

**Theorem 1.** *Assume in model (1) the rows of* ***E*** *being i.i.d. with the same covariance matrix* $\Sigma$*. Its eigenvalue decomposition is* $\Sigma = \boldsymbol{P}\boldsymbol{\Lambda}\boldsymbol{P}^T$*,* $\Lambda = diag(\lambda_j)$*,* $vec(\boldsymbol{\beta}) = \boldsymbol{\beta}_v$ *is the vectorized form of the* $\beta$ *parameter matrix and* $\hat{\beta}_v$ *is the vector of corresponding estimates. Under mild regularity conditions of the design matrix, the following result holds:*

$$(\hat{\boldsymbol{\beta}}_v - \boldsymbol{\beta}_v)^T(\boldsymbol{P}\boldsymbol{P}^T) \otimes (\boldsymbol{X}^T\boldsymbol{X})(\hat{\boldsymbol{\beta}}_v - \boldsymbol{\beta}_v) \xrightarrow{d} \sum_{j=1}^{q} \lambda_j \chi_j^2(p) \tag{10}$$

*where* $\chi_j^2(p)$ *is a collection of* $q$ *independent chi-square variables with* $p$ *degrees of freedom,* $\otimes$ *denoting the Kronecker product of two matrices.*

*Proof.* The complete proof can be found in [81], however, their notation has some differences with respect to the usual MMR notation adopted here. To help the reading of [81], we translate some symbols and provide details of the most important matrices. First, model in [81] is stated as:

$$\mathbf{Y}_i = \mathbf{X}_{i_B}^{\mathrm{T}}\boldsymbol{\beta} + \mathbf{E}_i \qquad i = 1, \cdots, n$$

$\mathbf{E}_i$ stands for a i.i.d. $q$-vector of errors, $\mathbf{X}_{i_B}$ is a $m \times q$ design matrix. Some other noticeable differences between notations are:

1. Bai's model equates the response of an *individual* $i$ sample, while model (1) stands for the full sample of $n$ individuals.

2. Bai's $m$ stands for the dimension of $\boldsymbol{\beta}$, then, $m = p \times q$.

Denoting in model (1) row $i$ of $\mathbf{X}$ as $\mathbf{X}_i$ (without $B$), both design matrices are related by:

$$\mathbf{X}_{i_B}^{\mathrm{T}} = \mathbf{I}_q \otimes \mathbf{X}_i$$

The mild regularity conditions described in the proposition refer specifically to condition (M6) in [81], that is, if $\mathbf{S}_n = \mathbf{X}_{1_B}\mathbf{X}_{1_B}^{\mathrm{T}} + \cdots + \mathbf{X}_{n_B}\mathbf{X}_{n_B}^{\mathrm{T}}$ then $\mathbf{S}_n$ must be non-singular for $n \geq n_0$ and

$$d_n^2 = \max_{1 \leq i \leq n} tr(\mathbf{X}_{i_B} \boldsymbol{S}_n^{-1} \mathbf{X}_{i_B}^{\mathrm{T}}) \to 0 \qquad as \quad n \to \infty$$

which can be interpreted as the individual design matrix having a leverage tending to zero, something that seems reasonable in practice. The remaining conditions (M1) to (M5) in [81] are trivially satisfied here. Finally, the auxiliary matrices $\mathbf{K}_n$ and $\mathbf{T}_n$ in the proof in [81] are, respectively:

$$\mathbf{K}_n = \mathbf{I}_q \otimes (\mathbf{X}^{\mathrm{T}}\mathbf{X})$$

$$\mathbf{T}_n = \boldsymbol{\Sigma} \otimes (\mathbf{X}^{\mathrm{T}}\mathbf{X})$$

□

The covariance structure of the data in the first assumption of the theorem implies a global covariance matrix equal to $cov(vec(\mathbf{E})) = \mathbf{I} \otimes \boldsymbol{\Sigma}$ ($\mathbf{I}$ is the $n \times n$ identity matrix).

Under the omnibus null hypothesis, $\boldsymbol{\beta}_{\mathrm{v}} = \mathbf{0}$, the limiting distribution in (10) allows to obtain the trace in the numerator of the test statistic in (4):

$$\mathrm{tr}(\mathbf{Y}^{\mathrm{T}}\mathbf{H}\mathbf{Y}) = \hat{\boldsymbol{\beta}}_{\mathrm{v}}^{\mathrm{T}}(\mathbf{P} \otimes (\mathbf{X}^{\mathrm{T}}\mathbf{X})^{\frac{1}{2}})(\mathbf{P}^{\mathrm{T}} \otimes (\mathbf{X}^{\mathrm{T}}\mathbf{X})^{\frac{1}{2}})\hat{\boldsymbol{\beta}}_{\mathrm{v}} \xrightarrow{d} \sum_{j=1}^{q} \lambda_j \chi_j^2(p)$$

Consider now a null hypothesis where the parameters from $p_0 + 1$ to $p$ are zero for all the $q$ dimensions, that is, a hypothesis on a subset of the $p \times q$ possible parameters. Under this null hypothesis, for any single column of $\mathbf{Y}$, the corresponding column in matrix $\boldsymbol{\beta}$ in equation (1) will be multiplied by:

$$\mathbf{R} = \left(\mathbf{0}_{(p-p_0)\times p_0}\, , \, \mathbf{I}_{p-p_0}\right)$$

And joining all the dimensions we have the matrix $\mathbf{R}_{\mathrm{v}}$:

$$\mathbf{R}_{\mathrm{v}} = \mathbf{I}_q \otimes \mathbf{R}$$

which allows to express synthetically the hypothesis in the following Lemma, that specifies the limiting distribution of the Anderson's test for any subset of parameters.

**Lemma 2.** Assume a null hypothesis where all the parameters from columns $p_0+1$ to $p$ in (1) are zero for all the $q$ dimensions, that is:

$$\mathbf{R}_{\mathrm{v}}\,\boldsymbol{\beta}_{\mathrm{v}} = \mathbf{0}$$

then the numerator of the statistic in (2) converges in law to:

$$\mathrm{tr}\left\{\mathbf{Y}^{\mathrm{T}}\left(\mathbf{X}(\mathbf{X}^{\mathrm{T}}\mathbf{X})^{-1}\mathbf{X}^{\mathrm{T}} - \mathbf{X}_0(\mathbf{X}_0^{\mathrm{T}}\mathbf{X})_0^{-1}\mathbf{X}_0^{\mathrm{T}}\right)\mathbf{Y}\right\} \xrightarrow{d} \sum_{j=1}^{q}\lambda_j\chi_j^2(p-p_0)$$

*Proof.* The demonstration has three parts. First, consider the design matrix partitioned in two boxes $\mathbf{X} = (\mathbf{X}_0, \mathbf{X}_1)$ where $\mathbf{X}_0$ corresponds to $\mathbf{X}$ without the columns associated to the subset of parameters assumed to be zero. Then, prove that $\mathbf{H} - \mathbf{H}_0$ is an idempotent matrix.

Second, obtain the limiting distribution of $\mathbf{R}_{\mathrm{v}}\,(\hat{\boldsymbol{\beta}}_{\mathrm{v}} - \boldsymbol{\beta}_{\mathrm{v}})$ applying Theorem 1 and the *Product Limit Normal Rule*, and then derive the convergence in law of the following expression:

$$\left(\mathbf{I}_q \otimes \left(\mathbf{R}\,(\mathbf{X}^{\mathrm{T}}\mathbf{X})^{-1}\,\mathbf{R}^{\mathrm{T}}\right)^{-\frac{1}{2}}\right)\mathbf{R}_{\mathrm{v}}\,(\hat{\boldsymbol{\beta}}_{\mathrm{v}} - \boldsymbol{\beta}_{\mathrm{v}}) \xrightarrow{d} N(\mathbf{0},\, \boldsymbol{\Sigma}\otimes\mathbf{I}_{p-p_0})$$

Third, consider again the block partitioning of $\mathbf{X}$ in boxes $\mathbf{X} = (\mathbf{X}_0, \mathbf{X}_1)$ and prove that

$$\begin{aligned}\mathrm{tr}(\hat{\boldsymbol{\beta}}^{\mathrm{T}}\mathbf{X}^{\mathrm{T}}\mathbf{X}\hat{\boldsymbol{\beta}} - \hat{\boldsymbol{\beta}}_0^{\mathrm{T}}\mathbf{X}_0^{\mathrm{T}}\mathbf{X}_0\hat{\boldsymbol{\beta}}_0) &= \mathbf{y}_{\mathrm{v}}^{\mathrm{T}}\left(\mathbf{I}_q \otimes \left(\mathbf{X}(\mathbf{X}^{\mathrm{T}}\mathbf{X})^{-1}\mathbf{X}^{\mathrm{T}} - \mathbf{X}_0(\mathbf{X}_0^{\mathrm{T}}\mathbf{X}_0)^{-1}\mathbf{X}_0^{\mathrm{T}}\right)\right)\mathbf{y}_{\mathrm{v}}\\ &= \hat{\boldsymbol{\beta}}_{\mathrm{v}}^{\mathrm{T}}\,\mathbf{R}_{\mathrm{v}}^{\mathrm{T}}\left(\mathbf{I}_q\otimes(\mathbf{R}\,(\mathbf{X}^{\mathrm{T}}\mathbf{X})^{-1}\,\mathbf{R}^{\mathrm{T}})^{-\frac{1}{2}}\right)\left(\mathbf{I}_q\otimes(\mathbf{R}\,(\mathbf{X}^{\mathrm{T}}\mathbf{X})^{-1}\,\mathbf{R}^{\mathrm{T}})^{-\frac{1}{2}}\right)\mathbf{R}_{\mathrm{v}}\,\hat{\boldsymbol{\beta}}_{\mathrm{v}}\end{aligned}$$

Under the null hypothesis:

$$\left(\mathbf{P}\otimes\left(\mathbf{R}\,(\mathbf{X}^{\mathrm{T}}\mathbf{X})^{-1}\,\mathbf{R}^{\mathrm{T}}\right)^{-\frac{1}{2}}\right)\mathbf{R}_{\mathrm{v}}\,\hat{\beta}_{\mathrm{v}} \xrightarrow{d} N(\mathbf{0},\, \boldsymbol{\Lambda}\otimes\mathbf{I}_{p-p_0})$$

Because the limit covariance matrix is diagonal, all the $(p-p_0)\times q$ components are independent. The squared elements summed in the numerator correspond to squared univariate normals with zero mean and variance $\lambda_j$. Finally, group in $\chi_j^2(p-p_0)$ variables the components with identical eigenvalue. □

Denominators in (4) and (2) are identical to the sum of squared error terms of the simple regressions on each column of $\mathbf{Y}$. A well-known result of the OLS asymptotic properties states each of these terms converges in probability to the corresponding diagonal element of $\boldsymbol{\Sigma}$. The assumptions for such convergence are analogous to Bai's assumptions. Therefore, the denominator of the Anderson's statistic converges in probability to the trace of $\boldsymbol{\Sigma}$, that is, to $\sum_{j=1}^{q}\lambda_j$. Then, applying Lemma 2 and Slutsky's theorem, the limiting distribution of the statistic is obtained. In practice, probability tails can be computed considering only its numerator (the weights of the linear combination of chi-squares will be $\lambda_j$) or the pseudo-F ratio (weights will be $\lambda_j/\sum_{j=1}^{q}\lambda_j$).

As regards data transformations, it is straightforward to show that any transformation of $\mathbf{Y}$ that preserves the independence of the observations results in the same type of limiting distribution. Note, however, that the numerator of the test statistic, the eigenvalues of the residual covariance matrix and the resulting weighted sum of independent chi-square variables will differ with respect to the ones computed on the untransformed data.

### Supplementary Note 2: Data generation in the simplex under $H_0$ and $H_1$

Given two points (i.e. vectors of proportions) in the $q-1$ simplex, $\boldsymbol{f}_1 = (f_{11}, \ldots, f_{1q})$ and $\boldsymbol{f}_2 = (f_{21}, \ldots, f_{2q})$, a problem of interest is to find the closest point in the simplex to $\boldsymbol{f}_1$, in the direction determined by $\boldsymbol{f}_2$, obtained by adding an amount $\Delta$ to $f_{11}$. We name this point $\boldsymbol{f}_1'$. In our context, we use the Hellinger distance to assess the dissimilarity between vectors of proportions, which is not Euclidean in its natural parametrization. Therefore, to find $\boldsymbol{f}_1'$ we need to respect the geometry induced by the distance.

The geometry of the Hellinger distance in the simplex is easier to visualize if we transform the proportions to their square root, $\boldsymbol{f}_i = (\sqrt{f_{i1}}, \ldots, \sqrt{f_{iq}})$, so that the vectors are located on the surface of the upper octant of a hypersphere of radius 1. In this surface, the shortest path between $\boldsymbol{f}_1$ and $\boldsymbol{f}_2$ is the geodesic given by:

$$f'_{1j} = \left\{ \sqrt{f_{1j}} \cos(d/2) + \frac{\sqrt{f_{2j}} - \sqrt{f_{1j}} \cos(\rho/2)}{\sin(\rho/2)} \sin(d/2) \right\}^2 \qquad j \in \{1, \ldots, q\} \tag{11}$$

where $d$ is the distance traveled along the geodesic between $\boldsymbol{f}_1$ and $\boldsymbol{f}_1'$, and $\rho$ the length of the arc of the hypersphere between $\boldsymbol{f}_1$ and $\boldsymbol{f}_2$, $\rho = 2\arccos(\sum_{j=1}^{q} \sqrt{f_{1j} f_{2j}})$.

After some straightforward algebra we can obtain the expression of the components $2 \ldots q$ along the geodesic satisfying $f'_{11} = f_{11} + \Delta$:

$$f'_{1j} = f_{1j} \left( 1 - \frac{\Delta}{1 - f_{11}} \right) \qquad j \in \{2, \ldots, q\} \tag{12}$$

During data generation in the multivariate proportion scenario, we employed equation (12) to generate $\boldsymbol{p}'$ from $\boldsymbol{p}$, that is, the multivariate mean under $H_1$ given the multivariate mean under $H_0$ and different values of $\Delta$. Moreover, to actually generate random observations of vectors of proportions with certain variability, while ensuring that $E(\boldsymbol{f}_i) = \boldsymbol{p}$ (or, equivalently, $E(\boldsymbol{f}_i) = \boldsymbol{p}'$ under $H_1$), we proceeded as follows:

1. Generate a vector of $q$ step sizes, $\boldsymbol{\delta}$, using any probability distribution, so that that $E(\delta_j) = 0$ and $Var(\delta_j) = \sigma_g^2$, for $j \in \{1, \ldots, q\}$. Specifically, we obtained $\delta_j \sim N(0, \sigma_g^2)$.
2. Select randomly the order in which the steps towards the simplex vertices $\{\boldsymbol{e}_1 = (1, 0, 0, \ldots, 0)$, $\boldsymbol{e}_2 = (0, 1, 0, \ldots, 0), \ldots$, $\boldsymbol{e}_q = (0, 0, \ldots, 1)\}$ are taken.
3. If $\boldsymbol{f}_i^{(0)} = \boldsymbol{p}$, for each $j = 1, \ldots q$, $\boldsymbol{f}_i^{(j)}$ is obtained from $\boldsymbol{f}_i^{(j-1)}$ advancing from $\boldsymbol{f}_i^{(j-1)}$ towards the vertex selected in the previous step, using equation (12) with $\Delta = \delta_j$. When $j = q$ the sample generated corresponds to the last step $\boldsymbol{f}_i = \boldsymbol{f}_i^{(q)}$.

Note that $\sigma_g$ should be small enough so that $\sum_{j=1}^{q} f_{ij} = 1$, $\forall i \in \{1, \ldots, n\}$. In addition, the variances of the response variables generated, $\sigma_{jj}^2$, depend on the parameters $\sigma_g$, $q$, $L$ and the probability distribution selected to generate $\delta_j$. In practice, for a given probability distribution, we selected different values of $\sigma_g$ to ensure $\overline{\boldsymbol{\sigma}_{jj}} = 0.03$ across different values of $q$ and $L$.
